# Modeling tissue co-regulation to estimate tissue-specific contributions to disease

**DOI:** 10.1101/2022.08.25.505354

**Authors:** Tiffany Amariuta, Katherine Siewert-Rocks, Alkes L. Price

## Abstract

Integrative analyses of genome-wide association studies (GWAS) and gene expression data across diverse tissues and cell types have enabled the identification of putative disease-critical tissues. However, co-regulation of genetic effects on gene expression across tissues makes it difficult to distinguish biologically causal tissues from tagging tissues. While previous work emphasized the potential of accounting for tissue co-regulation, tissue-specific disease effects have not previously been formally modeled. Here, we introduce a new method, tissue co-regulation score regression (TCSC), that disentangles causal tissues from tagging tissues and partitions disease heritability (or covariance) into tissue-specific components. TCSC leverages gene-disease association statistics across tissues from transcriptome-wide association studies (TWAS), which implicate both causal and tagging genes and tissues. TCSC regresses TWAS chi-square statistics (or products of z-scores) on tissue co-regulation scores reflecting correlations of predicted gene expression across genes and tissues. In simulations, TCSC distinguishes causal tissues from tagging tissues while controlling type I error. We applied TCSC to GWAS summary statistics for 78 diseases and complex traits (average *N* = 302K) and gene expression prediction models for 48 GTEx tissues. TCSC identified 21 causal tissue-trait pairs at 5% FDR, including well-established findings, biologically plausible novel findings (e.g. aorta artery and glaucoma), and increased specificity of known tissue-trait associations (e.g. subcutaneous adipose, but not visceral adipose, and HDL). TCSC also identified 17 causal tissue-trait covariance pairs at 5% FDR. For the positive genetic covariance between BMI and red blood cell count, brain substantia nigra contributed positive covariance while pancreas contributed negative covariance; this suggests that genetic covariance may reflect distinct tissue-specific contributions. Overall, TCSC is a precise method for distinguishing causal tissues from tagging tissues, improving our understanding of disease and complex trait biology.

## Introduction

Most diseases are driven by tissue-specific or cell-type-specific mechanisms, thus the inference of causal disease tissues is an important goal^1^. For many polygenic diseases and complex traits, disease-associated tissues have previously been identified via the integration of genome-wide association studies (GWAS) with tissue-level functional data characterizing expression quantitative trait loci (eQTLs)^2–5^, gene expression^6–9^, or epigenetic features^10–17^. However, it is likely that most disease-associated tissues are not actually causal, due to the high correlation of eQTL effects (resp. gene expression or epigenetic features) across tissues; the correlation of eQTL effects across tissues, i.e. tissue co-regulation, can arise due to shared eQTLs or distinct eQTLs in linkage disequilibrium (LD)^2,18,19,5^. One approach to address this involves comparing eQTL-disease colocalizations across different tissues^2^; however, this approach relies on colocalizations with disease that are specific to a single tissue, and may implicate co-regulated tagging tissues that colocalize with disease. Another approach leverages multi-trait fine-mapping methods to simultaneously evaluate all tissues for colocalization with disease^5^; however, this locus-based approach does not produce genome-wide estimates and it remains the case that many (causal or tagging) tissues may colocalize with disease under this framework. To our knowledge, no previous study has formally modeled genetic co-regulation across tissues to statistically disentangle causal from tagging tissues.

Here, we introduce a new method, tissue co-regulation score regression (TCSC), that disentangles causal tissues from tagging tissues and partitions disease heritability (or genetic covariance of two diseases/traits) into tissue-specific components. TCSC leverages gene-disease association statistics across tissues from transcriptome-wide association studies (TWAS)^20,21,18^. A challenge is that TWAS association statistics include the effects of both co-regulated tissues (see above) and co-regulated genes^18,22^. To address this, TCSC regresses TWAS chi-square statistics (or products of z-scores for two diseases/traits) on tissue co-regulation scores reflecting correlations of predicted gene expression across genes and tissues. TCSC is conceptually related to gene co-regulation score regression (GCSC)^22^, a method for identifying disease-enriched gene sets that models gene co-regulation but does not model tissue co-regulation. Distinct from previous methods that analyze each tissue marginally, TCSC jointly models contributions from each tissue to identify causal tissues (analogous to the distinction in GWAS between marginal association and fine-mapping^23^). We validate TCSC using extensive simulations using real genotypes with LD, including comparisons to RTC Coloc^2^, RolyPoly^6^, LDSC-SEG^7^, and CoCoNet^9^ (reviewed in ^1,24^). We apply TCSC to 78 diseases and complex traits (average *N* = 302K) and 48 GTEx tissues^19^, showing that TCSC recapitulates known biology and identifies biologically plausible novel tissue-trait pairs (or tissue-trait covariance pairs) while attaining increased specificity relative to previous methods.

## Results

### Overview of TCSC regression

TCSC estimates the disease heritability explained by *cis*-genetic components of gene expression in each tissue when jointly modeling contributions from each tissue; a formal definition of this quantity in terms of SNP-level effects is provided in the **Methods** section. We refer to tissues with nonzero contributions as “causal” tissues (with the caveat that joint-fit effects of gene expression on disease may not reflect biological causality; see **Discussion**). TCSC assumes that gene expression-disease effect sizes are independent and identically distributed (i.i.d.) across genes and tissues (while accounting for the fact that *cis*-genetic components of gene expression are correlated across genes and tissues); violations of this model assumption are explored via simulations below. TCSC leverages the fact that TWAS *χ*^2^ statistics for each gene and tissue include both causal effects of that gene and tissue on disease and tagging effects of *co-regulated* genes and tissues. We define co-regulation based on squared correlations in *cis*-genetic expression, which can arise due to shared causal eQTLs and/or LD between causal eQTLs^18^. TCSC determines that a tissue is causal for disease if genes and tissues with high co-regulation to that tissue have higher TWAS *χ^2^* statistics than genes and tissues with low co-regulation to that tissue.

In detail, let 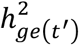 denote the disease heritability explained by the *cis*-genetic component of gene expression in tissue *t’*. The expected TWAS *χ*^2^ statistic for gene *g* and tagging tissue *t* is

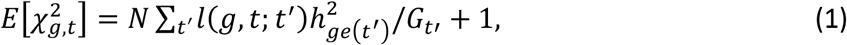

where *N* is GWAS sample size, *t’* indexes causal tissues, *l*(*g, t; t’*) are tissue co-regulation scores (defined as 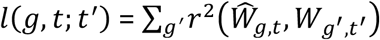, where *W*denotes the *cis*-genetic component of gene expression for a gene-tissue pair across individuals, 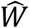 denotes the *cis*-predicted expression for a gene-tissue pair, the sum is over genes *g’* within +/- 1 Mb to gene *g*), and *G_t’_* is the number of significantly *cis*-heritable genes in tissue *t’*. A derivation of Equation (1) is provided in the **Methods** section. Equation (1) allows us to estimate 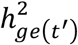 via a multiple linear regression of TWAS *χ*^2^ statistics (for each gene and tagging tissue) on tissue co-regulation scores (**Figure 1**); we note that tissue co-regulation scores reflect 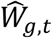 and *W_g’,t’_* but *estimated* tissue co-regulation scores reflect 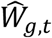 and 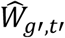, necessitating a bias correction step^22^ (**Methods**). To facilitate comparisons across diseases/traits, we primarily report the proportion of disease heritability explained by the *cis*-genetic component of gene expression in tissue 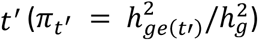, where 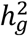 is the common variant SNP-heritability estimated by S-LDSC^13,25,26^.

**Figure 1.**
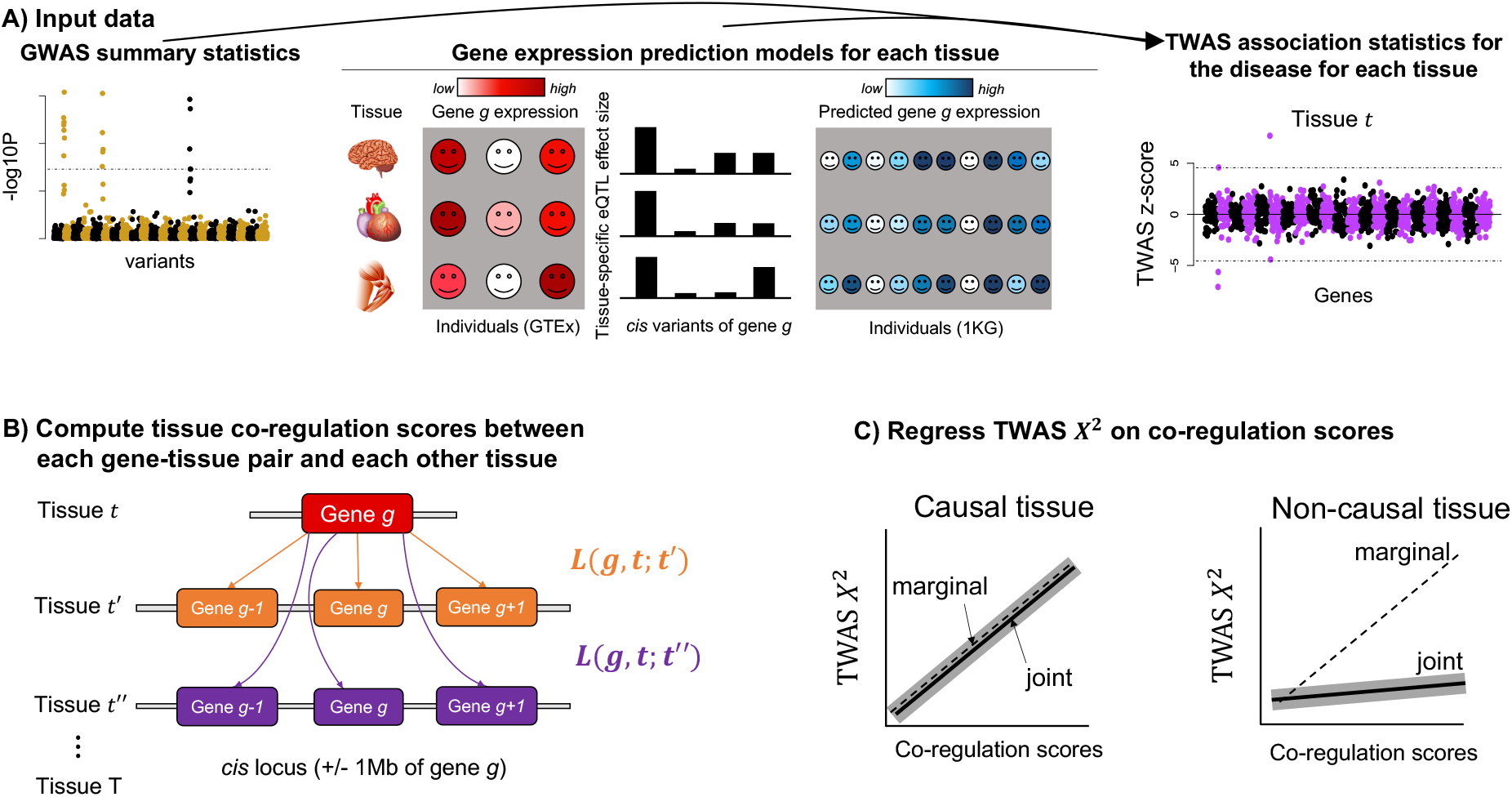
Overview of TCSC regression. (A) Input data to TCSC includes (1) GWAS summary statistics for a disease and (2) gene expression prediction models for each tissue, which are used to produce (3) TWAS summary statistics for the disease for each tissue. (B) TCSC computes tissue co-regulation scores *L*(*g, t; t’*) for each gene-tissue pair (*g, t*) with potentially causal tissues *t’*. (C) TCSC regresses TWAS chi-squares on tissue co-regulation scores to estimate tissue-specific contributions to disease. The shadow indicates the standard error of the TCSC estimate (joint models only).

TCSC can also estimate the genetic covariance between two diseases explained by *cis*-genetic components of gene expression in each tissue, using products of TWAS z-scores. In detail, let *ω_ge_*(*t’*) denote the genetic covariance explained by the *cis*-genetic component of gene expression in tissue *t’* (defined analogously to 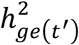; **Methods**). The expected product of TWAS z-scores in disease 1 and disease 2 for gene *g* and tagging tissue *t* is

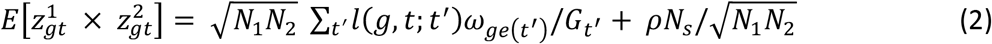

where *N*_1_ is GWAS sample size for disease 1, *N*_2_ is GWAS sample size for disease 2, *t’* indexes causal tissues, *l*(*g, t; t’*) are tissue co-regulation scores (see above), *G_t’_* is the number of significantly *cis*-heritable genes in tissue *t’* (**Methods**), *ρ* is the phenotypic correlation between disease 1 and disease 2, and *N_s_* is the number of overlapping GWAS samples between disease 1 and disease 2. Equation (2) allows us to estimate *ω_ge_*(*t’*) via a multiple linear regression of products of TWAS z-scores in disease 1 and disease 2 (for each gene and tagging tissue) on tissue co-regulation scores. We note that the last term in Equation (2) is not known a priori but is accounted for via the regression intercept, analogous to previous work^27^. To facilitate comparisons across diseases/traits, we primarily report the signed proportion of genetic covariance explained by the *cis*-genetic component of gene expression in tissue *t’* (*ζ_t’_* = *ω*_*ge*(*t’*)_/*ω_g_*), where *ω_g_* is the common variant genetic covariance estimated by cross-trait LDSC^28^.

We restrict gene expression prediction models and TWAS association statistics for each tissue to significantly *cis-*heritable genes in that tissue, defined as genes with significantly positive *cis-*heritability (2-sided *p* < 0.01; estimated using GCTA^29^) and positive adjusted-*R*^2^ in cross-validation prediction. We note that quantitative estimates of the disease heritability explained by the *cis*-genetic component of gene expression in tissue 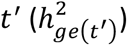 are impacted by the number of significantly *cis*-heritable genes in tissue *t’* (*G_t’_*), which may be sensitive to eQTL sample size (**Methods**). For each disease (or pair of diseases), we use a genomic block-jackknife with 200 blocks to estimate standard errors on the disease heritability (or covariance) explained by *cis*-genetic components of gene expression in each tissue, and compute 1-sided P-values for nonzero heritability (or 2-sided P-values for nonzero covariance) and false discovery rates (FDR) accordingly; we primarily report causal tissues with FDR < 5%. We use a 1-sided test for nonzero heritability because we are only interested in detecting positive tissue-specific contributions to heritability. Further details, including correcting for bias in tissue co-regulation scores arising from differences between *cis*-genetic vs. *cis*-predicted expression (analogous to GCSC^22^) and utilizing regression weights to improve power, are provided in the **Methods** section. We have publicly released open-source software implementing TCSC regression (see **Code Availability**), as well as all GWAS summary statistics, TWAS association statistics, tissue co-regulation scores, and TCSC output from this study (see **Data Availability**).

### Simulations

We performed extensive simulations to evaluate the robustness and power of TCSC, using the TWAS simulator of Mancuso et al.^30^ (see **Code Availability**). We used real genotypes from 1000 Genomes European to simulate gene expression values (for each gene and tissue) and complex trait phenotypes, and computed TWAS association statistics for each gene and tissue. In our default simulations, the number of tissues was set to 10. The gene expression sample size (in each tissue) varied from 100 to 1,500 (with the value of 300 corresponding most closely to the GTEx data^19^ used in our analyses of real diseases/traits; see below). The number of genes was set to 1,000 across chromosome 1; 100 of the 1,000 genes had nonzero (normally distributed) gene-disease effects in the causal tissue^31^. For each tissue, 500 genes were chosen to be *cis*-heritable. In the causal tissue and the three most highly genetically correlated tagging tissues, all 100 causal genes were *cis*-heritable. Each *cis*-heritable gene was assigned 5 causal *cis*-eQTLs within 50kb of the gene body, consistent with the upper range of independent eQTLs per gene detected in GTEx^19^ and other studies^32–35^. The *cis*-eQTL effect sizes for each gene were drawn from a multivariate normal distribution across tissues to achieve a specified level of co-regulation (see below), the *cis*-heritability of each gene was sampled from an exponential distribution, and neighboring co-regulated genes were assigned the same heritability to maximize gene-gene co-regulation. In each tissue, the average *cis*-heritability (across genes) was set to 0.08 (sd = 0.05, ranging from 0.01 to 0.40) in order to achieve an average estimated *cis*-heritability (*across significantly cis-heritable genes, estimated by GCTA^29^*) varying from 0.11 to 0.31 (across gene expression sample sizes), which matches empirical values from GTEx^19^. The proportions of expressed genes that were significantly *cis*-heritable and the proportion of neighboring genes with significant genetic correlation (of eQTL effects) were also matched to GTEx data^19^. The 10 tissues were split into three tissue categories to mimic biological tissue modules in GTEx^19^ (tissues 1-3, tissues 4-6, and tissues 7-10), and average *cis*-genetic correlations between tissues (averaged across genes) were set to 0.795 within the same tissue category, 0.722 between tissue categories, and 0.753 overall^36^ (**Methods**). The default GWAS sample size was set to 10,000. The 10 tissues included one causal tissue explaining 100% of trait heritability and nine non-causal tissues; 100% of trait heritability was explained by gene expression. Other parameter values were also explored, including other proportions of trait heritability explained by the causal tissue, other proportions of trait heritability not explained by gene expression, and other values of the number of causal tissues and the number of tagging tissues. Further details of the simulation framework are provided in the **Methods** section. We compared TCSC to four previously published methods: RTC Coloc^2^, RolyPoly^6^, LDSC-SEG^7^, and CoCoNet^9^. We caution that RolyPoly, LDSC-SEG, and CoCoNet do not use eQTL data, and thus the power of TCSC relative to these methods is likely to be highly sensitive to assumptions about the role of gene expression in disease architectures. We caution that the power of TCSC (and other methods) varies greatly with the choice of parameter settings (see below), thus the primary purpose of these simulations was to evaluate the robustness of TCSC relative to other methods.

We first evaluated the bias in TCSC estimates of the disease heritability explained by the *cis*-genetic component of gene expression in tissue 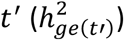, for both causal and non-causal tissues. For causal tissues, TCSC produced unbiased estimates of 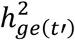 (**Figure 2A**, **Supplementary Table 1**); this implies that error in eQTL effect size estimates, which impacts TWAS statistics and co-regulation scores, does not bias TCSC estimates for causal tissues. A subtlety is that, as noted above, estimates of 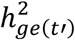 are impacted by the number of significantly *cis*-heritable genes in tissue *t’* (*G_t’_*), which may be sensitive to eQTL sample size. Estimates were conservative when setting *G_t’_* to the number of significantly *cis*-heritable genes, and unbiased when setting *G_t’_* to the number of true *cis*-heritable genes. For non-causal tissues, TCSC produced estimates of 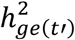, that were significantly positive when averaged across all simulations, but not large enough to substantially impact type I error (see below). In this analysis of bias in estimates of 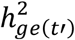, we could not include a comparison to RTC Coloc, RolyPoly, LDSC-SEG, or CoCoNet, because these methods do not provide quantitative estimates of 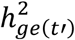.

**Figure 2.**
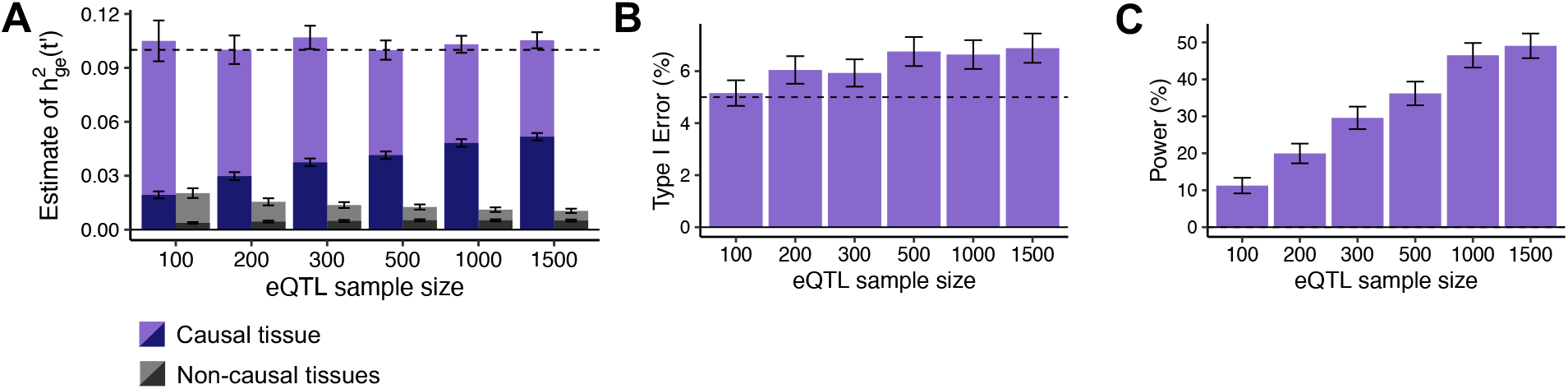
Robustness and power of TCSC regression in simulations. (A) Bias in estimates of disease heritability explained by the *cis*-genetic component of gene expression in tissue 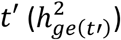 for causal (light and dark purple) and non-causal (light and dark gray) tissues, across 1,000 simulations per eQTL sample size. Light purple (resp. gray) indicates that *G_t_’* was set to the total number of true *cis*-heritable genes across tissues, dark purple (resp. gray) indicates that *G_t’_* was set to the number of significantly *cis*-heritable genes detected in each tissue. The dashed line indicates the true value of 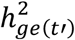 for causal tissues. (B) Percentage of estimates of 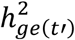 for non-causal tissues that were significantly positive at *p* < 0.05, across 1,000 simulations per eQTL sample size. The type I error for TCSC ranged from 5.2% to 6.9%. In comparison, we observed type I errors from 53%-86% for RTC Coloc, 32%-33% for LDSC-SEG, 11%-12% for RolyPoly, and 32%-38% for CoCoNet (**Supplementary Figure 1**, **Supplementary Table 2**). (C) Percentage of estimates of 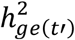 for causal tissues that were significantly positive at *p* < 0.05, across 1,000 simulations per eQTL sample size. Error bars denote 95% confidence intervals. Numerical results are reported in **Supplementary Table 1**.

We next evaluated the type I error of TCSC for non-causal tissues. The type I error of TCSC was approximately well-calibrated, ranging from 5.2% to 6.9% across eQTL sample sizes at a significance threshold of *p* = 0.05 (**Figure 2B**, **Supplementary Table 1**). In comparison, we observed type I errors from 53%-86% for RTC Coloc, 32%-33% for LDSC-SEG, 11%-12% for RolyPoly, and 32%-38% for CoCoNet, substantially greater than the type I error of TCSC (**Supplementary Figure 1**, **Supplementary Table 2**).

We next evaluated the power of TCSC for causal tissues. We determined that TCSC was moderately well-powered to detect causal tissues, with power ranging from 11%-49% across eQTL sample sizes at a nominal significance threshold of *p* < 0.05 (**Figure 2C**) (and 1%-18% at a stringent significance threshold of *p* < 0.004, corresponding to 5% per-trait FDR across tissues in these simulations; **Supplementary Table 1**). As noted above, the power of TCSC varies greatly with the choice of parameter settings (see below), thus the power of TCSC in real-world settings is best evaluated using real trait analysis. As expected, power increased at larger eQTL sample sizes, due to lower standard errors on point estimates of 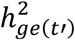 (**Figure 2A**). We also evaluated the power of RTC Coloc, RolyPoly, LDSC-SEG, and CoCoNet. For the only other method with type I error less than 15% (RolyPoly), power ranged from 14%-17% across eQTL sample sizes, substantially lower than TCSC (**Supplementary Figure 1**, **Supplementary Table 2**). We also used ROC curves to assess the relationship between the sensitivity (power) and specificity (one minus the false positive rate) of all 5 methods across 1,000 uniformly spaced p-value thresholds. TCSC attained the largest AUC (0.78, vs. 0.54-0.59 for other methods) (**Supplementary Figure 1**).

We similarly evaluated the robustness and power of TCSC when estimating tissue-specific contributions to the genetic covariance between two diseases/traits; we did not compare TCSC to RTC Coloc, RolyPoly, LDSC-SEG, and CoCoNet, which are not applicable to cross-trait analysis. We employed the same simulation framework described above and set the genetic correlation of the two simulated traits to 0.5. We first evaluated the bias in TCSC estimates of the genetic covariance explained by the *cis*-genetic component of gene expression in tissue *t’* (*ω*_*ge*(*t’*)_), for both causal and non-causal tissues (**Figure 3A**, **Supplementary Table 3**). For causal tissues, TCSC produced unbiased estimates of *ω*_*ge*(*t’*)_ (conservative estimates when setting *G_t’_* to the number of significantly *cis*-heritable genes, rather than the number of true *cis*-heritable genes), analogous to single-trait simulations. For non-causal tissues, TCSC again produced estimates of *ω*_*ge*(*t’*)_ that were significantly positive when averaged across all simulations, but not large enough to substantially impact type I error. We next evaluated the type I error of cross-trait TCSC for non-causal tissues. TCSC was well-calibrated with type I error ranging from 5.4%-6.7% at *p* < 0.05 (**Figure 3B**). Finally, we evaluated the power of cross-trait TCSC for causal tissues. We determined that cross-trait TCSC was modestly powered at realistic eQTL sample sizes, with power ranging from 8%-27% across eQTL sample sizes at *p* < 0.05 (**Figure 3C**) (and 1-6% power at *p* < 0.004 corresponding to 5% per-trait FDR across tissues in these simulations; **Supplementary Table 3**); as noted above, the power of TCSC varies greatly with the choice of parameter settings (see below). In ROC curve analysis, TCSC attained an AUC of 0.67 (**Supplementary Figure 1**).

**Figure 3.**
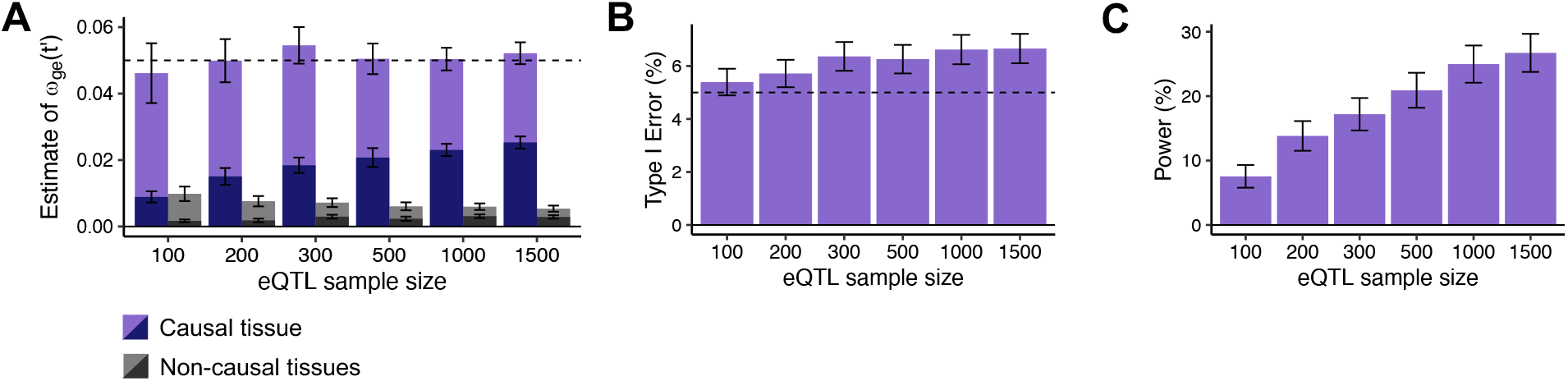
Robustness and power of cross-trait TCSC in simulations. (A) Bias in estimates of genetic covariance explained by the *cis*-genetic component of gene expression in tissue *t’* (*ω*_*ge*(*t’*)_) for causal (light and dark purple) and non-causal (light and dark gray) tissues, across 1,000 simulations per eQTL sample size. Light purple (resp. gray) indicates that *G_t’_* was set to the total number of true *cis*-heritable genes across tissues, dark purple (resp. gray) indicates that *G_t’_* was set to the number of significantly *cis*-heritable genes detected in each tissue. The dashed line indicates the true value of *ω*_*ge*(*t’*)_ for causal tissues. (B) Percentage of estimates of *ω*_*ge*(*t’*)_ for non-causal tissues that were significantly positive at *p* < 0.05, across 1,000 simulations per eQTL sample size. (C) Percentage of estimates of *ω*_*ge*(*t’*)_ for causal tissues that were significantly positive at *p* < 0.05, across 1,000 simulations per eQTL sample size. Error bars denote 95% confidence intervals. Numerical results are reported in **Supplementary Table 3**.

We performed 12 secondary analyses. First, we varied the eQTL sample size across tissues. Specifically, we set the eQTL sample size of the causal tissue to 300 individuals and the eQTL sample sizes of the non-causal tissues to range between 100 and 1,500 individuals. We observed inflated type I error for non-causal tissues (particularly those with larger eQTL sample sizes), implying that large variations in eQTL sample sizes may compromise type I error (**Supplementary Figure 2**). Second, we evaluated the robustness of TCSC when varying the number of expressed genes in the causal tissue under four scenarios: (i) only the 500 *cis*-heritable genes are expressed in the causal tissue, (ii) only 375 *cis*-heritable genes (including all 100 causal genes) are expressed in the causal tissue, (iii) only 225 *cis*-heritable genes (including all 100 causal genes) are expressed in the causal tissue, and (iv) only the 100 causal genes are expressed in the causal tissue. We determined that type I error remained approximately well-calibrated in all scenarios, and that power was dramatically improved and bias for non-causal tissues decreased as the number of tagging genes in the causal tissue decreased (**Supplementary Figures 3-4**); for causal tissues, estimates of 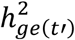 were upward biased when setting *G_t’_* to the number of true *cis*-heritable genes and unbiased when setting *G_t’_* to the number of significantly *cis*-heritable genes across tissues. Third, we varied the true values of 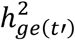 (or *ω*_*ge*(*t’*)_) for causal tissues. We determined that patterns of bias, type I error, and power were generally robust across different parameter values, although the smallest values resulted in lower power and greater bias for non-causal tissues (**Supplementary Figures 5-6**). Fourth, we varied the number of causal tissues, considering 1, 2, or 3 causal tissues. We observed that the power of TCSC decreased with multiple causal tissues but did not differ greatly between 2 and 3 causal tissues (**Supplementary Figures 7-8**); for causal tissues, estimates of 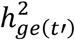 were upward biased when setting *G_t’_*, to the number of true *cis*-heritable genes. Fifth, we varied the number of non-causal tissues from 0 to 9. For causal tissues, TCSC estimates were upward biased with fewer tagging tissues but unbiased with more tagging tissues (**Supplementary Figures 9-10**). TCSC type I error and power were generally higher with fewer tagging tissues; this finding does not compromise our real trait analysis, which involve a large number of tissues. Sixth, we modified TCSC to not correct for bias in tissue co-regulation scores arising from differences between *cis*-genetic and *cis*-predicted expression. We determined that removal of bias correction resulted in conservative bias in estimates for causal tissues, increased type I error, and similar power (**Supplementary Figures 11-12**). Seventh, we modified TCSC to apply bias correction to the calculation of all correlations of *cis*-predicted expression contributing to co-regulation scores rather than only those involving the same gene and tissue, which resulted in a decrease in power, anti-conservative bias in estimates for causal tissues, and similar type I error rate (**Supplementary Figures 13-14**). Eighth, we modified TCSC to use bias-corrected co-regulation scores in the calculation of regression weights, which resulted in similar performance to the default setting (**Supplementary Figures 15-16**). We note that regression weights pertain to maximizing signal to noise and not avoiding bias in estimates of 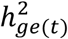; we continue to not perform bias correction when calculating regression weights, consistent with GCSC^22^. Ninth, we violated the model assumption that gene-disease effects are independent and identically distributed (i.i.d.) across tissues by including a second causal tissue whose gene-disease effects correlate with varying degree to the gene-disease effects of the original causal tissue (**Supplementary Figures 17-18**). We determined that while this increases noise to TCSC estimates, the estimates are generally unbiased and TCSC is able to powerfully identify the causal tissue, similar to the addition of a causal tissue where there are no shared gene-disease effects (see **Supplementary Figures 7-8**). Tenth, we violated the i.i.d. model assumption by duplicating the causal tissue. We determined that TCSC performs well, (e.g. frequently identifies both tissues as causal and estimates 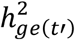 for both tissues without bias) despite the violation of model assumption (**Supplementary Figures 19-20**), similar to the previous analysis. Eleventh, we evaluated the robustness of TCSC in the presence of disease heritability that is not mediated via gene expression. We observed that all areas of TCSC performance are affected, with slightly increased type I error rates, decreased power in the case of larger non-mediate heritability, and upward bias in estimates of 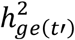 for causal tissues (**Supplementary Figures 21-22**). Finally, we evaluated the robustness of TCSC to variation in the window size used to identify co-regulated genes in the calculation of co-regulation scores and determined that TCSC performance was robust and type I error decreased with larger window sizes (**Supplementary Figures 23-24**). Further details of these secondary analyses are provided in the **Supplementary Note**.

### Identifying tissue-specific contributions to 78 diseases and complex traits

We applied TCSC to publicly available GWAS summary statistics for 78 diseases and complex traits (average *N* = 302K; **Supplementary Table 4**) and gene expression data for 48 GTEx tissues^19^ (**Table 1**) (see **Data Availability**). The 78 diseases/traits (which include 33 diseases/traits from UK Biobank^37^) were selected to have z-score > 6 for nonzero SNP-heritability (as in previous studies^13,25,38^), with no pair of diseases having squared genetic correlation > 0.1^28^ and substantial sample overlap (**Methods**). The 48 GTEx tissues were aggregated into 39 *meta-tissues* (average N = 266, range: *N* = 101-320 individuals, 23 meta-tissues with *N* = 320) in order to reduce variation in eQTL sample size across tissues (**Table 1** and **Methods**); below, we refer to these as “tissues” for simplicity. We constructed gene expression prediction models for an average of 3,993 significantly *cis*-heritable protein-coding genes (as defined above) in each tissue. We primarily report the proportion of disease heritability explained by the *cis*-genetic component of gene expression in tissue 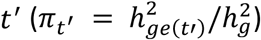, as well as its statistical significance (using per-trait FDR). We employ a per-trait FDR (as in ref.^39,40^) rather than a global FDR (as in ref.^7^), because power is likely to vary across traits and there are a sufficiently large number of independent quantities estimated per trait (*π_t’_* jointly estimated across 39 tissues); a global FDR is more appropriate when there are far fewer independent quantities estimated per trait, e.g. due to non-independent, marginal tissue associations in ref.^7^.

**Table 1.** GTEx meta-tissues and constituent tissues analyzed. For each meta-tissue we list the constituent tissue(s) and total sample size. Daggers denote meta-tissues with more than one constituent tissue; for these meta-tissues, each constituent tissue has equal sample size up to rounding error (an exception is the transverse intestine meta-tissue, which includes 176 transverse colon samples and all 144 small intestine samples).

| Meta-tissue | Constituent Tissue(s) | Sample Size |
| --- | --- | --- |
| Adipose Subcutaneous | Adipose Subcutaneous | 320 |
| Adipose Visceral Omentum | Adipose Visceral Omentum | 320 |
| Adrenal Gland | Adrenal Gland | 200 |
| Aorta Artery | Aorta Artery | 320 |
| † Brain Basal Ganglia | Putamen, Caudate, Nucleus Accumbens | 320 |
| † Brain Cereb. | Cerebellum, Cerebellar Hemisphere | 320 |
| † Brain Cortex | Frontal, Anterior, Cingulate | 320 |
| † Brain Limbic | Amygdala, Hippocampus, Hypothalamus | 320 |
| Brain Spinal Cord | Brain Spinal Cord | 115 |
| Brain Substantia Nigra | Brain Substantia Nigra | 101 |
| Breast Mammary Gland | Breast Mammary Gland | 320 |
| Coronary Artery | Coronary Artery | 180 |
| Cultured Fibroblasts | Cultured Fibroblasts | 320 |
| Esophagus Mucosa | Esophagus Mucosa | 320 |
| Esophagus Muscularis | Esophagus Muscularis | 320 |
| Heart Atrial Appendage | Heart Atrial Appendage | 320 |
| Heart Left Ventricle | Heart Left Ventricle | 320 |
| LCLs | LCLs | 116 |
| Liver | Liver | 183 |
| Lung | Lung | 320 |
| Minor Salivary Gland | Minor Salivary Gland | 118 |
| Muscle Skeletal | Muscle Skeletal | 320 |
| Ovary | Ovary | 140 |
| Pancreas | Pancreas | 252 |
| Pituitary | Pituitary | 220 |
| Prostate | Prostate | 186 |
| Skin (sun exposed) | Skin (sun exposed) | 320 |
| Skin (sun unexposed) | Skin (sun unexposed) | 320 |
| † Sigmoid Intestine | Sigmoid Colon, Gastroesophageal Junction | 320 |
| Spleen | Spleen | 185 |
| Stomach | Stomach | 269 |
| Tibial Artery | Tibial Artery | 320 |
| Tibial Nerve | Tibial Nerve | 320 |
| Testis | Testis | 277 |
| Thyroid | Thyroid | 320 |
| † Transverse Intestine | Transverse Colon, Small Intestine | 320 |
| Uterus | Uterus | 108 |
| Vagina | Vagina | 122 |
| Whole Blood | Whole Blood | 320 |

TCSC identified 21 causal tissue-trait pairs with significantly positive contributions to disease/trait heritability at 5% FDR, spanning 7 distinct tissues and 17 distinct diseases/traits (**Figure 4**, **Supplementary Table 5, Supplementary Figure 25**). Many of the significant findings recapitulated known biology, including associations of whole blood with blood cell traits such as white blood cell count (*π_t’_* = 0.21, s.e. = 0.064, *P* = *5.7 ×* 10^-4^) and liver with lipid traits such as LDL (*π_t’_* = 0.20, s.e. = 0.050, *P* = 2.9 × 10^-5^). We obtained independent GWAS summary statistics for 10 traits implicated in 13 significant tissue-trait pairs (**Supplementary Table 4**) and confirmed the same direction of effect for 13 of 13 tissue-trait pairs (including FDR < 5% for 7 of 13 tissue-trait pairs, FDR < 10% for 9 of 13 tissue-trait pairs) (**Supplementary Table 5**); however, FDR < 5% results are expected to include a small number of false positives, and our association of whole blood with major depressive disorder (FDR < 5% in primary analysis; same direction, FR = 84% in independent GWAS data) may be one of these. In our primary analysis, TCSC also identified 5 suggestive tissue-trait pairs with 5% < FDR < 10% (**Figure 4, Supplementary Table 6**).

**Figure 4.**
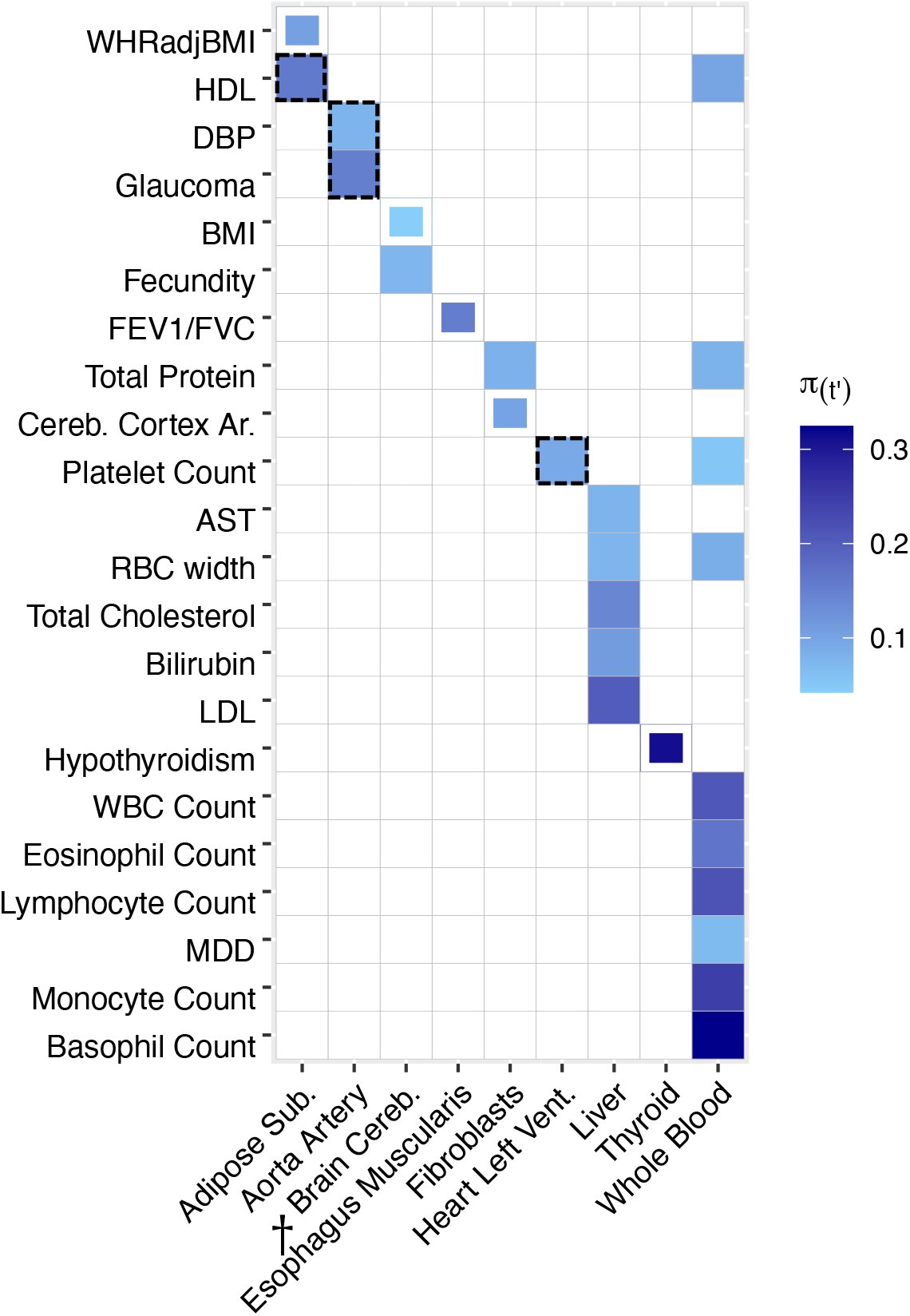
TCSC estimates tissue-specific contributions to disease and complex trait heritability. We report estimates of the proportion of disease heritability explained by the *cis*-genetic component of gene expression in tissue *t’* (*π_t’_*). We report tissue-trait pairs with FDR of 10% or lower, where full boxes denote FDR of 5% or lower and partial boxes denote FDR between 5% and 10%. Dashed boxes denote results that are highlighted in the main text. Tissues are ordered alphabetically. Daggers denote meta-tissues with more than one constituent tissue. Diseases/traits are ordered with respect to causal tissues. Numerical results are reported in **Supplementary Tables 5 and Supplementary Table 6** (for all traits). WHRadjBMI: waist-hip-ratio adjusted for body mass index. HDL: high-density lipoprotein. DBP: diastolic blood pressure. BMI: body mass index. FEV1/FVC: forced expiratory volume in one second divided by forced vital capacity. Cereb. Cortex Ar.: cerebral cortex surface area. AST: aspartate aminotransferase. LDL: low-density lipoprotein. WBC Count: white blood cell count. MDD: major depressive disorder.

TCSC also identified several biologically plausible findings not previously reported in the genetics literature. First, aorta artery was associated with glaucoma (*π_t’_* = 0.15, s.e. = 0.051, *P* = 1.3 × 10^-3^). TCSC also identified aorta artery as a causal tissue for diastolic blood pressure (DBP) (*π_t’_* = 0.078, s.e. = 0.024, *P* = 5.1 *×* 10^-4^), which is consistent with DBP measuring the pressure exerted on the aorta when the heart is relaxed^41^. High blood pressure is a known risk factor for glaucoma^42–46^, explaining the role of aorta artery in genetic susceptibility to glaucoma. Second, TCSC identified heart left ventricle (in addition to whole blood) as a causal tissue for platelet count (*π_t’_* = 0.091, s.e. = 0.031, *P* = 1.7 × 10^-3^), consistent with the role of platelets in the formation of blood clots in cardiovascular disease^47–50^. In cardiovascular disease, platelets are recruited to damaged heart vessels after cholesterol plaques rupture, resulting in blood clots due to the secretion of coagulating molecules^51^; antiplatelet drugs have been successful at reducing adverse cardiovascular outcomes^52^. Moreover, the left ventricle serves as a muscle to pump blood throughout the body^53^, likely modulating platelet counts and other blood cell counts, creating detectable changes in serum from which platelet counts are measured. Other significant findings are discussed in the **Supplementary Note**, and numerical results for all tissues and diseases/traits analyzed are reported in **Supplementary Table 6**.

TCSC also increased the specificity of known tissue-trait associations. For high density lipoprotein (HDL), previous studies reported that deletion of a cholesterol transporter gene in adipose tissue reduces HDL levels, consistent with the fact that adipose tissues are storage sites of cholesterol and express genes involved in cholesterol transport and HDL lipidation^54,55^. While there are three adipose tissues represented in the GTEx data that we analyzed (subcutaneous, visceral, and breast tissue), TCSC specifically identified subcutaneous adipose (*π_t’_* = 0.16, s.e. = 0.054, *P* = 1.5 × 10^-3^; **Figure 4**), but not visceral adipose or breast tissue (*P* > 0.05; **Supplementary Table 6**), as a causal tissue for HDL. Previous studies have established that levels of adiponectin, a hormone released by adipose tissue to regulate insulin, are significantly positively correlated with HDL^56–58^ and more recently, a study has reported that adiponectin levels are associated specifically with subcutaneous adipose tissue and not visceral adipose tissue^59^; thus, the specific role of subcutaneous adipose tissue in HDL may be due to a causal mechanism related to adiponectin. We note that TCSC did not identify liver as a causal tissue for HDL (FDR > 5%), which may be due to limited power in liver due to smaller eQTL sample size. For waist-hip ratio adjusted for BMI (WHRadjBMI), previous studies reported colocalization of WHRadjBMI GWAS variants with *cis*-eQTLs in subcutaneous adipose, visceral adipose, liver, and whole blood^60^, consistent with WHRadjBMI measuring adiposity in the intraabdominal space which is likely regulated by metabolically active tissues^61^. TCSC specifically identified subcutaneous adipose as a suggestive finding (*π_t’_* = 0.10, s.e. = 0.037, *P* = 2.4 × 10^-3^, 5% < FDR < 10%; **Figure 4**), but not visceral adipose, breast, liver, or whole blood (*P* > 0.05; **Supplementary Table 6**), as a causal tissue for WHRadjBMI. The causal mechanism may involve adiponectin secreted from subcutaneous adipose tissue, which is negatively correlated with WHRadjBMI^62^. We note that the *P* value distributions across traits are similar for subcutaneous adipose (median *P* = 0.42) and visceral adipose (median *P* = 0.56) and are comparable to the other 37 analyzed (median *P* = 0.20 – 0.84, **Supplementary Table 7**). For BMI, previous studies have broadly implicated the central nervous system, but did not reveal more precise contributions^63,13,64,65,7,66^. TCSC specifically identified brain cereb. as a suggestive finding (*π_t’_* = 0.042, s.e. = 0.015, *P* = 2.6 × 10^-3^, 5% < FDR < 10%), but not brain cortex or brain limbic (P > 0.05; **Supplementary Table 6**), as a causal tissue for BMI. This finding is consistent with a known role for brain cerebellum in biological processes related to obesity including endocrine homeostasis^67^ and feeding control^68^; recently, a multi-omics approach has revealed cerebellar activation in mice upon feeding^69^.

We performed a secondary analysis in which we removed tissues with eQTL sample size less than 320 individuals, as these tissues may often be underpowered (**Figure 2C**). Results are reported in **Supplementary Figure 26** and **Supplementary Table 8.** The number of causal tissue-trait pairs with significantly positive contributions to disease/trait heritability (at 5% FDR) increased from 21 to 23, likely due to a decrease in multiple hypothesis testing burden from removing underpowered tissues. The 23 significant tissue-trait pairs reflect a gain of 8 newly significant tissue-trait pairs (and a loss of 6 formerly significant tissue-trait pairs, of which 5 were lost because the tissue was removed), but estimates of *π_t’_* for each significant tissue-trait pair were not statistically different from our primary analysis (**Supplementary Table 9**). Notably, among the newly significant tissue-trait pairs, whole blood was associated with hypothyroidism (*π_t’_* = 0.100, s.e. = 0.032, *P* = 8.9 × 10^-4^); we note that thyroid had a quantitatively large but only nominally significant association *(π_t_!* = 0.452, s.e. = 0.225, *P* = 0.02, FDR = 26%). Esophagus muscularis (rather than lung tissue) was associated with the lung trait FEV1/FVC^70^ *(π_t’_* = 0.167, s.e. = 0.056, *P* = *1.4 ×* 10^-3^). This result may be explained by the fact that smooth muscle in the lung is known to affect FEV1/FVC and influence pulmonary disease pathopysiology^71^, and this unobserved causal tissue is likely highly co-regulated with the smooth muscle of the esophagus, which is indeed the site from which the GTEx study sampled the esophagus muscularis tissue^19^. Other newly significant findings are discussed in the **Supplementary Note**, and numerical results for all tissues and diseases/traits are reported in **Supplementary Table 8**.

We also performed a brain-specific analysis in which we applied TCSC to 41 brain traits (average *N* = 226K, **Supplementary Table 10**) while restricting to 13 individual GTEx brain tissues (**Supplementary Table 11**), analogous to previous work^7^. The 41 brain traits reflect a less stringent squared genetic correlation threshold of 0.25; we relaxed our threshold so that we would have a substantial number of brain traits to analyze, as many would were excluded under the original threshold of 0.1. The 13 GTEx brain tissues were analyzed without merging tissues into meta-tissues, and irrespective of eQTL sample size (range: *N* = 101-189 individuals); we expected power to be limited due to the eQTL small sample sizes and substantial co-regulation among individual brain tissues. TCSC identified 8 brain tissue-brain trait pairs at 5% FDR (**Supplementary Figure 27, Supplementary Table 12**). For ADHD, TCSC identified brain hippocampus as a causal tissue (*π_t’_* = 0.127, s.e. = 0.045, *P* = 2.5 × 10^-3^), consistent with the correlation between hippocampal volume and ADHD diagnosis in children^72^. A recent ADHD GWAS identified a locus implicating the *FOXP2* gene^73^, which has been reported to regulate dopamine secretion in mice^74^; hippocampal activation results in the firing of dopamine neurons^75^. For BMI, TCSC identified brain amygdala (*π_t’_* = 0.054, s.e. = 0.023, *P* = 8.3 × 10^-3^) and brain cerebellum (*π_t’_* = 0.039, s.e. = 0.016, *P* = 7.0 × 10^-3^) as causal tissues, consistent with previous work linking the amygdala to obesity and dietary self-control^76^, although no previous study has implicated the amygdala in genetic regulation of BMI. As for brain cerebellum, previous research has implicated the cerebellar function in dietary behavior, rather than strictly regulation motor control function^67–69^. We note that the brain-specific analysis is expected to have greater power to identify tissue-trait pairs than the analysis of **Figure 4** due to the smaller number of total tissues in the model (as simulations show higher power for TCSC when there are fewer tagging tissues; **Supplementary Figure 9**). Other significant findings are discussed in the **Supplementary Note**, and numerical results for all brain tissues and brain traits analyzed are reported in **Supplementary Table 12**.

### Comparisons of TCSC to other methods

We compared TCSC to two previous methods, RTC Coloc^2^ and LDSC-SEG^7^, that identify disease-critical tissues using gene expression data. RTC Coloc identifies disease-critical tissues based on tissue specificity of eQTL-GWAS colocalizations. LDSC-SEG identifies disease-critical tissues based on heritability enrichment of specifically expressed genes. We included RTC Coloc in these comparisons because it is the only other method that analyzes eQTL data and included LDSC-SEG because we believe it is the most widely used method. We note that RTC Coloc and LDSC-SEG analyze each tissue marginally, whereas TCSC jointly models contributions from each tissue to identify causal tissues (analogous to the distinction in GWAS between marginal association and fine-mapping^23^). Thus, we hypothesized that RTC Coloc and LDSC-SEG may output multiple highly statistically significant associated tissues for a given trait, whereas TCSC may output a single causal tissue with weaker statistical evidence of causality. To assess whether TCSC indeed attains higher specificity, we evaluated the results of each method both for causal tissues identified by TCSC and for the most strongly co-regulated tagging tissue (based on Spearman *ρ* for estimated eQTL effect sizes, averaged across genes, from ref.^19^). Our primary analyses focused on 7 traits with at least one tissue-trait association for each of the three methods (**Methods**).

Results for the 7 traits are reported in **Figure 5** and **Supplementary Table 13**; results for all 17 diseases/traits with causal tissue-trait associations identified by TCSC (**Figure 4**) are reported in **Supplementary Figure 28** and **Supplementary Table 14**, and complete results for all diseases/traits and tissues included in these comparisons are reported in **Supplementary Table 15**. We reached three main conclusions. First, for a given disease/trait, RTC Coloc typically implicates a broad set of tissues (not just strongly co-regulated tissues) (**Figure 5A**); for example, for WBC count, RTC Coloc implicated 8 of 10 tissues in **Figure 5**. This is consistent with our simulations, in which RTC Coloc suffered a high type I error rate and had a substantially lower AUC than TCSC (**Supplementary Figure 1**). Second, for a given disease/trait, LDSC-SEG typically implicates a small set of strongly co-regulated tissues (**Figure 5B**); for WBC count, LDSC-SEG implicated 3 of 8 tissues in **Figure 5**, consisting of whole blood and spleen (which are strongly co-regulated) plus breast tissue. This is consistent with our simulations, in which LDSC-SEG suffered a substantial type I error rate and had a substantially lower AUC than TCSC (**Supplementary Figure 1**). Third, for a given disease/trait, TCSC typically implicates one causal tissue (**Figure 5C**); for WBC count, TCSC implicated only whole blood as a causal tissue, with even the most strongly co-regulated tagging tissue reported as non-significant. This is consistent with our simulations, in which TCSC attained moderate power to identify causal tissues with approximately well-calibrated type I error. However, we caution that the higher specificity of TCSC in identifying unique causal tissues may be accompanied by incomplete power to identify secondary causal tissues; accordingly, we observed less significant (lower -log_10_P-value and lower -log_10_FDR) results for causal tissues in **Figure 5C** than in **Figure 5A** and **Figure 5B**) (**Supplementary Table 14**). We also observed similar patterns when comparing TCSC to RTC Coloc and LDSC-SEG in the brain-specific analysis of **Supplementary Figure 27** (**Supplementary Figure 29**, **Supplementary Table 16, Supplementary Note**). Based on simulations, we expect that RTC Coloc and LDSC-SEG both attain higher power at the cost of higher false positives.

**Figure 5.**
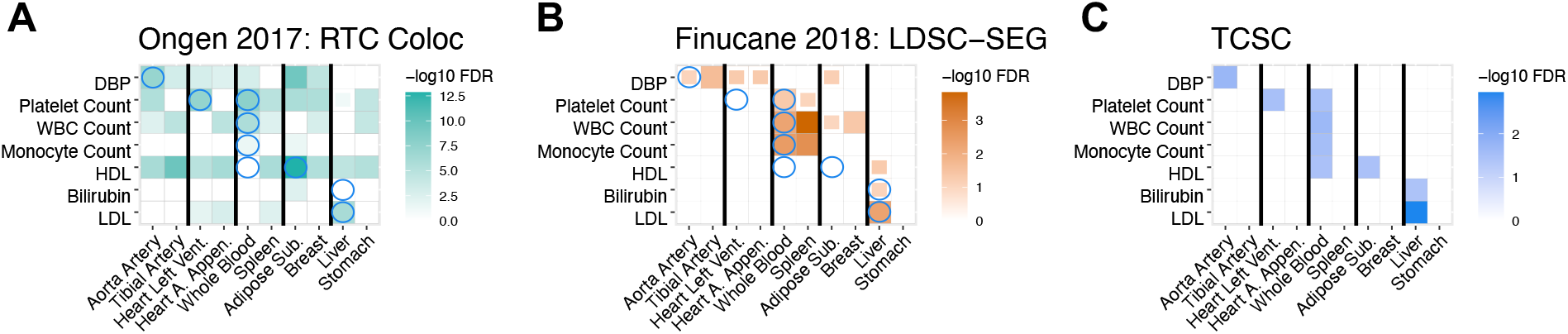
Comparison of disease-critical tissues identified by RTC Coloc, LDSC-SEG and TCSC. We report -log_10_FDR values for (A) RTC Coloc, (B) LDSC-SEG, (C) TCSC, across 7 traits with at least one significantly associated tissues (at FDR 5%) for each of the three methods and 10 tissues consisting of the causal tissues identified by TCSC and the most strongly co-regulated tagging tissues, ordered consecutively. We report tissue-trait pairs with FDR of 10% or lower, where full boxes denote FDR of 5% or lower and partial boxes denote FDR between 5% and 10%. Blue circles in panels (A) and (B) denote the causal tissue-trait pairs identified by TCSC. Numerical results are reported in **Supplementary Table 13**.

### Identifying tissue-specific contributions to the genetic covariance between two diseases/traits

We applied cross-trait TCSC to 262 pairs of disease/traits (**Supplementary Table 17**) and gene expression data for 48 GTEx tissues^19^ (**Table 11**) (see **Data Availability**). Of 3,003 pairs of the 78 disease/traits analyzed above, the 262 pairs of diseases/traits were selected based on significantly nonzero genetic correlation (*p* < 0.05 / 3,003; see **Methods**). The 48 GTEx tissues were aggregated into 39 meta-tissues, as before (**Table 1** and **Methods**). We primarily report the signed proportion of genetic covariance explained by the *cis*-genetic component of gene expression in tissue *t’* (*ζ_t’_* = *ω*_*ge*(*t’*)_ /*ω_g_*), as well as its statistical significance (using per-trait FDR). We note that the direction of effect of tissue-specific contributions to the genetic covariance between two traits may be in the opposite direction of the global covariance between two traits, analogous to how local contributions to genome-wide genetic correlation may be in the opposite direction of the genome-wide genetic correlation^77–80^.

TCSC identified 17 causal tissue-trait covariance pairs with significant contributions to trait covariance at 5% FDR, spanning 12 distinct tissues and 13 distinct trait pairs (**Figure 6A**, **Supplementary Table 18**). For 16 of the 17 causal tissue-trait covariance pairs, the causal tissue was non-significant for *both* constituent traits in the single-trait analysis of **Supplementary Table 8**. Findings that recapitulated known biology included both examples involving a tissue-trait pair that was significant in the single-trait analysis (marked by an underline in **Figure 6A**, **Figure 4**) and examples in which both tissue-trait pairs were non-significant in the single-trait analysis (**Supplementary Table 6**). Consistent with the significant contribution of liver to LDL heritability in the single-trait analysis, TCSC identified a suggestive positive contribution of liver to the genetic covariance of LDL and total cholesterol (*ζ_t’_* = 0.090, s.e. = 0.029, *P* = 1.0 × 10^-3^, 5% < FDR < 10%), and consistent with the positive contributions of whole blood to eosinophil count heritability and to platelet count heritability in the single-trait analysis, TCSC identified a significant positive contribution of whole blood to the genetic covariance of eosinophil count and platelet count (*ζ_t’_* = 0.30, s.e. = 0.10, *P* = 2.3 × 10^-3^). TCSC also identified 15 suggestive tissue-trait covariance pairs with 5% < FDR < 10% (**Figure 6A**, **Supplementary Table 19**).

**Figure 6.**
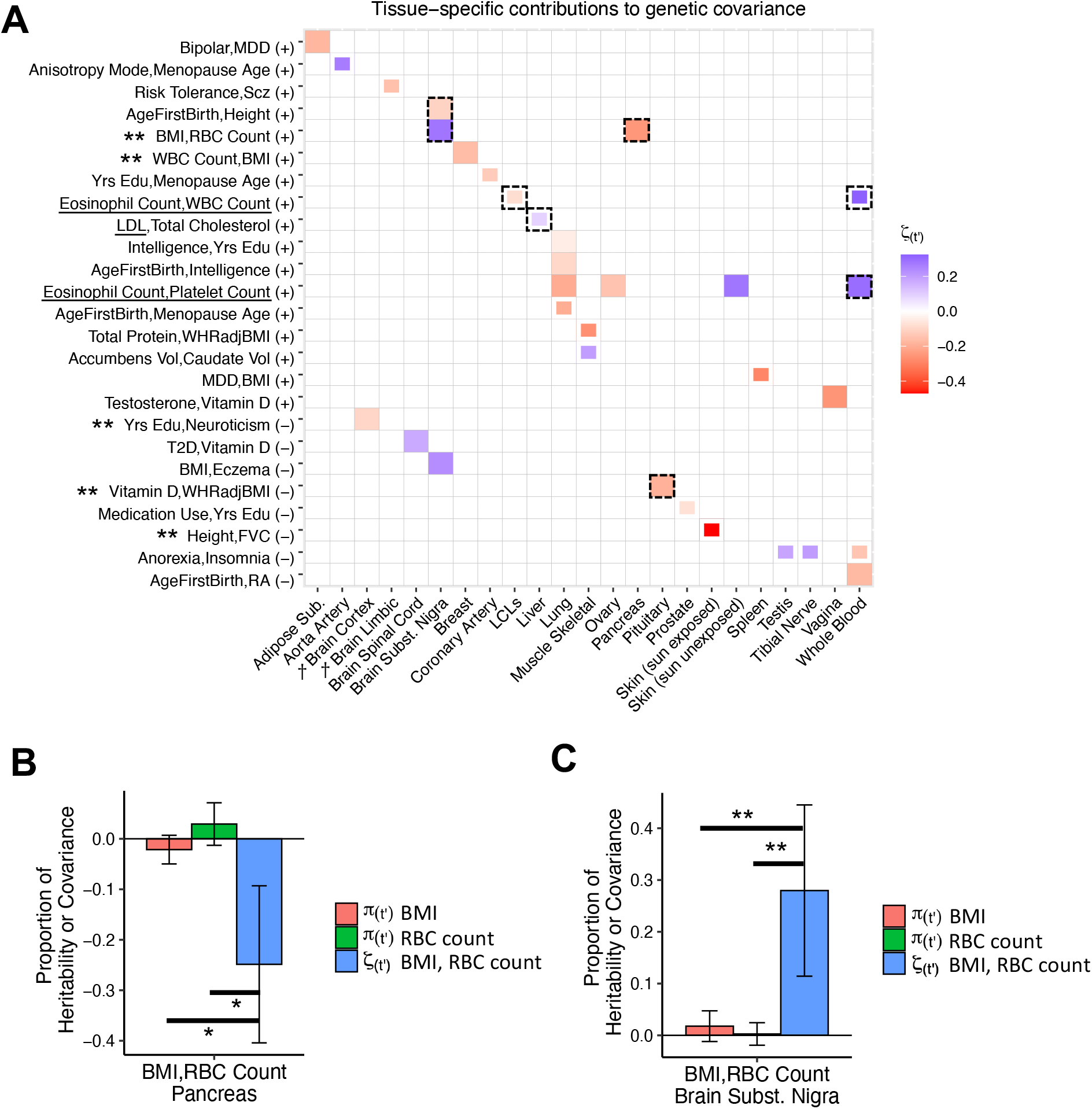
Cross-trait TCSC estimates tissue-specific contributions to the genetic covariance of two diseases/traits. (A) We report estimates of the signed proportion of genetic covariance explained by the *cis*-genetic component of gene expression in tissue *t’* (*ζ_t’_*). We report tissuetrait covariance pairs with FDR of 10% or lower, where full boxes denote FDR of 5% or lower and partial boxes denote FDR between 5% and 10%. Dashed boxes denote results that are highlighted in the main text. Tissues are ordered alphabetically. Daggers denote meta-tissues with more than one constituent tissue. Trait pairs are ordered by positive (+) or negative (-) genetic covariance, and further ordered with respect to causal tissues. Underlined traits are those for which TCSC identified a causal tissue in **Figure 4**: for eosinophil count, WBC count, and platelet count the causal tissue was whole blood, and for LDL the causal tissue was liver. Double asterisks denote trait pairs for which the differences between the tissue-specific contribution to covariance and the tissue-specific contributions to heritability were significant for both constituent traits and the tissue-specific contributions to heritability were non-significant for both constituent traits. Numerical results are reported in **Supplementary Tables 18-19**. BMI: body mass index. RBC Count: red blood cell count. WBC Count: white blood cell count. LDL: low-density lipoprotein. Yrs Edu: years of education. WHRadjBMI: waist-hip-ratio adjusted for body mass index. Accumbens Vol: brain accumbens volume. Caudate Vol: brain caudate volume. MDD: major depressive disorder. Scz: Schizophrenia. T2D: type 2 diabetes. FVC: forced vital capacity. RA: rheumatoid arthritis. (B) For BMI and red blood cell count (RBC Count), we report estimates of the proportion of trait heritability for each trait and signed proportion of genetic covariance explained by the *cis*-genetic component of gene expression in pancreas. Lines with asterisks denote significant differences at 10% FDR between respective estimates, assessed by jackknifing the differences. (C) For BMI and red blood cell count (RBC Count), we report estimates of the proportion of trait heritability for each trait and proportion of genetic covariance explained by the *cis*-genetic component of gene expression in the brain substantia nigra. Lines with double asterisks denote significant differences at 5% FDR between respective estimates, assessed by jackknifing the differences. Numerical results are reported in **Supplementary Table 21**.

TCSC identified several biologically plausible findings not previously reported in the genetics literature. First, brain substantia nigra had a significantly positive contribution to the genetic covariance of BMI and red blood cell count (RBC count) (*ζ_t’_* = 0.28, s.e. = 0.084, *P* = 4.6 × 10^-4^), while pancreas had a significantly negative contribution (*ζ_t’_* = −0.25, s.e. = 0.079, *P* = 8.7 × 10^-4^). In the brain, energy metabolism is regulated by oxidation and previous work has shown that red blood cells play a large role in these metabolic processes as oxygen sensors^81^; in addition, previous studies have reported differences in the level of oxidative enzymes in red blood cells between individuals with high BMI and low BMI^82,83^, suggesting that genes regulating oxidative processes might have pleiotropic effects on RBC count and BMI. In the pancreas, pancreatic inflammation (specifically acute pancreatitis) is associated with reduced levels of red blood cells, or anemia^84^, while pancreatic fat is associated with metabolic disease and increased BMI^85^. Once again, the contrasting results for brain substantia nigra and pancreas suggest that genetic covariance may reflect distinct tissue-specific contributions. Second, brain substantia nigra had a significantly negative contribution to the genetic covariance of age at first birth and height (*ζ_t’_*, = −0.11, s.e. = 0.032, *P* = 4.5 × 10^-4^). Previous work in *C. elegans* reported that fecundity is positively regulated by dopamine^86,87^, which is produced in the substantia nigra^88^. Therefore, it is plausible that reproductive outcomes related to fecundity, such as age at first birth, are also regulated by dopamine via the substantia nigra. Dopamine also plays a role in regulating the levels of key growth hormones such as IGF-1 and IGF-BP3^89^ and has been previously shown to be associated with height^90^. Third, pituitary had a significantly negative contribution to the genetic covariance of vitamin D and WHR | BMI (*ζ_t’_* = −0.19, s.e. = 0.057, *P* = 4.5 × 10^-4^). Irregularities in pituitary development are associated with decreased vitamin D levels and decreased IGF-1 levels, the latter of which is integral for bone development and is directly proportional to body proportion phenotypes such as WHR | BMI^91–93^. Fourth, LCLs had a suggestive negative contribution to the genetic covariance of eosinophil count and white blood cell count (*ζ_t’_* = −0.081, s.e. = 0.028, *P* = 1.8 × 10^-3^, 5% < FDR < 10%, in contrast to the suggestive positive contribution of whole blood: *ζ_t’_* = 0.32, s.e. = 0.12, *P* = 2.4 × 10^-3^, 5% < FDR < 10%). This is plausible as previous studies have reported the suppression of proliferation of lymphocytes (the white blood cell hematopoietic lineage from which LCLs are derived) by molecules secreted from eosinophils^94–96^. The contrasting results for whole blood and LCLs suggest that genetic covariance may reflect distinct tissue-specific contributions. Other significant findings are discussed in the **Supplementary Note**. Numerical results for all tissues and disease/trait pairs analyzed are reported in **Supplementary Table 19**.

As noted above, for 16 of the 17 causal tissue-trait covariance pairs, the causal tissue was non-significant for both constituent traits. We sought to formally assess whether differences in tissue-specific contributions to genetic covariance vs. constituent trait heritability were statistically significant. Specifically, for each causal tissue-trait covariance pair, we estimated the differences between the tissue-specific contribution to covariance (*ζ_t’_*) and the tissue-specific contributions to heritability for each constituent trait (*π_t’_*) (and estimated standard errors by jackknifing differences across the genome). We note that *ζ_t’_* and *π_t’_* are both signed proportions and are therefore on the same scale, thus the scenario in which these two quantities are equal is a natural and parsimonious null. We identified five tissue-trait covariance pairs for which these differences were statistically significant at 5% FDR for both constituent traits and *π_t’_* was non-significant for both constituent traits (marked by double asterisks in **Figure 6A**, **Supplementary Table 20**). For BMI and RBC count, negative contribution of pancreas (**Figure 6B**) and the positive contribution of brain substantia nigra (**Figure 6C**) to genetic covariance were each larger than the respective contributions of those tissues to BMI and RBC count heritability, which were non-significant. Other examples are discussed in the **Supplementary Note**. Numerical results for all tissues and trait pairs are reported in **Supplementary Table 20**. These findings were consistent with simulations we performed in which TCSC frequently detected tissue-specific contributions to covariance while failing to detect tissue-specific contributions to heritability for *both* traits, both in our original simulation framework and in a new simulation framework in which tissue-specific contributions to covariance were greater than contributions to heritability (**Supplementary Table 21**).

## Discussion

We developed a new method, tissue co-regulation score regression (TCSC), that disentangles causal tissues from tagging tissues and partitions disease heritability (or genetic covariance of two diseases/traits) into tissue-specific components. We applied TCSC to 78 diseases and complex traits and 48 GTEx tissues, identifying 21 tissue-trait pairs (and 17 tissue-trait covariance pairs) with significant tissue-specific contributions. TCSC identified biologically plausible novel tissue-trait pairs, including associations of aorta artery with glaucoma, esophagus muscularis with FEV1/FVC, and heart left ventricle with platelet count. TCSC also identified biologically plausible novel tissue-trait covariance pairs, including a negative contribution of LCLs to the covariance of eosinophil count and white blood cell count (in contrast to the positive contribution of whole blood) and a positive contribution of brain substantia nigra and a negative contribution of pancreas to the covariance of BMI and red blood cell count; in particular, our findings suggest that genetic covariance may reflect distinct tissue-specific contributions.

TCSC differs from previous methods in jointly modeling contributions from each tissue to disentangle causal tissues from tagging tissues (analogous to the distinction in GWAS between marginal association and fine-mapping^23^). We briefly discuss several other methods that use eQTL or gene expression data to identify disease-associated tissues. RTC Coloc identifies disease-associated tissues based on tissue specificity of eQTL-GWAS colocalizations^2^; this study made a valuable contribution in emphasizing the importance of tissue co-regulation, but did not model tissue-specific effects, such that RTC Coloc may implicate many tissues (**Figure 5A**). LDSC-SEG identifies disease-critical tissues based on heritability enrichment of specifically expressed genes^7^; this distinguishes a focal tissue from the set of all tissues analyzed, but does not distinguish closely co-regulated tissues (**Figure 5B**). MaxCPP models contributions to heritability enrichment of fine-mapped eQTL variants across tissues or meta-tissues^4^; although this approach proved powerful when analyzing eQTL effects that were meta-analyzed across all tissues, it has limited power to identify disease-critical tissues: fine-mapped eQTL annotations for blood (resp. brain) were significant conditional on annotations constructed using all tissues only when meta-analyzing results across a large set of blood (resp. brain) traits (Fig. 4 of ref.^4^). eQTLenrich compares eQTL enrichments of disease-associated variants across tissues^3^; this approach produced compelling findings for eQTL that were aggregated across tissues, but tissue-specific analyses often implicated many tissues (Fig. 1d of ref.^3^). MESC estimates the proportion of heritability causally mediated by gene expression in assayed tissues^97^; this study made a valuable contribution in its strict definition and estimation of mediated effects (see below), but did not jointly model distinct tissues and had limited power to distinguish disease-critical tissues (Fig. 3 of ref.^97^). CAFEH leverages multi-trait fine-mapping methods to simultaneously evaluate all tissues for colocalization with disease^5^; however, this locus-based approach does not produce genome-wide estimates and it remains the case that many (causal or tagging) tissues may colocalize with disease under this framework. Likewise, methods for identifying tissues associated to disease/trait covariance do not distinguish causal tissues from tagging tissues^98,99^.

We note several limitations of our work. First, TCSC requires tissue-specific eQTL data (thus requiring genotype/gene expression data in substantial sample size), whereas some methods (LDSC-SEG^7^, RolyPoly^6^, and CoCoNet^9^) only require gene expression data in limited sample size. However, TCSC attains lower type I error and higher AUC than those methods in our simulations (**Supplementary Figure 1**); and its results are generally consistent in independent GWAS data (**Supplementary Table 5**), although all methods likely produce some false positives. Moreover, methods that only use gene expression data exclude contributions to disease from genes that are ubiquitously expressed but have cell-type-specific functionality or cell-type-specific genetic regulation such as transcription factors, which are widely believed to orchestrate large transcriptional programs important to disease^100^. Second, joint-fit effects of gene expression on disease may not reflect biological causality; if a causal tissue or cell type is not assayed^101^, TCSC may identify a co-regulated tissue (e.g. a tissue whose cell type composition favors a causal cell type) as causal or may identify a set of co-regulated tissues that collectively tag the causal tissue as causal. We anticipate that this limitation will become less severe as potentially causal tissues, cell types and contexts are more comprehensively assayed. Third, TCSC does not achieve a strict definition or estimation of mediated effects; this is conceptually appealing and can, in principle, be achieved by modeling non-mediated effects, but may result in limited power to distinguish disease-critical tissues^97^. Fourth, TCSC has low power at small eQTL sample sizes; in addition, TCSC estimates are impacted by the number of significantly *cis*-heritable genes in a focal tissue, which can lead to conservative bias at small eQTL sample sizes. We anticipate that these limitations will become less severe as eQTL sample sizes increase. Fifth, TCSC is susceptible to large variations in eQTL sample size, which may compromise type I error; therefore, there is a tradeoff between maximizing the number of tissues analyzed and limiting the variation in eQTL sample size. Sixth, TCSC assumes that causal gene expression-disease effects are independent across tissues; this assumption may become invalid for tissues and cell types assayed at high resolution. However, we verified via simulations that TCSC performs well when this model assumption is violated (**Supplementary Figures 17-20**). Seventh, TCSC does not formally model measurement error in tissue coregulation scores, but instead applies a heuristic bias correction. We determined that the bias correction generally performs well in simulations. Eighth, TCSC does not produce locus-specific estimates or identify causal tissues at specific loci. However, genome-wide results from TCSC may be used as a prior for locus-based methods (analogous to GWAS fine-mapping with functional priors^102^). Ninth, TCSC performs less well in the presence of disease heritability that is not mediated through gene expression (**Supplementary Figures 21-22**). Tenth, we did not apply TCSC to single-cell RNA-seq (scRNA-seq) data, which represents a promising new direction as scRNA-seq sample sizes increase^103–105,35^; we caution that scRNA-seq data may require new eQTL modeling approaches^103^. Finally, we focused our cross-trait analyses on relatively independent traits from the single-trait analysis, to enable comparisons with single-trait results (**Figure 6B, 6C**); cross-trait analysis of more strongly genetically correlated traits is a future direction of high interest. Despite these limitations, TCSC is a powerful and generalizable approach for modeling tissue co-regulation to estimate tissue-specific contributions to disease.

## Supporting information

TCSC_SupplementaryTables_Rev

TCSC_SupplementaryNote_Rev

## Code Availability

TCSC software including a quick start tutorial: https://github.com/TiffanyAmariuta/TCSC/Mancuso Lab TWAS Simulator: https://github.com/mancusolab/twas_sim.

FUSION software: http://gusevlab.org/projects/fusion/.

## Data Availability

We have made 78 GWAS summary statistics and 41 brain-specific summary statistics publicly available at https://github.com/TiffanyAmariuta/TCSC/tree/main/sumstats, TWAS association statistics publicly available at https://alkesgroup.broadinstitute.org/TCSC/TWAS_sumstats/, tissue co-regulation scores publicly available at https://github.com/TiffanyAmariuta/TCSC/tree/main/coregulation_scores, and TCSC output publicly available at https://github.com/TiffanyAmariuta/TCSC/tree/main/results.

## Acknowledgements

We thank Huwenbo Shi, Martin Zhang, and Benjamin Strober for helpful discussions. This work was funded by NIH grants U01 HG009379, R01 MH101244, R37 MH107649, R01 HG006399, R01 MH115676 and U01 HG012009.

## Methods

### TCSC regression

TCSC leverages the fact that the TWAS *χ*^2^ statistic for a gene-tissue pair includes the direct effects of the gene on the disease as well as the tagging effects of co-regulated tissues and genes with shared eQTLs or eQTLs in LD. Thus, genes that are co-regulated across many tissues will tend to have higher *χ*^2^ statistics than genes regulated in a single tissue. TCSC determines that a tissue causally contributes to disease if genes with high co-regulation to the tissue have higher TWAS *χ*^2^ statistics than genes with low co-regulation to the tissue.

We model the genetic component of gene expression as a linear combination of SNP-level effects:

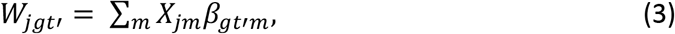

where *W_jgt_* is the *cis*-genetic component of gene expression in individual *j* for gene *g* and tissue *t’*, *X_jm_* is the standardized genotype of individual *j*for SNP *m*, and *β_gtm_* is the standardized effect of the *m^th^* SNP on the *cis*-genetic component of gene expression of gene *g* in tissue *t’*. We define the *cis*-genetic component of gene expression *W_jgt_* to have mean 0 and variance 1 and *β_gt’m_* to have mean 0 and variance 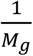, where *M_g_* is the number of *cis* variants for gene *g*.

TCSC assumes that true gene-disease effects are identically distributed (i.i.d.) across genes and tissues while accounting for the fact that *cis*-genetic components of gene expression (and *cis*-genetic predictions of gene expression) are correlated^1^ (see **Supplementary Figures 17-20** for simulations where gene expression-trait effect sizes are not i.i.d. across genes and tissues; TCSC performs well despite violations of model assumptions). The high correlation of *cis-*eQTLs across tissues leads to tagging from co-regulated tissues^2^. We model phenotype as a linear combination of genetic components of gene expression across genes in different tissues:

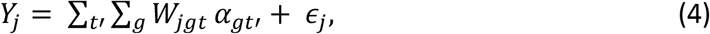

where *Y_j_* is the (binary or continuous-valued) phenotype of individual *j*, *α_gt_* is the standardized effect size of the *cis*-genetic component of gene expression on disease and *∈_j_* is the component of phenotype not explained by cis-genetic components of gene expression. We emphasize that we model disease as a function of the unobserved true *cis*-genetic component of gene expression *W_jgt’_*, *not* the genetically predicted value 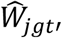 obtained from gene expression prediction models. Equation (4) can be rewritten in terms of SNP-level effects:

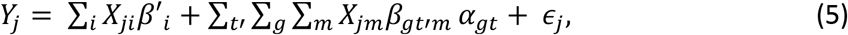

where *β’_t_* are direct SNP-disease effects not mediated through gene expression.

We define the disease heritability explained by *cis*-genetic expression across all tissues as follows:

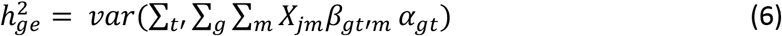

Because *W_jgt_* has mean 0 and variance 1 and *α_gt_* are assumed to be i.i.d. across genes and tissues (see above), Equation (5) implies that:

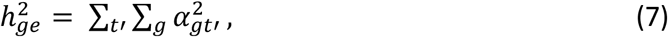

analogous to the relationship between SNP effect sizes and SNP-heritability^27^: 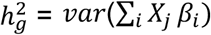. We emphasize that the respective terms in Equation (5) for each tissue *t’* are independent as *α_gt’_* are assumed to be i.i.d. across genes and tissues. It follows that the disease heritability explained by a particular tissue *t’* is

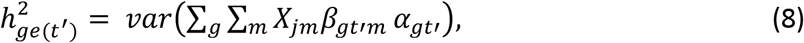

which given that *W_jgt’_* has mean 0 and variance 1 and *α_gt’_* is i.i.d. across genes, reduces to:

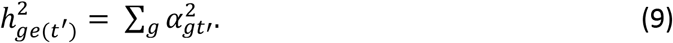

Equation (7) and Equation (9) imply that 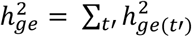. Now, let *α_gt’_* be a random variable drawn from a normal distribution with mean zero and tissue-specific variance *var*(*α_gt’_*) = *τ_t’_*. Then

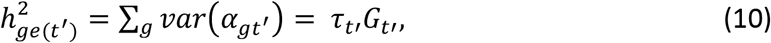

where *G_t’_* is the number of significantly *cis*-heritable genes in the model. In simulations, we demonstrate that when there are similar numbers of *cis*-heritable genes across tissues, setting *G_t’_* to the total number of unique *cis*-heritable genes produces unbiased estimates in TCSC for the causal tissue; however, when there are varying numbers of *cis*-heritable genes across tissues (fewer in the causal tissue), this produces upward biased estimates (**Supplementary Figures 3-4**) and thus setting *G_t’_* to the number of significantly *cis*-heritable genes in tissue *t’* is recommended. With this variance term, we can define a polygenic model that relates TWAS *χ*^2^ statistics to co-regulation scores, which explicitly model the covariance structure of the *χ*^2^ statistics. This strategy is analogous to modeling the dependence of GWAS *χ*^2^ statistics on LD scores^27^.

In a TWAS, the estimated value of the gene-disease effect size *α_gt’_* is proportional to the correlation of the *cis*-genetic components of gene expression and their true gene-disease effect sizes for nearby genes across tissues, analogous to GCSC^22^:

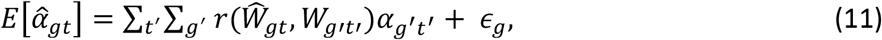

where 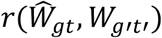 is the estimated correlation in *cis*-genetic predicted expression between gene *g in tissue t* and genes *g’* in tissue *t’. ∈_g_* is the component of phenotype not explained by *cis*-genetic components of gene expression, with mean 0 and variance 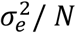.

The value of the TWAS *χ*^2^ is proportional to the squared estimated disease-gene effect size and the GWAS sample size *N* as follows:

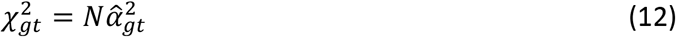

Using the equations (9) and (10), we can write the expectation of TWAS *χ*^2^ as follows:

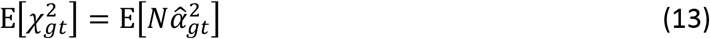

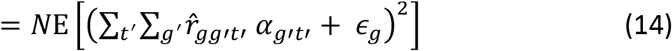

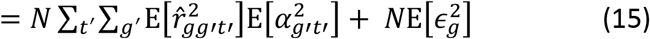

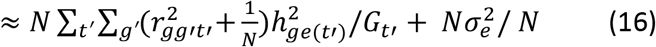

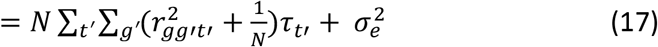

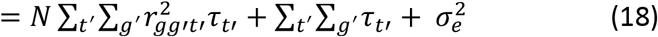

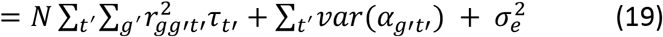

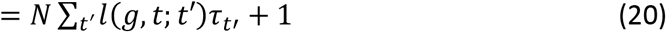

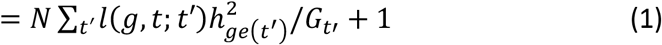

To go from Equation (15) to Equation (16) we use the following relationship from the derivation of LDSC^13^:

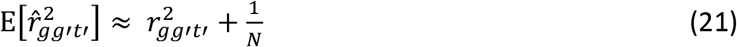

We go from Equation (19) to Equation (20) because the variance of the phenotype *Y_j_* is 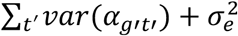 and is equal to one. We also introduce the notation that 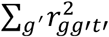 are the tissue and gene co-regulation scores *l*(*g*, *t*; *t’*), see below. We are interested in estimating *τ_t’_*, the per-gene disease heritability explained by the cis-genetic component of gene expression in tissue *t’*. From the derivation, the genome-wide tissue-specific contribution to disease heritability is estimated as

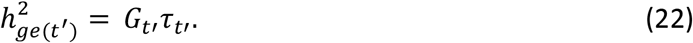

For the analysis of tissue-specific contributions to the covariance between two diseases, we can extend TCSC by using products of TWAS z-scores. Following the polygenic model described above, the expected product of TWAS z-scores in disease 1 and disease 2 for gene *g* and tagging tissue *t* is

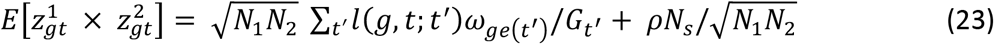

where *N*_1_ is GWAS sample size for disease 1, *N*_2_ is GWAS sample size for disease 2, *t’* indexes causal tissues, *l*(*g*, *t*; *t’*) are tissue co-regulation scores (see below), *ω*_*ge*(*t’*)_ is the genetic covariance explained by the *cis*-genetic component of gene expression in tissue *t’*, *G_t’_* is the number of significantly *cis*-heritable genes in tissue *t’* (see below), *ρ* is the phenotypic correlation between disease 1 and disease 2, and *N_s_* is the number of overlapping GWAS samples between disease 1 and disease 2. The last term represents the intercept^28^, and while we use a free intercept in the multivariate regression on co-regulation scores, the estimation of this term only plays a role in the estimation of regression weights (see below).

For estimates of 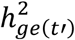 and *ω*_*ge*(*t’*)_, we use a free intercept; the estimation of 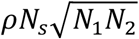 serves only to inform the heteroscedasticity weights (see below) and is not used in the multivariate TCSC regression to estimate *ω*_*ge*(*t’*)_. To estimate standard errors, we use a genomic block jackknife over 200 genomic blocks with an equal number of genes in each. The standard deviation is computed as the square root of the weighted variance across the jackknife estimates (where the weight of each block is equal to the sum of the regression weights for the genes in that block) multiplied by 200 blocks. We expect that the jackknife standard error will be conservative relative to the empirical standard error across estimates due to variation in causal signal across loci^106^.

### Estimating tissue co-regulation scores and correcting for bias

We define the co-regulation score of gene *g* with tissues *t* and *t’* as

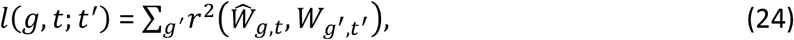

where *W* denotes the *cis*-genetic component of gene expression for a gene-tissue pair across individuals, 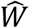 denotes the *cis*-predicted expression for a gene-tissue pair, and genes *g’* are within +/- 1 Mb of the focal gene *g*. TCSC corrects for bias in tissue co-regulation scores arising from differences between *cis*-genetic vs. *cis*-predicted expression (analogous to GCSC^22^). We apply bias correction to co-regulation scores in the special case whe n *g* = *g’* and *t* = *t’*. While co-regulation scores aim to estimate 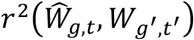, the squared correlation of the predicted *cis*-genetic component of expression of gene *g* and tissue *t* (corresponding to the TWAS 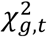 statistic) with the *actual cis*-genetic component of gene expression of gene *g’* in tissue *t’*, when *g* = *g’* and *t* = *t’*, the estimated value of 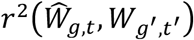 will always equals one because the estimate is based on 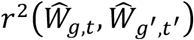. However, this implies that predictions of the cis-genetic component of expression are perfectly accurate, which is unlikely to be the case. Therefore, the estimated value of 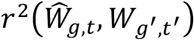 if left to equal one will cause co-regulation scores to be systematically inflated.

Therefore, when *g* = *g’* and *t* = *t’*, we set

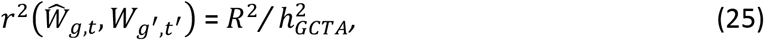

where *R*^2^ is the cross-validation prediction statistic of the gene expression model for gene *g* in tissue *t* and 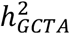 is the GCTA-estimated *cis*-heritability of gene expression for gene *g* in tissue *t*. The quotient 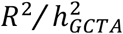 is the accuracy of the gene expression prediction model, which reflects the upper bound on how much the *cis*-predicted expression can be correlated with the true *cis*-genetic component of gene expression. While we only consider genes with 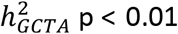, the uncertainty in 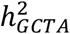 estimates should be modest and therefore not greatly impact our bias correction. We note that TWAS tests the null hypothesis that a specific weighted linear combination of SNPs is not associated with disease (and does not test the null hypothesis that the *cis*-genetic component of gene expression is not associated with disease).

### TCSC regression weights

TCSC uses three sets of regression weights to increase power (analogous to GCSC^22^). The first regression weight is inversely proportional to *L*(*g, t*), the total co-regulation score of each gene-tissue pair summed across tissues *t’*:

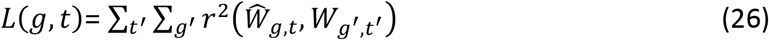

(without applying bias correction; see above), which allows TCSC to properly account for redundant contributions of co-regulated genes to TWAS *χ*^2^ statistics.

The second regression weight is inversely proportional to *T*(*g, t*), the number of tissues in which a gene is significantly *cis-*heritable:

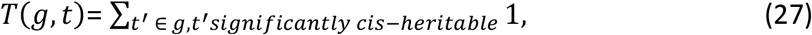

thereby up-weighting signal from genes that are regulated in a limited number of tissues and preventing TCSC from attributing more weight to genes that are co-regulated across many tissues.

The third regression weight is inversely proportional to *H*_*h*^2^_(*g, t*), the heteroscedasticity of *χ*^2^ statistics, and is computed differently for estimates of 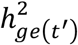 than for estimates of *ω*_*ge*(*t’*)_, (analogous to GCSC^22^ and cross-trait LDSC^28^, respectively).

For estimates of 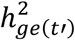, we estimate *H*_*h*^2^_(*g, t*) in two steps. First, we make a crude estimate of heritability explained by predicted expression 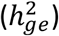 as follows:

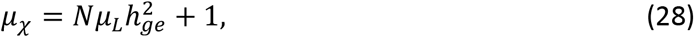

where *μ_χ_* is the mean *χ*^2^ statistic:

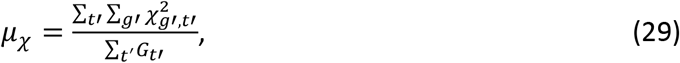

where *N* is the GWAS sample size, *g’* iterates over significantly *cis*-heritable genes and *t’* iterates over tissues, and *μ_L_* is the mean value of total co-regulation across tissues *t’*,

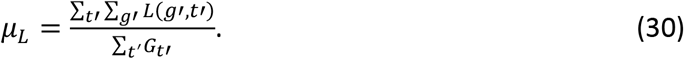

Then, we compute the heteroscedasticity for each significantly *cis*-heritable gene-tissue pair as

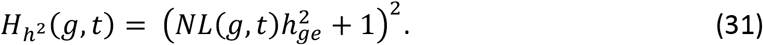

Finally, we combine the three regression weights as follows:

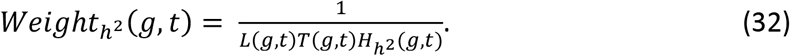

For estimates of *ω*_*ge*(*t’*)_, we estimate *H_ω_*(*g, t*) in two steps. First, we regress the products of TWAS z-scores on total tissue co-regulation scores, *L*(*g, t*), using regression weights, *Weight_ω_*(*g, t*), computed as follows:

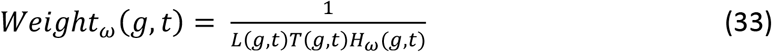

where *H_ω_*(*g, t*) is *first* estimated as follows:

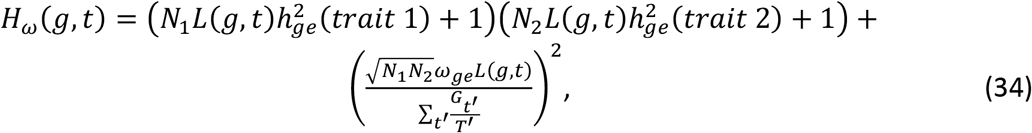

where 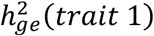 is the crude heritability estimate for trait 1 and 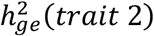 is the crude heritability estimate for trait 2, *ω_ge_* is estimated as 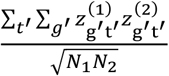, *N*_1_ is the sample size of the first GWAS, *N*_2_ is the sample size of the second GWAS, and *T’* is the total number of tissues in the regression.

Second, we use the regression intercept to estimate the product *ρN_s_*:

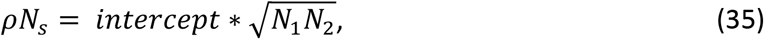

where *ρ* represents the phenotypic correlation between trait 1 and 2 and *N_s_* represents the number of shared samples between GWAS 1 and 2. We also use the coefficient of the regression to update our estimate of *ω_ge_*, such that we may update the heteroscedasticity weight as follows:

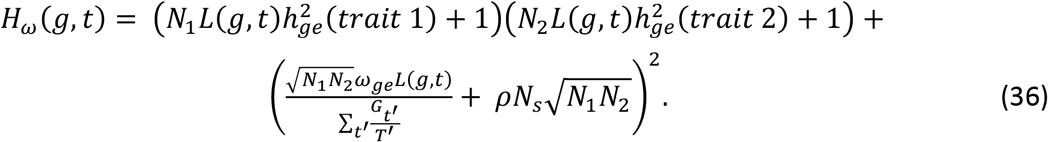

Finally, we combine the three regression weights as follows:

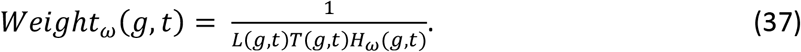

### Simulating TCSC

We employed a widely used TWAS simulation framework (Mancuso Lab TWAS Simulator, see **Code Availability**) to assess the power, bias, and calibration of TCSC in the presence of co-regulation across genes and tissues. We simulated a genome in which there are 1,000 protein-coding genes from chromosome 1, of which 100 (10%) are causal^31^. Each primary simulation consists of 10 tissues, of which at least one is causal, defined as having nonzero gene-disease effect sizes. We create a covariance structure among tissues mimicking empirical GTEx data. We use a previously published method to estimate the causal cross-tissue correlation of eQTL effect sizes which is 0.75^36^. We observe that not all GTEx tissues are equally correlated to one another. We estimate three different cross-tissue eQTL correlation quantities: (1) average correlation across all pairs of tissues = 0.75, (2) average correlation across similar tissues = 0.80, e.g. brain (13 in GTEx) or adipose (2 in GTEx) tissues, and (3) average correlation across dissimilar tissues, e.g. pairs of brain and adipose tissues = 0.74. To represent these biological modules, we let simulated tissues 1-3 have higher correlation of true eQTL effects to one another than to other tissues; likewise for tissues 4-6 and 7-10. We set covariance parameters, described below, such that the similar tissues had an average eQTL correlation of 0.789 across genes, dissimilar tissues have an average eQTL correlation of 0.737, and the average eQTL correlation across any pair of tissues is 0.751. We use real genotypes from European individuals in the 1000 Genomes Project to define the pairwise SNP LD structure which is used to simulate genotypes, gene expression traits, and complex traits/diseases. We simulate each gene having 5 true *cis*-eQTLs, based on the upper bound of empirical data from GTEx^19^ and others^35^, as well as the value used in other TWAS simulation methods^34^. Between pairs of co-regulated tissues, the same gene shares 3 *cis*-eQTLs. Between pairs of co-regulated genes in the same tissue, 3 *cis*-eQTLs are shared. The minimum allowed *cis-*heritability of a gene is 0.01 in our simulations. *Cis-*heritability is approximated as the sum of squared true *cis*-eQTL effect sizes, as done previously^22^. Effect sizes for the 3 shared eQTLs across tissues are sampled from a multivariate normal distribution with mean 0 and a variance-covariance matrix. We define the variance and covariance terms of this matrix such that (1) the proportion of genes detected as significantly *cis-*heritable by GCTA at a given sample size and (2) the average *cis* heritability of detected genes at a given sample size match empirical observations from GTEx data at sample sizes N = 100, 200, 300 and 500. As a result, the diagonal of the variance-covariance matrix, e.g. the variance term, is set to 0.075, and the off-diagonal elements are set to the product of the variance term and the desired correlation for each tissue pair, described above.

For each of 1,000 independent simulations per analysis, we simulate a GWAS (N = 10,000) by creating a complex trait which is the summation of the genetic components of causal gene expression (in the causal tissue). We use simulated genotypes based on the LD structure of 1000 Genomes. Gene-disease effect sizes are drawn from a normal distribution with mean 0 and variance 1. In cross-trait TCSC analysis, effect sizes across genes between the two traits are correlated with default *R_g_* = 0.5. To simulate a GWAS trait, we first compute the genetic component of each gene, which is the product of GWAS cohort genotypes and eQTL effects, such that we have 100 gene-specific traits. We then add noise to each gene-specific trait such that the total variance of the phenotype explained by the five eQTLs from the causal tissue is equal to a specified value; the value of 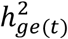 in primary simulations is 10%. Then, we multiply each gene-specific trait by the causal gene-disease effect size, consistent with the additive generative model of gene-level effects on trait (see above). Finally, we take the sum across all gene-specific traits to make one complex trait, where the total variance of the trait explained by gene effects from the causal tissue is 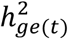, e.g. 10%.

We simulate an eQTL cohort of various gene expression sample sizes (N = 100, 200, 300, 500, 1000, 1500) using simulated genotypes based on the LD structure of 1000 Genomes. We simulate total gene expression in the eQTL cohort by adding a desired amount of noise to the genetic component of gene expression, e.g. the product of individual genotypes and true eQTL effect sizes, with variance equal to one minus the gene expression heritability, which is the sum of squared eQTL effects. Next, we fit gene expression prediction models by regressing the total gene expression on eQTL cohort genotypes of *cis* variants using lasso regularization, a standard approach used in TWAS. We define significantly *cis-*heritable genes as genes with GCTA heritability *P* value < 0.01^21^ and heritability estimate > 0, and adjusted-*R^2^* > 0 in cross-validation prediction.

Then we estimate co-regulation scores at each different eQTL sample size by predicting gene expression into a cohort of 500 individuals, to approximate the size of the European sample of 1000 Genomes (*N* = 489). Using significantly *cis*-heritable genes from each tissue at a given sample size, we estimate gene and tissue co-regulation scores *l*(*g, t; t’*) as described above, including bias correction. In simulations, *cis* genes are defined as genes within the same 1 Mb block.

Then we apply TWAS to individual-level simulated GWAS data and gene expression prediction models. We predict gene expression into each of the 10,000 GWAS cohort individuals across all significantly *cis-*heritable genes for each tissue. We regress each complex trait on predicted gene expression to obtain TWAS z-scores. Finally, we run TCSC by regressing TWAS *χ*^2^ statistics, or products of TWAS z-scores, on bias-corrected gene and tissue co-regulation scores.

### Simulating other tissue-disease association methods

We simulated four tissue-trait association methods: RTC Coloc^2^, LDSC-SEG^7^, RolyPoly^6^, or CoCoNet^9^. First, we simulated RTC Coloc method^2^ by leveraging our existing TCSC simulation framework such that both methods could be compared via application to same simulated data. We used the same simulated GWAS cohort of 10,000 individuals as in our TCSC simulations and then followed the steps of the RTC Coloc method as published. Briefly, we perform a genomewide association study using our simulated complex trait and the genotypes of our simulated GWAS cohort and select null variants with similar LD properties. Then, we simulate an eQTL cohort consisting of total gene expression and genotypes, using the same underlying true eQTL effect sizes as for TCSC simulations. Then, we perform colocalization analysis of GWAS variants with eQTLs, across 10 tissues at 6 different eQTL sample sizes, to obtain the regulatory trait concordance (RTC) score. This is repeated for the set of null variants. Next, we perform colocalization analysis of eQTL variants between pairs of tissues to obtain tissue-sharing RTC scores, and similarly repeat this for null variants. GWAS-eQTL RTC scores are divided by tissue-sharing RTC scores summed across variants. Tissue-specific enrichment is computed as the ratio of this quotient to the null quotient. The enrichment *P* value is obtained using a Wilcox test comparing the values of the quotient to the values of the null quotient.

Second, we simulated the three methods that utilize GWAS data and total expression across tissues: LDSC-SEG^7^, RolyPoly^6^, and CoCoNet^9^. To this end, we retained the full GWAS summary statistics from the RTC Coloc analysis above. We separately simulated total expression across tissues in which the 100 causal genes in addition to 200 randomly selected genes were positively differentially expressed in the causal tissue and the two tagging tissues in the same simulated “module” as the causal tissue, e.g. with higher genetic correlation of gene regulatory effects. We also selected 100 random non-causal genes to be negatively differentially expressed in the causal tissue and the other two module tissues. For the remaining 7 tagging tissues, we randomly selected 300 genes to be positively differentially expressed, some of which at random will be causal genes, and let the remaining 700 genes be negatively differentially expressed. Then, as previously done^7^, we calculated the t-statistics for the specific expression of each gene in each tissue. While we have modules of tissues that are more highly correlated to one another, these within-module tissues were excluded from the calculation of t-statistics, as previously done^7^. Finally, we created SNP-based annotations for each tissue, across 1000 simulations, and across 6 sample sizes, in which SNPs within +/- 100 kb of a specifically expressed gene is assigned a value of 1 and 0 otherwise, as previously done^7^. Then, we calculated LD scores and partitioned the heritability of our simulated complex traits. For the simulations of RolyPoly and CoCoNet, we installed the following R packages: rolypoly and CoCoNet and used the simulated data above to run each method. While CoCoNet does not technically use GWAS summary statistics, but rather gene-based “outcome variables”, we used the label of causal or non-causal for each gene in each tissue of our simulations as the outcome variable.

### Gene expression prediction models and tissue co-regulation scores in GTEx data

We downloaded GTEx v8 gene expression data for 49 tissues. We excluded tissues with fewer than 100 samples, e.g. kidney cortex (n = 69). We retained only European samples for each tissue, as labeled by GTEx via PCA of genotypes. We constructed gene expression models for two scenarios: (1) subsampling to 320 individuals including meta-analyzed tissues (**Table 1**) or (2) using all European samples per tissue. We recommend meta-analyzing gene expression prediction models across tissues in the case of tissues with low eQTL sample size (e.g. < 320 samples) and high pairwise genetic correlation (e.g. > 0.93). We determined in simulations that TCSC is sensitive to eQTL sample size differences, such that a tagging tissue with larger sample size than a causal tissue can produce false positive results; the subsampling approach was designed to mitigate this issue. For the subsampling procedure, we first set aside tissues with more than 320 samples; we chose 320 based on the average GTEx tissue sample size (*N* = 271) and robustness of TCSC in simulations at *N* = 300. Then, we grouped tissues with genetic correlation, e.g. marginal effect size correlation as reported by GTEx, with *R_g_* > 0.93, an arbitrary threshold that produced biologically plausible groups of related tissues, separating groups of brain tissues based on cranial compartment. We meta-analyzed gene expression prediction models for these grouped tissues in order to achieve a total sample size of 320 individuals where each tissue contributed an approximately equal number of samples, using an inverse-variance weighted meta-analysis across genes that were significantly *cis*-heritable in two or more constituent tissues. The prediction weights of genes that were significantly *cis*-heritable in a single constituent tissue were left unmodified.

To construct gene expression prediction models, we applied FUSION^21^ (**Code Availability**) to individual-level GTEx data by regressing measured gene expression on genotypes of common variants (MAF > 0.05) and covariates provided by GTEx^19^. FUSION uses several different regression models: single eQTL, elastic net, lasso, and BLUP and the following covariates: sex, 5 genotyping principal components, PEER factors^107^, and assay type. We defined significantly *cis-*heritable genes as protein-coding genes with GCTA heritability *p* < 0.01^21^, heritability estimate > 0, and adjusted-*R^2^* > 0 in cross-validation prediction.

We used gene expression prediction models of significantly *cis*-heritable genes to predict expression into 489 European individuals from 1000 Genomes^108^. We then estimated tissue co-regulation scores using Equation (24) and Equation (25), where *cis*-predicted gene expression is used to estimate the *cis*-genetic component of gene expression.

### GWAS summary statistics and TWAS association statistics

We collected GWAS summary statistics from 78 relatively independent heritable complex diseases and traits (average N = 302K) with heritability z-score > 6. We estimated the heritability of all summary statistics and genetic correlation of all pairs of summary statistics. We excluded traits with heritability z-score < 6, using S-LDSC with the baseline-LD v2.2 model^13,25,26^ and as done previously^25^. We excluded one of each pair of traits that are *both* genetically correlated and have significantly overlapping samples. Specifically, for any pair of non-UK Biobank traits with an estimated sample overlap greater than the following threshold-squared cross-trait LDSC intercept / (trait 1 S-LDSC intercept * trait 2 S-LDSC intercept) > 0.1^28^-the trait with the larger SNP heritability z-score was retained. For any pair of UK Biobank traits with a squared genetic correlation > 0.1, the trait with the larger SNP heritability z-score was retained^38^. In total, this procedure resulted in 78 sets of relatively independent GWAS summary statistics. We limited all analyses (including cross-trait analyses) to the 78 relatively independent traits in order to avoid redundant findings across single-trait (and cross-trait) analyses. For the brain-specific analysis, we first selected brain-related diseases and complex traits, e.g. psychiatric disorders and behavioral phenotypes, excluding multi case-control studies and case vs case studies. Then, we applied our standard filters as described above, but relaxing the threshold of squared genetic correlation to 0.25.

We used FUSION^21^ (**Code Availability**) to compute TWAS association statistics for each pair of signed GWAS summary statistics and each significantly *cis-*heritable gene-tissue pair, across the two scenarios described above. We further removed genes within the MHC (chromosome 6, 29 Mb - 33 Mb) and TWAS *χ*^2^> 80 or *χ*^2^> 0.001*N*, where *N* is the GWAS sample size, as previously used for quality control in the heritability analysis of GWAS summary statistics^13^. TCSC scales linearly with the number of genes and quadratically with the number of tissues. After all input datasets are created and processed, running TCSC on a single real GWAS trait with 39 tissues takes about two minutes.

### RTC Coloc and LDSC-SEG analysis of GWAS summary statistics and GTEx tissues

We downloaded supplementary tables for the RTC coloc method^2^ and for LDSC-SEG^7^. For traits in our set of 78 GWAS summary statistics that were not analyzed by the LDSC-SEG study and for traits that are inherently brain-related (as these traits require a different procedure for generating tissue-specific gene sets), we ran LDSC-SEG ourselves. To this end, we downloaded LD scores for GTEx tissues and specifically expressed gene set SNP-level annotations (https://alkesgroup.broadinstitute.org/LDSCORE/LDSC_SEG_ldscores/) and ran LDSC-SEG as previously described^7^. For brain-related traits, we additionally ran a brain-specific analysis using LDSC-SEG, also as previously described^7^. Briefly, specifically expressed genes were determined via a t-test of the sentinel brain tissue against all other brain tissues, rather than against all other non-brain GTEx tissues, as done in the primary analysis of the LDSC-SEG study. For traits in our set that were not analyzed by the RTC Coloc study, of which there were few, we did not apply their method, as it was too computationally intensive to apply to real trait data.

