## Supplementary material for "Modeling tissue co-regulation to estimate tissue-specific contributions to disease": TCSC_SupplementaryNote_Rev

**Supplementary Material for**
***Modeling tissue co-regulation to estimate tissue-specific contributions to disease***

**Supplementary Table Captions**

**Supplementary Table 1. Numerical results for robustness and power of TCSC regression in simulations (Figure 2).** Across six different eQTL sample sizes, we evaluate the causal and null bias in estimates of disease heritability explained by the *cis*-genetic component of gene expression in tissue $t'$ ($h^2_{ge(t')}$), the type I error, and the power of TCSC. The standard errors (SE) are computed as the standard deviation of measurements across simulations divided by the square root of the number of simulations, e.g. 1,000. Type I error is measured as the percentage of estimates of $h^2_{ge(t')}$ for non-causal tissues that were significantly positive for non-causal tissues at $p < 0.05$ for nominal significance or at 5% FDR across tissues. Power is measured as the percentage of estimates of $h^2_{ge(t')}$ for causal tissues that were significantly positive at $p < 0.05$ for nominal significance or at 5% FDR across tissues.

**Supplementary Table 2. Type I error and power of RTC Coloc, LDSC SEG, RolyPoly, and CoCoNet in simulations.** We implemented all methods as previously described and applied it to our TCSC simulation framework, such that the same eQTL effect sizes and co-regulation was used. We performed 1,000 simulations of LDSC SEG, RolyPoly, and CoCoNet and 100 simulations of RTC Coloc, due to the complexity and prohibitively large computation time of RTC Coloc.

**Supplementary Table 3. Numerical results for robustness and power of cross-trait TCSC in simulations (Figure 3).** Across six different eQTL sample sizes, we evaluate the causal and null bias on the estimate of tissue-specific contributions to covariance, the type I error, and the power of cross-trait TCSC. The standard errors (SE) are computed as the standard deviation of measurements across simulations divided by the square root of the number of simulations, e.g. 1,000. Type I error is measured as the percentage of estimates of $\omega_{ge(t')}$ for non-causal tissues that were significantly positive at $p < 0.05$ for nominal significance or at 5% FDR across tissues. Power is measured as the percentage of estimates of $\omega_{ge(t')}$ for causal tissues that were significantly positive at $p < 0.05$ for nominal significance or at 5% FDR across tissues.

**Supplementary Table 4. List of 78 diseases and complex traits analyzed in primary analyses.** We selected 78 diseases/traits, 33 of which are from UK Biobank, such that all summary statistics have SNP-heritability z-score > 6 and no pair of traits have a squared genetic correlation greater than 0.1 as well as substantial sample overlap. We report the SNP-heritability, standard error, z-score, GWAS sample size, trait nickname used to index traits in plots and tables, and the name of the most closely related trait analyzed in previous studies[1,2].

**Supplementary Table 5. Numerical results for tissue-specific contributions to disease and complex trait heritability (Figure 4).** For each significant tissue-trait pair identified by TCSC as reported in **Figure 4**, we report the value of $\pi_{t'}$, or the proportion of SNP-heritability explained by tissue-specific predicted gene expression, and the false discovery rate for each finding. We also report the result of an independent analysis for traits for which we could easily obtain independent GWAS summary statistics.

**Supplementary Table 6. Numerical results for tissue-specific contributions to disease and complex trait heritability for all tissues and diseases/traits analyzed.** For every tissue-trait pair analyzed by TCSC, across 39 tissues and 78 diseases/traits, we report the phenotypic variance explained by tissue-specific predicted gene expression, $h^2_{ge(t')}$, the jackknife standard error of this quantity, the nominal *P* value, the FDR calculated across tissues per-trait, the value of $\pi_{t'}$, and the standard error of $\pi_{t'}$. We estimated the standard error of this quantity using a genomic block jackknife. We note that no value of $h^2_{ge(all\ tissues)}$ exceeds the SNP-heritability for a given trait. The largest value of $h^2_{ge(all\ tissues)}$ is 0.68, which is for red blood cell distribution width.

**Supplementary Table 7. Median jackknife *P* values across traits for each tissue.** Here we report the median jackknife *P* value across traits for each tissue. For pairs of tissues with high genetic correlation, if the median jackknife *P* value is substantially different across traits, this means TCSC is systematically more likely to identify the tissue with lower median jackknife *P* value as a causal tissue relative to the other, and this might suggest an issue in quality of gene expression prediction models from the tissue with larger median jackknife *P* value.

**Supplementary Table 8. Numerical results for tissue-specific contributions to disease and complex trait heritability in secondary analysis of 23 tissues, removing tissues with small eQTL sample size.** To increase the power of TCSC to identify causal tissues, we removed tissues with eQTL sample size less than 320. As a result, we analyzed 23 tissues across 78 diseases/traits. We report the same quantities reported in **Supplementary Table 6**.

**Supplementary Table 9. Statistical significance of differences between TCSC estimates in primary and secondary analyses.** For every significant tissue-trait pair identified in the primary analysis (analysis of 39 tissues, **Figure 4**), we assessed if the value of $\pi_{t'}$ was significantly different than the value produced in the secondary analysis (analysis of 23 tissues, **ST8**). We used a genomic block jackknife to assess the difference and using a two-sided test, identified that no differences were significantly nonzero at 10% FDR.

**Supplementary Table 10. List of 41 brain diseases/traits analyzed in brain-specific analysis.** We performed a brain-specific TCSC analysis to exploit the diversity of brain tissues provided by GTEx (n = 13 brain tissues). We analyzed 41 brain diseases/traits using a similar trait selection procedure as was used to select the 78 diseases/traits previously analyzed. However, we first selected for diseases/traits that were behavioral or a known cognitive disorder and iteratively removed traits until no pair of traits had a squared genetic correlation greater than 0.25.

**Supplementary Table 11. List of 13 GTEx brain tissues analyzed in brain-specific analysis.** For the brain-specific TCSC analysis, we built gene expression prediction models using all European samples from each of 13 GTEx brain tissues, without subsampling or meta-analysis. Here we list the name and eQTL sample size of each GTEx brain tissue, e.g. tissue names beginning with "Brain_". For this analysis, we excluded several tissues relevant to the central nervous system, including pituitary ($N$ = 220) and tibial nerve ($N$ > 320).

**Supplementary Table 12. Numerical results for tissue-specific contributions to disease and complex trait heritability in brain-specific analysis.** For every brain tissue and brain trait analyzed in the brain-specific TCSC analysis, we report $\pi_{t'}$, its standard error, and false discovery rate.

**Supplementary Table 13. Numerical results for comparison of disease-critical tissues identified by RTC Coloc, LDSC-SEG and TCSC for 5 representative traits (Figure 5).** For every tissue-trait pair shown in **Figure 5**, we report the FDR and -$\log_{10}$FDR of the association statistic for each method (enrichment statistic for Ongen 2017 RTC Coloc, tau* S-LDSC statistic for Finucane 2018 LDSC-SEG, and $\pi_{t'}$ for TCSC). The seven traits are the ones having at least one significantly associated tissue across the three methods with the largest SNP-heritability $z$-score. The tissues reported here are the causal tissues for each of the five traits as well as the most genetically correlated tissue (using marginal eQTL effect sizes).

**Supplementary Table 14. Numerical results for comparison of disease-critical tissues identified by RTC Coloc, LDSC-SEG and TCSC for all 21 diseases/traits with causal tissue-trait associations identified by TCSC.** We report the FDR and -$\log_{10}$FDR of the association statistic for each method across all traits shown in **Figure 4** and each tissue with an association statistic with FDR < 5%.

**Supplementary Table 15. Numerical results for comparison of disease-critical tissues identified by RTC Coloc, LDSC-SEG and TCSC for all diseases/traits and tissues included in these comparisons.** We report the FDR and -$\log_{10}$FDR for every tissue-trait pair (39 tissues, 78 traits) across each of three compared methods. A value of NA indicates that the tissue-trait pair was not analyzed by the corresponding method.

**Supplementary Table 16. Numerical results for comparison of disease-critical tissues identified by RTC Coloc, LDSC-SEG and TCSC for brain-specific analysis.** We report the FDR and -$\log_{10}$FDR for the brain-specific analysis for each of 41 brain traits, 13 brain tissues, and 3 methods, restricting to tissues and traits with a TCSC finding at FDR < 10%, plotted in **SF29**. A value of NA indicates that the tissue-trait pair was not analyzed by the corresponding method.

**Supplementary Table 17. List of 262 pairs of diseases/traits analyzed by cross-trait TCSC.** We computed the genetic correlation between all pairs of 78 diseases/traits and selected those pairs with genetic correlation $P$ value < 0.05/3,003 pairs of traits, e.g. using a Bonferroni correction threshold. Here, we report the genetic correlation $z$-score of these pairs and the estimate of the covariance.

**Supplementary Table 18. Numerical results for tissue-specific contributions to the genetic covariance of two diseases/traits (Figure 6A).** For all tissue-trait covariance pairs identified by

TCSC at 10% FDR, we report the value of $\zeta_{t'}$, or the proportion of covariance explained by predicted gene expression in tissue $t'$ and the FDR.

**Supplementary Table 19. Numerical results for tissue-specific contributions to the genetic covariance of two diseases/traits for all tissues and disease/trait pairs analyzed.** For all tissue-trait covariance pairs analyzed by TCSC, we report the estimated tissue-specific covariance, its jackknife standard error, nominal *P* value, $\zeta_{t'}$ and corresponding standard error, FDR, and genome-wide covariance for the trait pair.

**Supplementary Table 20. List of tissue-trait covariance pairs and reported differences in tissue-specific contributions to genetic covariance vs. constituent trait heritability.** For every pair of traits implicated by **Figure 6** and for each of 38 tissues, we assess the difference between $\zeta_{t'}$ and $\pi_{t'}$ for each trait. We identified five tissue-trait covariance pairs for which the difference was significantly nonzero while the value of $\pi_{t'}$ was not significantly different than zero at a significance threshold of 5% FDR across tissues per-trait.

**Supplementary Table 21. Scenarios where TCSC has more power in the cross-trait analysis than in the single-trait analysis.** We used primary simulations and performed new simulations in which tissue-specific contributions to covariance where greater than tissue-specific contributions to heritability in order to report the percentage of simulations in which the causal tissue was detected in the cross-trait analysis but not detected in both of the single-trait analyses.

### Supplementary Note

#### *Secondary simulation analyses*

In **Supplementary Figure 5**, we varied the value of $h^2_{ge(t')}$ for causal tissues across different eQTL sample sizes. In panel A, we observe that at type I error is more consistent across different values of $h^2_{ge(t')}$ at smaller eQTL sample sizes. At larger eQTL sample sizes, smaller values of $h^2_{ge(t')}$ have the lowest error rates, with the smallest value not significantly different than 5%. In panel B, we observe that small values of $h^2_{ge(t')}$ result in low power, medium values of $h^2_{ge(t')}$ result in the greatest observed power, and (potentially unrealistically) large values of $h^2_{ge(t')}$ result in mediocre power. In panel C, estimates of $h^2_{ge(t')}$ are unbiased across sample sizes and different true values of $h^2_{ge(t')}$. In panel D, there is greater null bias for small eQTL sample sizes and larger values of $h^2_{ge(t')}$ in the causal tissue. These patterns are similar when varying the value of $\omega_{ge(t')}$ in **Supplementary Figure 6**.

In **Supplementary Figure 7**, we varied the number of causal tissues in the TCSC model. In panel A, the type I error tends to decrease with an increasing number of causal tissues. In panel B, the power to detect two or three causal tissues was significantly less than the power to detect a single causal tissue. In panel C, estimates of $h^2_{ge(t')}$ for causal tissues have anti-conservative bias for two or more tissues. In panel D, null bias decreases when there are multiple causal tissues. We observe similar patterns in **Supplementary Figure 8** when varying the number of causal tissues in cross-trait TCSC simulations.

In **Supplementary Figure 9**, we varied the number of non-causal tissues in the TCSC regression. In panel A, type I error decreased when increasing the number of non-causal tissues. In panel B, power was greatest for one or two tagging tissues, but decreased with every additional tagging tissue. In panel C, the estimate of $h^2_{ge(t')}$ for causal tissues had anti-conservative bias when there were fewer than 9 tagging tissues and unbiased when there were 9 tagging tissues. In panel D, null bias of $h^2_{ge(t')}$ for non-causal tissues is not significantly different than zero where there is only one tagging tissue; the most extreme case of anti-conservative null bias occurs at middle numbers of tagging tissues. We observed similar patterns for cross-trait TCSC in **Supplementary Figure 10**.

#### *Other findings of tissue-specific contributions to disease and complex trait heritability in primary TCSC analysis.*

WHRadjBMI (waist-hip-ratio conditional on body mass index) and subcutaneous adipose tissue ($\pi_{t'}$ = 0.10, s.e. = 0.037, $P = 2.4 \times 10^{-3}$). A previous study comparing subcutaneous adipose tissue to visceral adipose tissue found that the level of adiponectin, a hormone released by adipose tissue to regulate insulin, is specifically associated with subcutaneous adipose tissue and not visceral adipose tissue; and, adiponectin levels are significantly negatively correlated with waist-hip-ratio[3]. Furthermore, LDSC SEG found WHRadjBMI to be associated not only with subcutaneous adipose, but also with visceral adipose tissue. While RTC Coloc finds many WHR-associated tissues, it was able to distinguish subcutaneous adipose (FDR = $2.9 \times 10^{-4}$) from visceral adipose (FDR = 1).

HDL (high density lipoprotein) with subcutaneous adipose tissue ($\pi_{t'}$ = 0.159, s.e. = 0.054, $P$ = $1.5 \times 10^{-3}$) and whole blood ($\pi_{t'}$ = 0.098, s.e. = 0.034, $P$ = $1.8 \times 10^{-3}$). Previous work has implicated subcutaneous adipose tissue in mediating HDL levels, as this tissue stores cholesterol and expresses genes involved in cholesterol transport and HDL lipidation[4]. The relationship with whole blood is likely due to the role that red blood cells play in cholesterol transport, while being a large proportion of cells in whole blood samples[5]. Notably, TCSC did not identify liver as a causal tissue for HDL, and this might be due to the smaller eQTL sample size of liver which limits the power to detect this association.

BMI (body mass index) and brain cerebellum ($\pi_{t'}$ = 0.042, s.e. = 0.015, $P$ = $2.6 \times 10^{-3}$). While several studies have found that the central nervous system is enriched for genetic variation associated with BMI and obesity[6,2,7], the precise causal brain tissue is uncertain. Neither LDSC SEG nor RTC Coloc can distinguish between highly co-regulated brain tissues, such as the cerebellum and cortex. Previous studies have indicated that the brain cerebellum takes part in regulating feeding control (for example via connection to the hypothalamus) and therefore can have substantial impacts on obesity related traits and diseases[8]. Moreover, differential activity has been observed in the brain cerebellum in individuals experiencing hunger, thirst, or satiation[8]. Furthermore, a different study associated the brain cerebellum with endocrine homeostasis, suggesting that the cerebellum plays several important biological roles, rather than strictly motor control[9]. A more recent multi-omics approach identified that cerebellar nuclei in mice are activated when they are eating and even suggests a potential therapeutic target for the management of excessive eating behavioral traits[10].

Fecundity and brain cerebellum ($\pi_{t'}$ = 0.075, s.e. = 0.024, $P$ = $9.1 \times 10^{-4}$). This is consistent with the known relationship between fertility and energy metabolism, involving hormone secretion, which is largely regulated by the brain. However, previous studies have specifically linked fertility-related hormonal dysregulation to the hypothalamus and brainstem[11,12].

Total protein and fibroblasts ($\pi_{t'}$ = 0.079, s.e. = 0.025, $P$ = $7.0 \times 10^{-4}$) and whole blood ($\pi_{t'}$ = 0.081, s.e. = 0.027, $P$ = $1.5 \times 10^{-3}$). Fibroblasts are cells that play diverse roles across the tissues of the body, markedly producing protein complexes that constitute the extracellular matrices that define the structure of fibroblasts[13]. Serum protein is a quantity measured from whole blood, explaining the second relationship.

Cerebral cortex surface area and fibroblasts ($\pi_{t'}$ = 0.10, s.e. = 0.034, $P$ = $1.8 \times 10^{-3}$). Tissue surface areas are likely related to developmental processes governing body proportions. As stated in the main text, TCSC identified fibroblasts (and skeletal muscle) as causal tissues for height, the most commonly studied anthropometric phenotype, which suggests that fibroblasts, as a connective tissue, likely regulates the growth of different organs and tissues.

Lipid traits and liver: AST ($\pi_{t'}$ = 0.077, s.e. = 0.025, *P* = $9.2 \times 10^{-4}$), RBC width ($\pi_{t'}$ = 0.077, s.e. = 0.027, *P* = $1.9 \times 10^{-3}$), total cholesterol ($\pi_{t'}$ = 0.14, s.e. = 0.044, *P* = $5.3 \times 10^{-4}$), Bilirubin ($\pi_{t'}$ = 0.11, s.e. = 0.036, *P* = $1.0 \times 10^{-3}$). These causal tissue-trait pairs are reasonable as the liver is the production center of cholesterol and phospholipids.

Blood cell traits and whole blood: eosinophil count ($\pi_{t'}$ = 0.17, s.e. = 0.052, *P* = $6.5 \times 10^{-4}$), lymphocyte count ($\pi_{t'}$ = 0.22, s.e. = 0.053, *P* = $2.1 \times 10^{-5}$), monocyte count ($\pi_{t'}$ = 0.25, s.e. = 0.078, *P* = $7.5 \times 10^{-4}$). These causal tissue-trait pairs are reasonable as these different blood cell populations are present in whole blood.

MDD (Major depressive disorder) and whole blood ($\pi_{t'}$ = 0.068, s.e. = 0.022, *P* = $1.3 \times 10^{-3}$). This is consistent with reports of elevated immune system cytokines in MDD cases[14].

***Other findings of tissue-specific contributions to disease and complex trait heritability in secondary analysis of 23 tissues, removing tissues with small eQTL sample size***

BMI and tibial nerve: This is broadly consistent with the role of the central nervous system in BMI[15,6,16,17,2,18], although the precise causal relationship that might exist between tibial nerve and BMI is not straightforward.

Additional causal tissues for platelet count identified in this secondary analysis include brain cortex, esophagus muscularis, and fibroblasts. Regarding the brain, platelets are often found in blood vessels and are key participants in thrombosis, or the clotting of blood vessels[19,20]. Moreover, platelets have been linked to inflammation of death of neurons in the cortex and hippocampus[21]. Regarding the esophagus muscularis, high platelet counts are associated with greater severity of esophageal cancer, likely due to the angiogenic properties of platelets, e.g. creating new blood vessels[22]. Regarding fibroblasts, these cells are known to be recruited to sites of blood clots, caused by platelets, to remedy the clot[23]. Therefore, increased presence of fibroblasts likely reduces platelet activity in individuals with greater susceptibility to vascular clotting. While it is possible that platelet count may have a diverse tissue-specific genetic basis, this result could also be caused by an absent causal tissue or cell type that is co-regulated with these three newly detected tissues.

Sleep duration and breast tissue: melatonin is a hormone whose levels are considered protective for breast cancer risk[24]. Melatonin is also a common supplement taken to promote sleep. However, melatonin is produced in the brain, and therefore the causal relationship from breast tissue to sleep duration is unclear.

Height with fibroblasts and muscle skeletal tissue. Skin tissue has been shown to widely express growth factors, including embryonic growth factor which plays a key role in fetal development[25]. Fibroblasts are the predominant cell type of skin tissue. Skeletal muscle is one of the most likely causal tissues for anthropometric, or skeletal growth, traits such as height, consistent with previous genetic studies identifying enrichments of height-associated genetic variation near genes regulated in skeletal muscle[6,2], which includes colocalization with eQTLs regulating key growth factors such as IGFBP-3[26].

RBC count with fibroblasts and whole blood: Red blood cells and fibroblasts work together during tissue remodeling processes of extracellular matrices[27,28]. However, these studies suggest that red blood cells stimulate fibroblasts to secrete important tissue remodeling molecules, such as interleukin-8 and metalloproteinases. As a blood cell population, the causal relationship between whole blood and red blood cell count is expected.

Eosinophil count with fibroblasts and muscle skeletal tissue: Similar to the role of red blood cells in tissue remodeling described above, eosinophils also interact with fibroblasts in tissue remodeling and fibrosis, although typically in response to inflammation and allergy[29]. Eosinophils have previously been implicated in myopathy, or muscular disease[30], likely due to their recruitment in response to allergy, infection, or cancer.

Testosterone and muscle skeletal tissue: atrophy of skeletal muscle is associated with lower levels of testosterone, a hormone produced by the testes and understood to be regulated by brain tissues[31,32]. These studies suggest that there is a causal relationship of testosterone on muscle skeletal tissue, rather than the reverse relationship suggested by TCSC.

***Other findings of tissue-specific contributions to disease and complex trait heritability in brain-specific analysis***

Caudate volume and accumbens: In individuals with major depressive disorder, the basal ganglia, of which the nucleus accumbens is a component, has an attenuated response to positive stimuli compared to healthy controls; and, it has been observed that this associates with reduced caudate volume[33].

Anisotropy mode with accumbens and cerebellum: Mode of anisotropy reflects the organization of white matter fibers in the brain and is used to suggest abnormalities in brain connections[34]. Therefore, any brain tissue connected to white matter could be causal for morphological anisotropy mode; indeed the nucleus accumbens and cerebellum have connections to white matter[35,36].

Schizophrenia with brain frontal cortex, brain cerebellum, and brain caudate. The association with the frontal cortex is consistent with previous studies reporting differences in gray and white matter volumes in schizophrenia cases vs. controls within the prefrontal cortex[37,38]. Previous large-scale genetic studies identified enrichments of schizophrenia-associated variants in gene sets regulating excitatory and inhibitory neurons[2,39,40], but did not distinguish the origin of this enrichment among the cortex, hippocampus, and amygdala. The association with the cerebellum might be due to its large proportion of neurons, and is also consistent with previous reports of decreased blood flow within the cerebellum in schizophrenia patients[41]. The association with caudate is consistent with early studies reporting schizophrenia-like characteristics in patients with damaged caudate projections[42,43]. While TCSC often identifies one causal tissue for a given trait, the identification of three causal tissues for schizophrenia may reflect a diverse tissue-specific genetic basis for the disease, the absence of the true causal tissue or cell type and its co-regulation with analyzed tissues, or the common presence of the true causal cell type among each of the three tissues.

Bipolar disorder with caudate and cerebellum: This is consistent with previous work linking reduced cerebellar volume to anxiety-related disorders[44,45] and is similarly consistent with previous work associating reduced caudate volumes with bipolar disorder[46].

Reaction time and cerebellum: This is consistent with previous studies in patients and monkeys with reduced reaction time and cerebellar lesions[47].

Cerebral cortex width with frontal cortex and spinal cord: Intuitively, the frontal cortex has a causal effect on the tissue of the same name. While the connection between spinal cord and cerebral cortex is not as straightforward, the spinal cord and hypothalamus are connected via hypothalamic projections[48] and hypothalamic projections to the cerebral cortex are responsible for propagating autonomic signaling[49].

Starting age of smoking habit and frontal cortex: This is consistent with previous work reporting that development of the frontal cortex during adolescence is associated with behaviors and lifestyle choices, such as smoking[50].

Brainstem volume and spinal cord: This is consistent with the brainstem being the connection point of the brain to the spinal cord[51].

***Comparison of disease-critical tissues identified by RTC Coloc, LDSC-SEG and TCSC for brain-specific analysis***

In the brain-specific analysis, patterns of LDSC-SEG and RTC Coloc were striking. First, LDSC-SEG did not identify heritability enrichments in any brain tissues other than cerebellum and cortex, suggesting that these two tissues are the only disease relevant parts of the brain, although this is highly unlikely. For example, for four traits LDSC-SEG produced very similar enrichments for the frontal cortex and the cortex. TCSC attributed these associations to the brain cerebellum, and in the specific case of schizophrenia, also implicated the frontal cortex. Second, six of the ten brain traits, for which TCSC identified a causal tissue at 10% FDR, had no associated tissue according to LDSC-SEG; these traits coincided with traits not analyzed by the RTC Coloc study. We note that the RTC Coloc study did not analyze all GTEx tissues; brain amygdala, spinal cord, and substantia nigra were omitted from their study. Lastly, RTC Coloc found 8 of 8 tested tissues shown in **Supplementary Figure 29** to be associated with schizophrenia and four of 8 tested tissues to be associated with BMI, a superset of the tissues implicated by TCSC.

***Other significant findings of tissue-specific contributions to the genetic covariance of two diseases/traits***

Negative contribution of brain cortex to the genetic covariance of neuroticism and years of education ($\zeta_{t'}$ = -0.10, s.e. = 0.029, *P* = $2.1 \times 10^{-4}$). When certain personality traits underlie neuroticism, such as conscientiousness, neuroticism has been shown to be positively correlated with educational success[52]. The specific implication of the brain cortex, as opposed to other brain tissues, has not be reported previously in the literature.

Positive contribution of the brain spinal cord to the genetic covariance of type 2 diabetes (T2D) and vitamin D ($\zeta_{t'}$ = 0.17, s.e. = 0.052, *P* = $5.5 \times 10^{-4}$). Vitamin D is a known neurosteroid, which affects various brain functions including calcium signaling and cellular differentiation[53], and reduced vitamin D is a prominent risk factor for infectious diabetes as well as diabetes mellitus (a subset of which is T2D)[54], explaining the negative covariance identified by TCSC.

Negative contribution of breast tissue to the genetic covariance of white blood cell count and BMI ($\zeta_{t'}$ = -0.16, s.e. = 0.041, *P* = $4.1 \times 10^{-5}$). This observation is consistent with many previous studies reporting an association of elevated white blood cell counts with breast cancer, as these cells are a biomarker of inflammation and are predictive of other cancers and cardiovascular disease[55-57]. One of these studies investigated this relationship in the context of BMI and found that in premenopausal women, individuals with lower BMI and breast cancer had elevated white blood cell counts[57]. This direction of effect is consistent with TCSC's detection of tissue-specific negative covariance between white blood cell count and BMI, despite a genome-wide positive genetic correlation of these two traits.

Negative contribution of lung to the genetic covariance of age at first birth and intelligence ($\zeta_{t'}$ = -0.096, s.e. = 0.026, *P* = $1.2 \times 10^{-4}$). First, previous work has found that older age at first birth is associated with reduced risk of lung cancer involving regulation by steroid hormones; and while some studies consider age at first birth to be a causal protective factor, this relationship might be better explained by reverse causality[58-61]. Indeed, TCSC is not impacted by reverse causality as phenotype cannot influence gene expression-modifying genetic variation. Second, positive health outcomes, including lung function, are genetically associated with cognitive traits in GWAS, although the causal mechanisms are poorly understood[62-64]. However, the direction of effect estimated by TCSC is inconsistent with these findings, possibly suggesting distinct causal mechanisms of lung tissue on these traits.

Negative contribution of lung to the genetic covariance of intelligence and years of education ($\zeta_{t'}$ = -0.045, s.e. = 0.013, *P* = $2.2 \times 10^{-4}$). As stated above, we would expect lung genes with a positive effect on intelligence to have a consistent direction of effect on years of education. While this is not what TCSC concludes, this may suggest distinct causal mechanisms of lung tissue on these traits.

Negative contribution of pituitary to the genetic covariance of vitamin D and WHRadjBMI ($\zeta_{t'}$ = -0.19, s.e. = 0.057, *P* = $4.5 \times 10^{-4}$). Previous work has established a relationship between vitamin D and bone structure development, which is directly related to WHRadjBMI[65]. Other work has suggested that this may be due to the positive correlation between vitamin D levels and growth hormone levels, such as IGF-1[66]. Moreover, irregularities in pituitary development (specifically pituitary stalk interruption syndrome) are associated with reduced IGF-1, in which individuals also have reduced serum levels of vitamin D[67].

Contributions to the genetic covariance of eosinophil count and platelet count by lung($\zeta_{t'}$ = -0.20, s.e. = 0.068, $P$ = $1.5 \times 10^{-3}$), ovary ($\zeta_{t'}$ = -0.15, s.e. = 0.052, $P$ = $2.2 \times 10^{-3}$), skin ($\zeta_{t'}$ = 0.28, s.e. = 0.087, $P$ = $7.1 \times 10^{-4}$), and whole blood ($\zeta_{t'}$ = 0.30, s.e. = 0.105, $P$ = $2.2 \times 10^{-3}$). Eosinophils and platelets are highly co-regulated, with eosinophils secreting platelet-activating enzymes[68]. Therefore, it is expected that across multiple tissues, genes have pleiotropic effects on eosinophil count and platelet counts. It is also possible that such genes have a direct effect on eosinophil count and a secondary effect mediated be eosinophils on platelet count.

Negative contribution of vagina to the genetic covariance of testosterone and vitamin D ($\zeta_{t'}$ = -0.25, s.e. = 0.079, $P$ = $6.5 \times 10^{-4}$). Previous work has shown that vaginal tissue growth and differentiation were improved as a result of increased vitamin D levels[69]. Similarly, testosterone is a hormone that plays a key role in healthy vaginal function[70]. Since TCSC detected negative covariance for these two traits, it is likely that they are regulated by distinct sets of genes.

Negative contribution of whole blood to the genetic covariance of age at first birth and rheumatoid arthritis ($\zeta_{t'}$ = -0.17, s.e. = 0.053, $P$ = $6.6 \times 10^{-4}$). The association between rheumatoid arthritis (RA) and whole blood, which is comprised of many immune cell types, is logical. However, previous work has not reported an association between whole blood, or the immune system, and age at first birth. Moreover, it is not immediately clear why genes that increase risk for RA would also increase age at first birth. We hypothesize that the underlying mechanism pertains to age-related changes in an individual's immune system which might affect reproductive behavior later in life, as risk for RA and other autoimmune diseases increases.

Negative contribution of spleen to the genetic covariance of major depressive disorder (MDD) and BMI ($\zeta_{t'}$ = -0.29, s.e. = 0.039, $P$ = $7.3 \times 10^{-4}$). As discussed above, whole blood was detected as causal tissue for MDD likely due to the role of the cytokines in regulation of MDD; the spleen plays a key role in the immune system. It is widely known that obesity, or high BMI, is associated with irregularities in immune cell counts[71].

Negative contribution of coronary artery to the genetic covariance of years of education and menopause age ($\zeta_{t'}$ = -0.13, s.e. = 0.115, $P$ = $7.3 \times 10^{-4}$). This is consistent with previous work indicating that reduced coronary artery disease risk is associated with more years of education via a Mendelian randomization study[72]. Late menopause is considered a protective factor coronary artery disease[73]. This biological consistency would suggest a positive covariance, therefore we might conclude that distinct sets of genes regulate years of education and menopause age in coronary artery.

Positive contribution of aorta artery to the genetic covariance of mode of anisotropy and menopause age ($\zeta_{t'}$ = 0.27, s.e. = 0.084, $P$ = $7.8 \times 10^{-4}$). Previous work has associated calcification in the aorta with bone loss, specifically in postmenopausal women[74]. Mode of anisotropy, from brain MRI which is a measure of structural cellular organization, is not a well-studied complex trait and has a lack of literature evidence to support any role for aorta, and similarly for any other tissue.

Contributions to the genetic covariance of anorexia and insomnia by tibial nerve ($\zeta_{t'}$ = 0.20, s.e. = 0.063, *P* = $7.8 \times 10^{-4}$), testis ($\zeta_{t'}$ = 0.18, s.e. = 0.062, *P* = $1.7 \times 10^{-3}$), and whole blood ($\zeta_{t'}$ = -0.14, s.e. = 0.053, *P* = $3.9 \times 10^{-3}$). The explanation for tibial nerve can be found in the main text. The association with whole blood might be explained by previous work demonstrating the role of the immune system in anorexia[75] and insomnia[76]. The immune regulation impacting insomnia is specifically discussed in the context of the central nervous system, further supporting the association with the tibial nerve, a central nervous system tissue. Testis size has also been associated with sleep irregularities[77]. Separately, the reduced production of the androgen hormone in the testis, or hypogonadism, is a comorbidity of male anorexia[78].

Negative contribution of prostate to the genetic covariance of medication use and years of education ($\zeta_{t'}$ = -0.075, s.e. = 0.024, *P* = $7.8 \times 10^{-4}$). This is consistent with previous studies establishing a negative association between drug use and prostate health outcomes[79]. The positive covariance detected with years of education is not supported by any literature evidence and could be a false positive.

Positive contribution of muscle skeletal to the genetic covariance of brain accumbens volume and caudate volume ($\zeta_{t'}$ = 0.20, s.e. = 0.063, *P* = $8.0 \times 10^{-4}$). Previous work indicates that musculoskeletal tissue likely influences biological processes the brain via regulation of energy metabolism[80].

Negative contribution of muscle skeletal to the genetic covariance of total protein and WHRadjBMI ($\zeta_{t'}$ = -0.27, s.e. = 0.084, *P* = $8.0 \times 10^{-4}$). This is consistent with known regulation in musculoskeletal tissue influencing waist-hip-ratio[81]. Musculoskeletal tissue is also related to protein levels, as restricted protein intake leads to dysregulation and morphology of skeletal muscle[82].

Negative contribution of skin to the genetic covariance of height and FVC ($\zeta_{t'}$ = -0.47, s.e. = 0.155, *P* = $1.2 \times 10^{-3}$). Height and forced vital capacity (FVC) are genetically correlated as both are affected by body proportions and growth-regulating processes. Skin tissue has been shown to widely express growth factors, including embryonic growth factor which plays a key role in fetal development[25].

Negative contribution of lung to the genetic covariance of age at first birth and menopause age ($\zeta_{t'}$ = -0.20, s.e. = 0.067, *P* = $1.3 \times 10^{-3}$). This is consistent with a negative association between earlier menopause age and healthy pulmonary function[83,84]. As described above, older age at first birth is associated with improved lung cancer outcomes[85].

Negative contribution of adipose subcutaneous to the genetic covariance of bipolar disorder and major depressive disorder ($\zeta_{t'}$ = -0.18, s.e. = 0.051, *P* = $2.7 \times 10^{-4}$). This is consistent with an expanding body of literature supporting a bidirectional link between obesity and depression[86].

Negative contribution of the meta-tissue brain limbic to the genetic covariance of risk tolerance and schizophrenia ($\zeta_{t'}$ = -0.15, s.e. = 0.049, *P* = $8.5 \times 10^{-4}$). Risk taking (or impulsivity) has previously been linked to both schizophrenia and bipolar disorder[87].

***Other examples with significant differences in tissue-specific contributions to genetic covariance vs. constituent trait heritability***

Negative contribution of skin (sun exposed) to the genetic covariance of height and FVC: Skin does not explain a nonzero proportion of heritability for height or FVC; however, skin does explain a significant amount of positive covariance, although the genome-wide covariance of this trait pair is negative.

Negative contribution of breast to the genetic covariance of WBC count and BMI: Breast is not a causal tissue for either trait, although it does explain a significant amount of negative covariance between the two traits, although the genome-wide covariance of this trait pair is positive.

Negative contribution of brain cortex to the genetic covariance of years of education and neuroticism: Brain cortex is not a causal tissue for either trait, consistent with what we found in the brain-specific analysis. However, brain cortex explains a significant amount of positive covariance between the two traits, although the genome-wide covariance is negative.

Negative contribution of pituitary to the genetic covariance of vitamin D and WHRadjBMI: While pituitary does not explain a nonzero proportion of the heritability of either trait, it does explain a significant amount of positive covariance between the two traits, although the genome-wide covariance is negative.

### Supplementary Figures

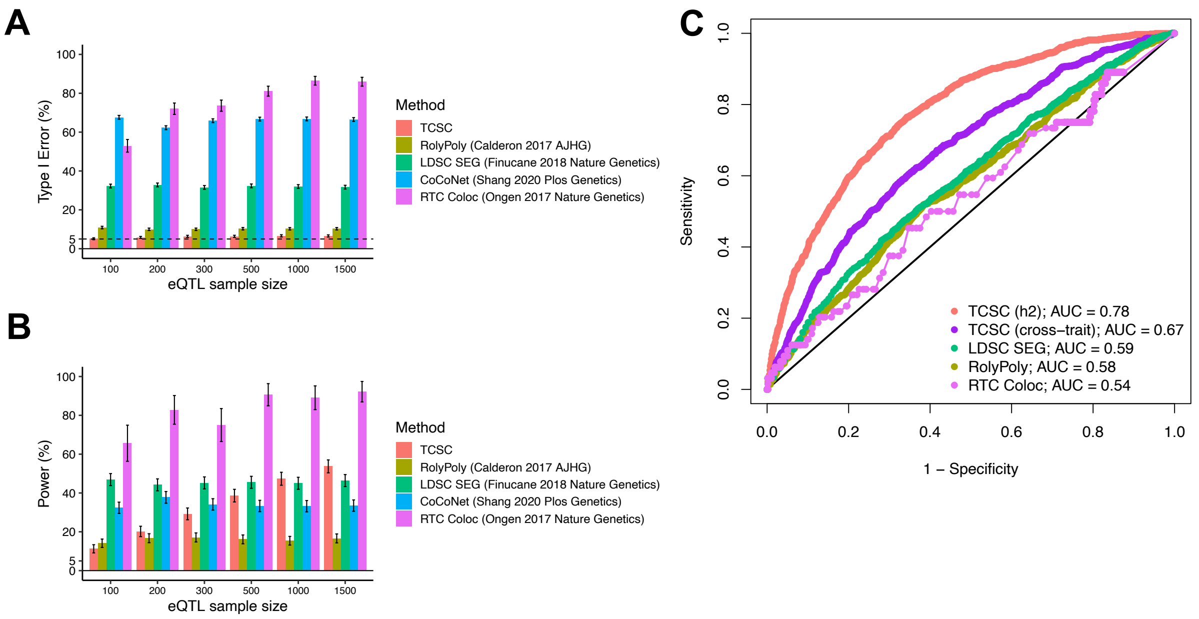


**Supplementary Figure 1. Comparison of tissue-trait association methods with TCSC in simulations.** (A) Percentage of estimates of $h^2_{ge(t')}$ for non-causal tissues that were significantly positive at $p < 0.05$, across 1,000 simulations per eQTL sample size for TCSC versus 1000 simulations per eQTL sample size for LDSC SEG, CoCoNet, and RolyPoly and 100 simulations per eQTL sample size for RTC Coloc due to the computationally intensive nature of the method. $h^2_{ge(t\ causal)}$ is set to 10% and GWAS sample size is set to 10,000. (B) Percentage of estimates of $h^2_{ge(t')}$ for causal tissues that were significantly positive at $p < 0.05$, across 1,000 simulations per eQTL sample size for TCSC versus 1000 simulations per eQTL sample size for LDSC SEG, CoCoNet, and RolyPoly and 100 simulations per eQTL sample size for RTC Coloc due to the computationally intensive nature of the method. Error bars in panels A and B represent 95% confidence intervals, computed using the standard error of the mean. (C) Receiver operating characteristic (ROC) curves for each method, including cross-trait TCSC, across 1000 uniformly spaced p-values between 0 and 1 used as the threshold to identify a causal tissue at a simulation eQTL sample size of 300, most closely matching real data analysis. We note that CoCoNet cannot be compared here because the method identifies the single most likely causal tissue using maximum likelihood estimation rather than via p-value. Numerical results are provided in **Supplementary Tables 1 and 2**.

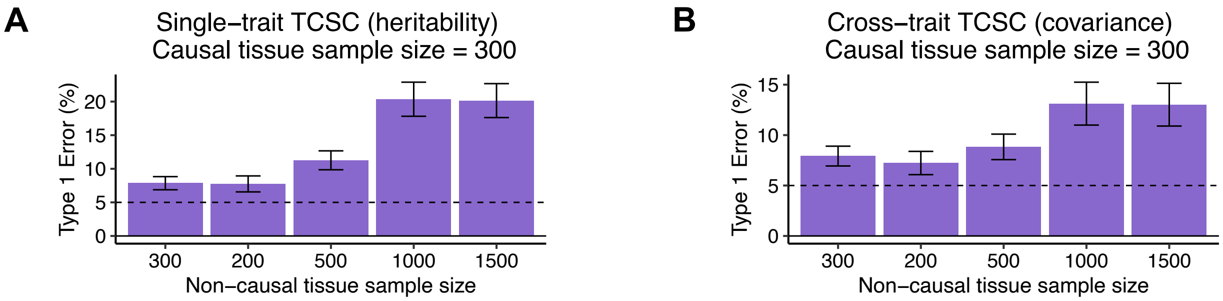


**Supplementary Figure 2. Type I error of TCSC regression in simulations with large variations in eQTL sample size of non-causal tissues.** We performed 1,000 simulations in which each simulation had one causal tissue (gene expression sample size = 300 individuals) and nine non-causal tissues with the following sample sizes: 300, 200, 300, 500, 1000, 1500, 200, 300, 500. (A) We report the false positive rate for non-causal tissues as $h^2_{ge(t')}$ > 0 at *p < 0.05* which is not well-controlled, demonstrating the need for comparable gene expression sample sizes across tissues in TCSC. (B) We report the false positive rate for non-causal tissues as $\omega_{ge(t')}$ > 0 at *p < 0.05* which is not well-controlled, demonstrating the need for comparable gene expression sample sizes across tissues in TCSC. Error bars represent 95% confidence intervals, computed using the standard error of the mean.

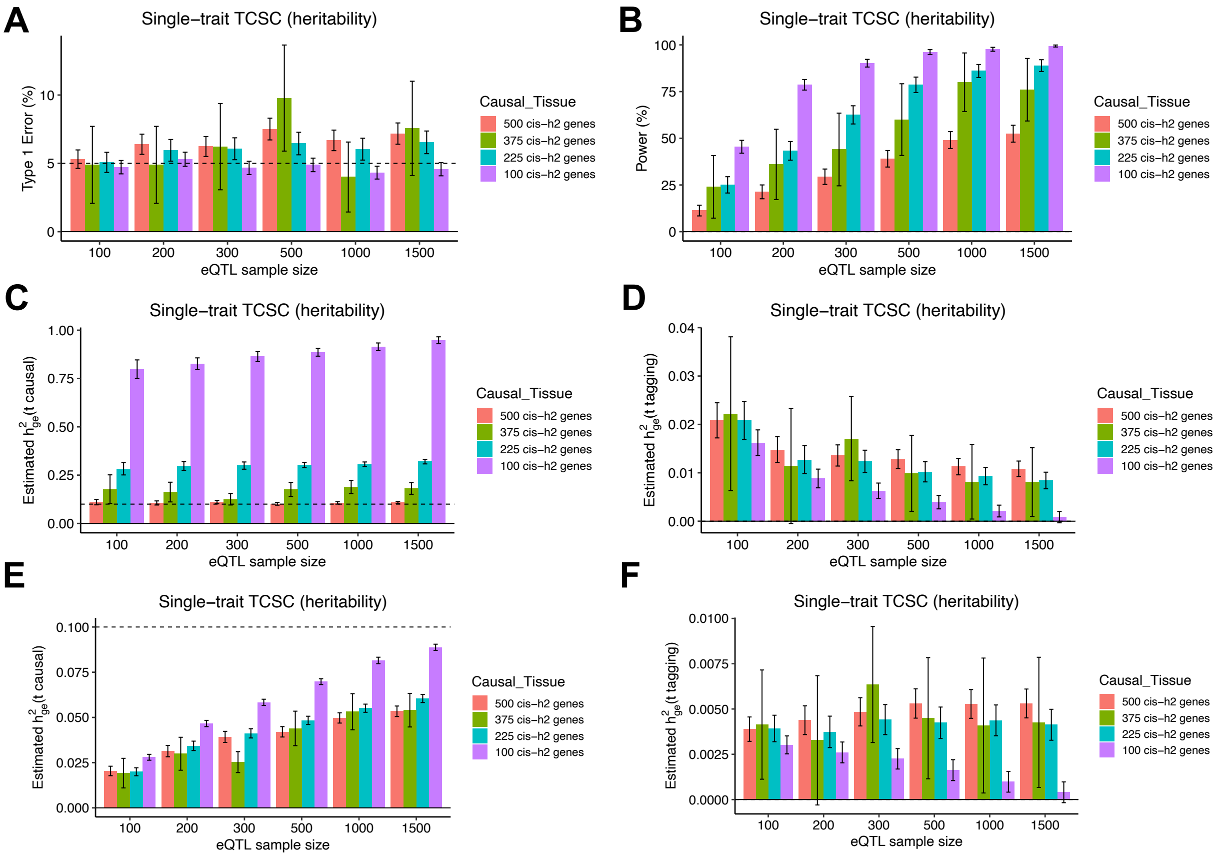


**Supplementary Figure 3. Robustness and power of TCSC regression in simulations when the causal tissue has fewer cis-heritable genes than tagging tissues.** (A) Type I error across 1,000 simulations per scenario of changing the number of expressed genes in the causal tissue. False positive event is defined as $h^2_{ge(t')} > 0$ for non-causal tissues at *p < 0.05*. (B) Power to detect the causal tissue as a proportion of 1,000 simulations per scenario. A true positive event is defined as $h^2_{ge(t')} > 0$ for causal tissues at *p < 0.05*. (C) Bias on causal estimates of $h^2_{ge(t')}$ for different scenarios. Dashed line indicates true value of $h^2_{ge(t')}$. The value of $G_{t'}$ is set to the total number of unique *cis*-heritable genes across all tissues. (D) Bias on non-causal estimates of $h^2_{ge(t')}$ for different scenarios. The value of $G_{t'}$ is set to the total number of unique *cis*-heritable genes across all tissues. (E) Bias on causal estimates of $h^2_{ge(t')}$ for different scenarios. Dashed line indicates true value of $h^2_{ge(t')}$. $G_{t'}$ is set to the number of significantly *cis*-heritable genes detected in each tissue. (F) Bias on non-causal estimates of $h^2_{ge(t')}$ for different scenarios. $G_{t'}$ is set to the number of significantly *cis*-heritable genes detected in each tissue. For all panels, error bars represent 95% confidence intervals, computed using the standard error of the mean.

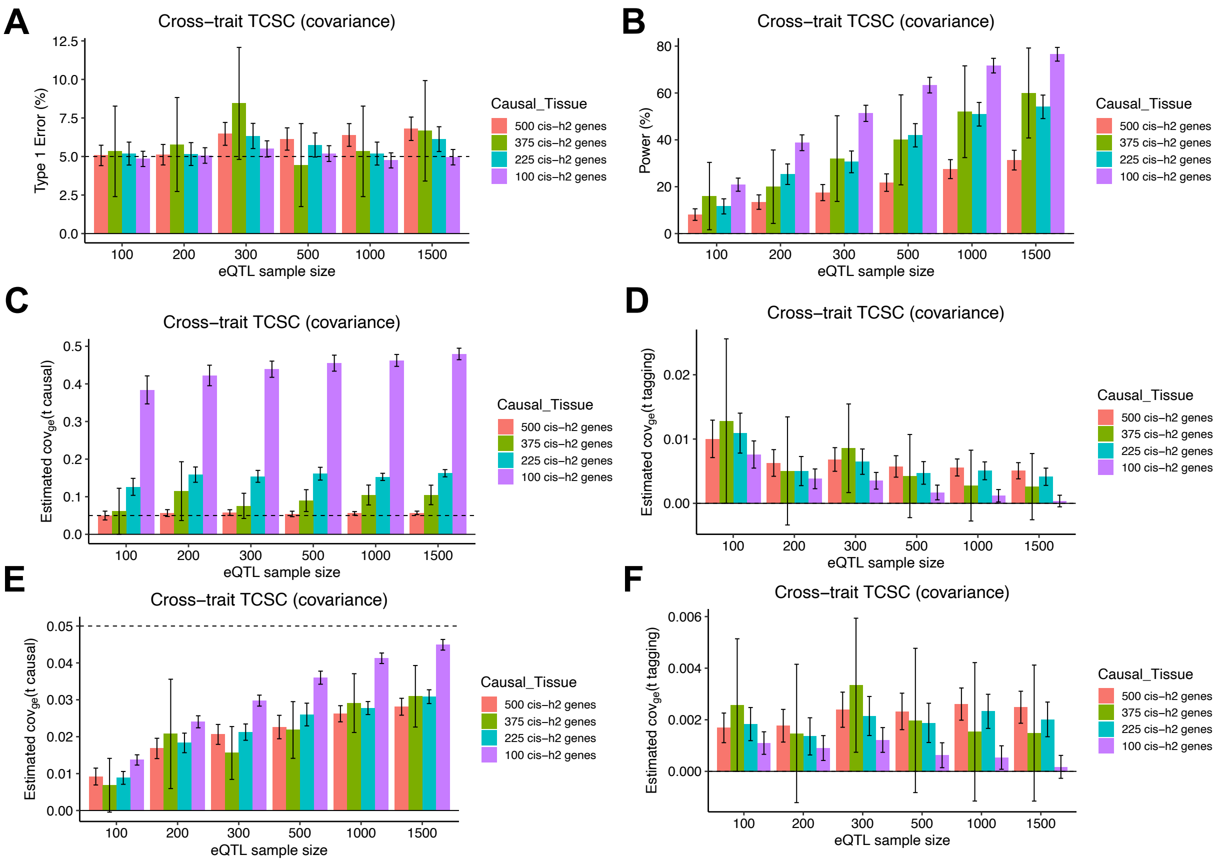


**Supplementary Figure 4. Robustness and power of TCSC regression in simulations when the causal tissue has fewer cis-heritable genes than tagging tissues.** (A) Type I error across 1,000 simulations per scenario of changing the number of expressed genes in the causal tissue. False positive event is defined as $\omega_{ge(t')} > 0$ for non-causal tissues at $p < 0.05$. (B) Power to detect the causal tissue as a proportion of 1,000 simulations per scenario. A true positive event is defined as $\omega_{ge(t')} > 0$ for causal tissues at $p < 0.05$. (C) Bias on causal estimates of $h^2_{ge(t')}$ for different scenarios. Dashed line indicates true value of $\omega_{ge(t')}$. The value of $G_{t'}$ is set to the total number of unique *cis*-heritable genes across all tissues. (D) Bias on non-causal estimates of $\omega_{ge(t')}$ for different scenarios. The value of $G_{t'}$ is set to the total number of unique *cis*-heritable genes across all tissues. (E) Bias on causal estimates of $\omega_{ge(t')}$ for different scenarios. Dashed line indicates true value of $\omega_{ge(t')}$. $G_{t'}$ is set to the number of significantly *cis*-heritable genes detected in each tissue. (F) Bias on non-causal estimates of $\omega_{ge(t')}$ for different scenarios. $G_{t'}$ is set to the number of significantly *cis*-heritable genes detected in each tissue. For all panels, error bars represent 95% confidence intervals, computed using the standard error of the mean.

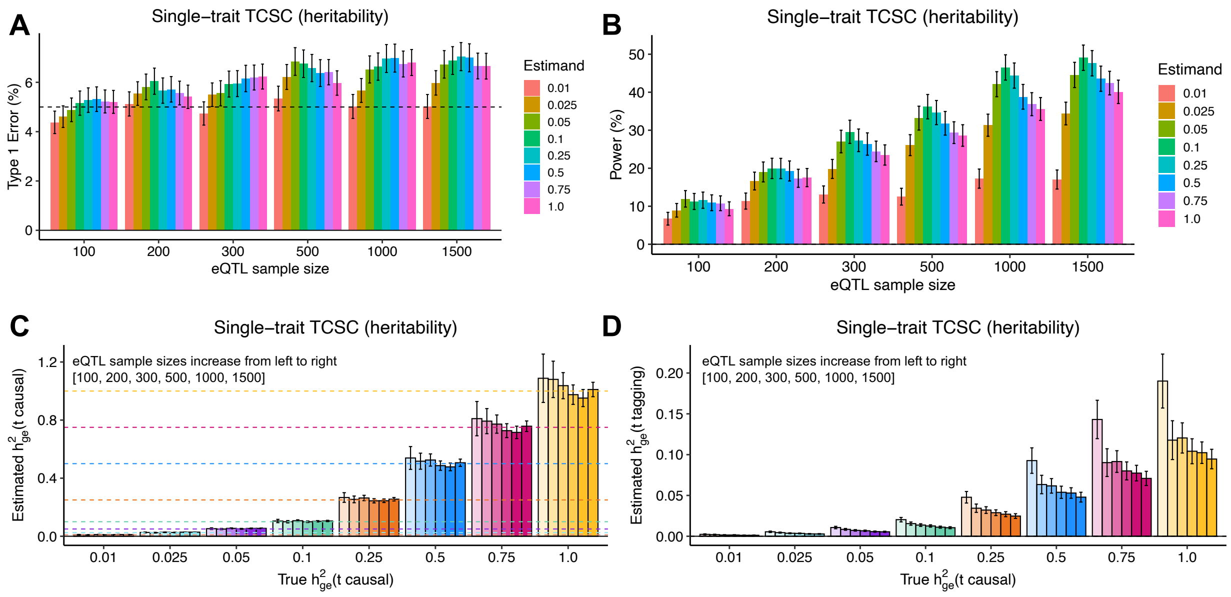


**Supplementary Figure 5. Robustness and power of TCSC regression in simulations with different values of $\boldsymbol{h^2_{ge(t')}}$.** (A) Type I error across 1,000 simulations per true value of $h^2_{ge(t')}$ in the causal tissue. False positive event is defined as $h^2_{ge(t')}$ > 0 for non-causal tissues at *p < 0.05*. (B) Power to detect the causal tissue as a proportion of 1,000 simulations per true value of $h^2_{ge(t')}$ in the causal tissue. A true positive event is defined as $h^2_{ge(t')}$ > 0 for causal tissues at *p < 0.05*. (C) Bias on causal estimates of $h^2_{ge(t')}$ for different true values of the causal tissue $h^2_{ge(t')}$. Dashed lines indicate true values of $h^2_{ge(t')}$. (D) Bias on non-causal estimates of $h^2_{ge(t')}$ for different true values of the causal tissue $h^2_{ge(t')}$. Error bars represent 95% confidence intervals, computed using the standard error of the mean. The value of $G_{t'}$ is set to the total number of unique *cis*-heritable genes across all tissues.

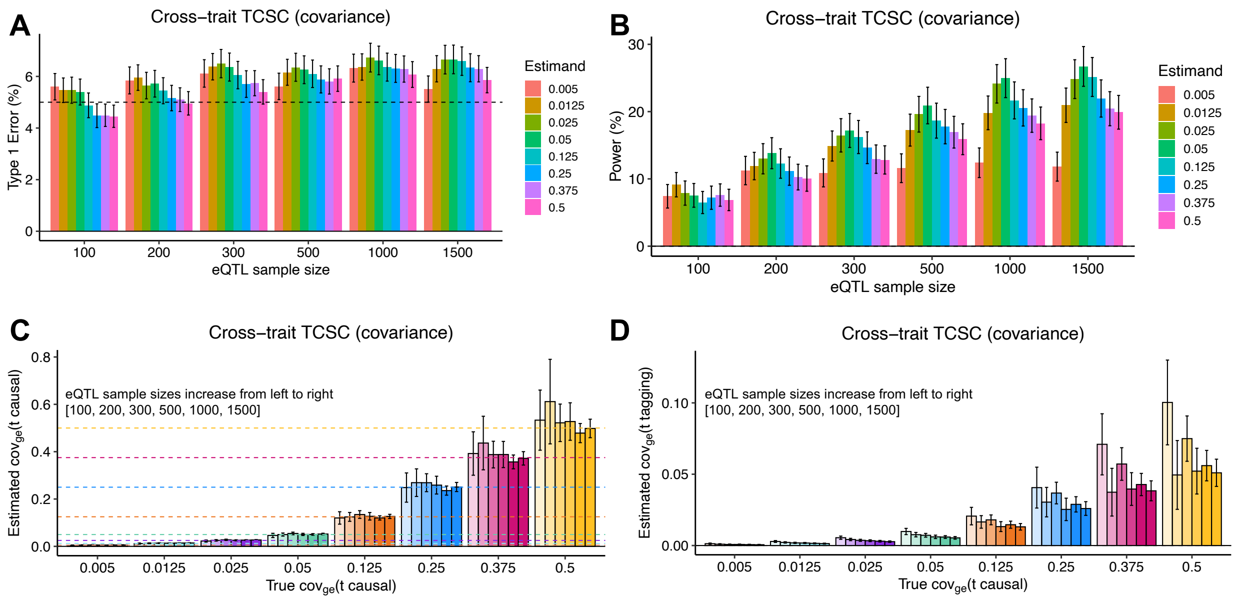


**Supplementary Figure 6. Robustness and power of cross-trait TCSC regression in simulations with different values of $\boldsymbol{\omega_{ge(t)}}$.** (A) Type I error across 1,000 simulations per true value of $\omega_{ge(t')}$ in the causal tissue. False positive event is defined as $\omega_{ge(t')} > 0$ for non-causal tissues at *p < 0.05*. (B) Power to detect the causal tissue as a proportion of 1,000 simulations per true value of $\omega_{ge(t)}$ in the causal tissue. A true positive event is defined as $\omega_{ge(t')} > 0$ for causal tissues at *p < 0.05*. (C) Bias on causal estimates of $\omega_{ge(t')}$ for different true values of the causal tissue $\omega_{ge(t')}$. Dashed lines indicate true values of $\omega_{ge(t')}$. (D) Bias on non-causal estimates of $\omega_{ge(t')}$ for different true values of the causal tissue $\omega_{ge(t')}$. Error bars represent 95% confidence intervals, computed using the standard error of the mean. The value of $G_{t'}$ is set to the total number of unique *cis*-heritable genes across all tissues.

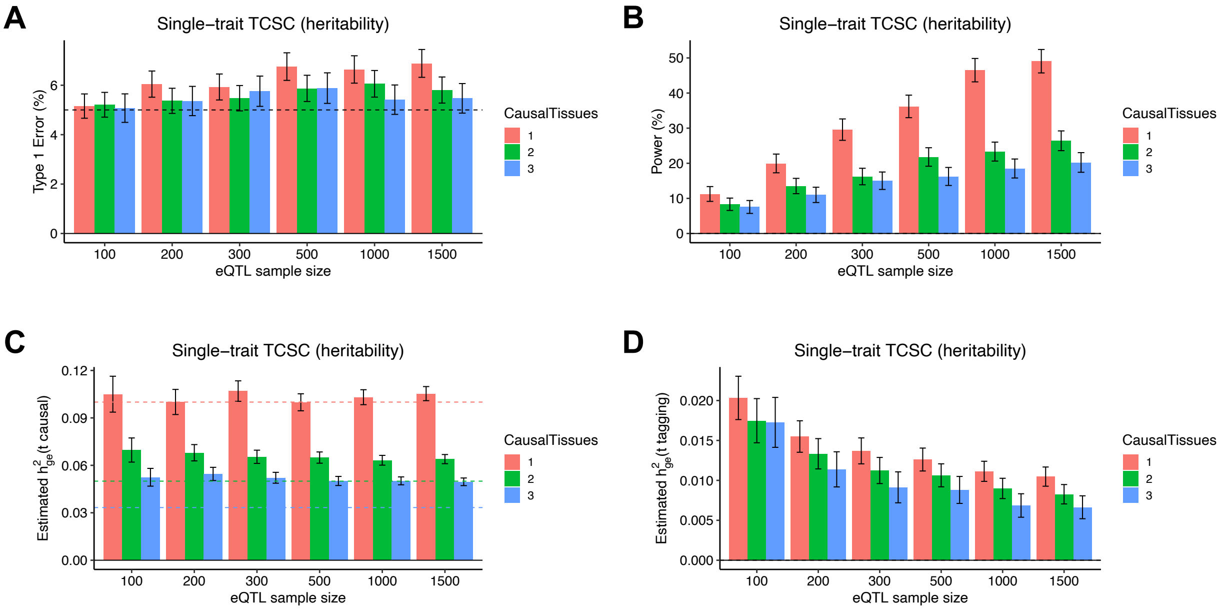


**Supplementary Figure 7. Robustness and power of TCSC regression in simulations with different numbers of causal tissues.** (A) Type I error across 1,000 simulations for each different causal tissue architecture. A single causal tissue (pink) represents the primary simulation analysis. Other architectures include two causal tissues (green) and three causal tissues (blue). False positive event defined as $h^2_{ge(t')}$ > 0 for non-causal tissues at *p < 0.05*. (B) Power to detect the causal tissue as a proportion of 1,000 simulations, in which $h^2_{ge(t')}$ > 0 for causal tissues at *p < 0.05*. (C) Bias on estimates of $h^2_{ge(t')}$ in causal tissues for different numbers of causal tissues in the model. The dashed line indicates that the true value of $h^2_{ge(t')}$ = 0.1. (D) Bias on estimates of $h^2_{ge(t')}$ in non-causal tissues for different numbers of causal tissues in the model. Error bars represent 95% confidence intervals, computed using the standard error of the mean. The value of $G_{t'}$ is set to the total number of unique *cis*-heritable genes across all tissues.

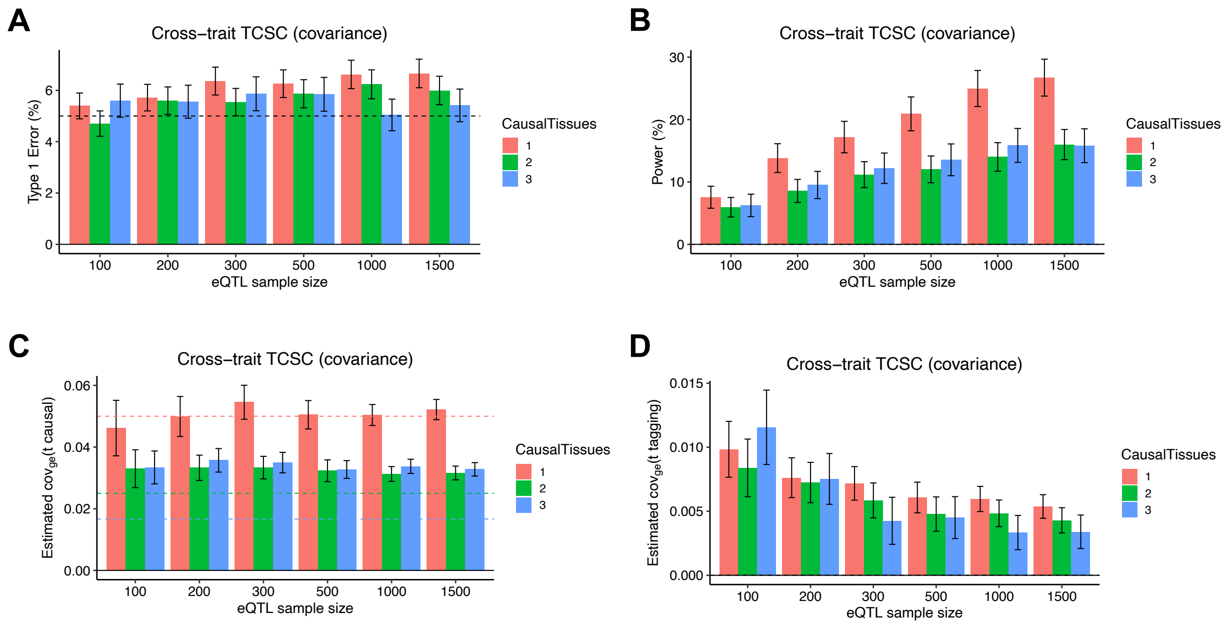


**Supplementary Figure 8. Robustness and power of cross-trait TCSC regression in simulations with different numbers of causal tissues.** (A) Type I error across 1,000 simulations for each different causal tissue architecture. A single causal tissue (pink) represents the primary simulation analysis. Other architectures include two causal tissues (green) and three causal tissues (purple). False positive event defined as $\omega_{ge(t')} > 0$ for non-causal tissues at $p < 0.05$. (B) Power to detect the causal tissue as a proportion of 1,000 simulations, in which $\omega_{ge(t')} > 0$ for non-causal tissues at $p < 0.05$. (C) Bias on estimates of $\omega_{ge(t')}$ in causal tissues for different numbers of causal tissues in the model. The dashed line indicates that the true value of $\omega_{ge(t')} = 0.05$. (D) Bias on estimates of $\omega_{ge(t')}$ in non-causal tissues for different numbers of causal tissues in the model. Error bars represent 95% confidence intervals, computed using the standard error of the mean. The value of $G_{t'}$ is set to the total number of unique *cis*-heritable genes across all tissues.

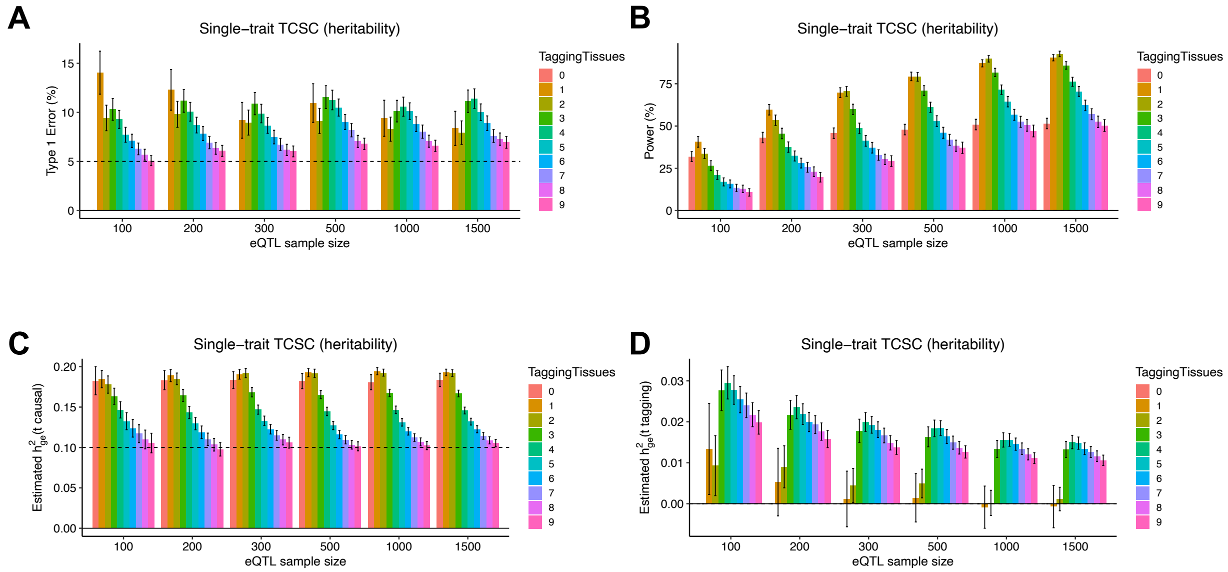


**Supplementary Figure 9. Robustness and power of TCSC regression in simulations with different numbers of non-causal tissues.** (A) Type I error across 1,000 simulations involving a variable number of non-causal tissues in the presence of a single causal tissue. False positive event defined as $h^2_{ge(t')}$ > 0 for non-causal tissues at *p < 0.05*. Note, when there are 0 tagging tissues, there is no measurement of type I error. (B) Power to detect the causal tissue as a proportion of 1,000 simulations in which $h^2_{ge(t')}$ > 0 for causal tissues at *p < 0.05*. (C) Bias on estimates of $h^2_{ge(t')}$ for the causal tissue, while changing the number of non-causal tissues in the model. The dashed line indicates that the true value of $h^2_{ge(t')}$. (D) Bias on estimates of $h^2_{ge(t')}$ for non-causal tissues, while changing the number of non-causal tissues in the model. Error bars represent 95% confidence intervals, computed using the standard error of the mean. The value of $G_{t'}$ is set to the total number of unique *cis*-heritable genes across all tissues.

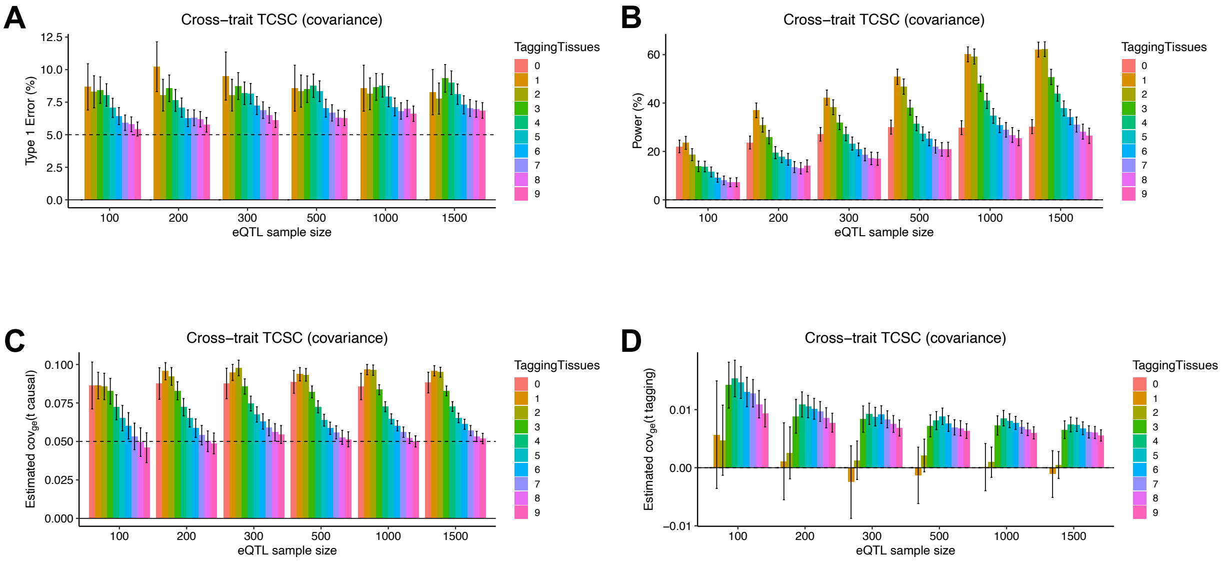


**Supplementary Figure 10. Robustness and power of cross-trait TCSC regression in simulations with different numbers of non-causal tissues.** (A) Type I error across 1,000 simulations involving a variable number of non-causal tissues in the presence of a single causal tissue. False positive event defined as $\omega_{ge(t')} > 0$ for non-causal tissues at $p < 0.05$. Note, when there are 0 tagging tissues, there is no measurement of type I error. (B) Power to detect the causal tissue as a proportion of 1,000 simulations, in which $\omega_{ge(t')} > 0$ for causal tissues at $p < 0.05$. (C) Bias on estimates of $\omega_{ge(t')}$ for the causal tissue, while changing the number of non-causal tissues in the model. The dashed line indicates that the true value of $\omega_{ge(t')}$ . (D) Bias on estimates of $\omega_{ge(t')}$ for non-causal tissues, while changing the number of non-causal tissues in the model. Error bars represent 95% confidence intervals, computed using the standard error of the mean. The value of $G_{t'}$ is set to the total number of unique *cis*-heritable genes across all tissues.

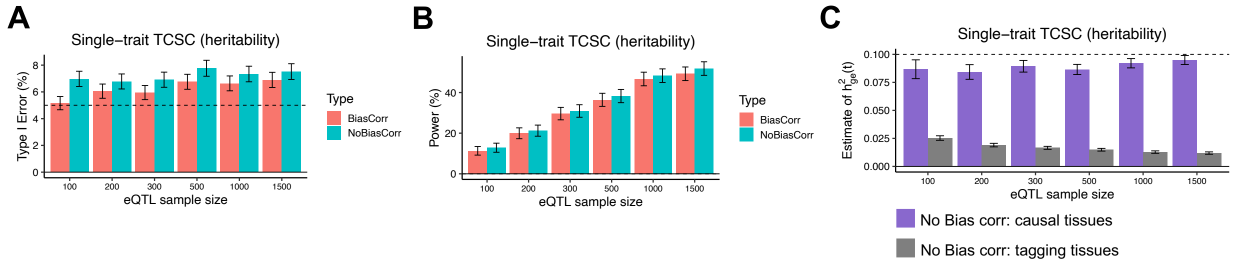


**Supplementary Figure 11. Robustness and power of TCSC regression with or without correction for bias in tissue co-regulation scores in simulations.** (A) Type I error across 1,000 simulations involving non-causal tissues for each of two scenarios: (1) “BiasCorr”: tissue co-regulation scores estimated using bias correction as in primary simulations (pink) vs (2) “NoBiasCorr”: tissue co-regulation scores estimated without bias correction (green). False positive event defined as $h^2_{ge(t')}$ > 0 for non-causal tissues at *p < 0.05*. (B) Power to detect the causal tissue as a proportion of 1,000 simulations, in which $h^2_{ge(t')}$ > 0 for causal tissues at *p < 0.05*. (C) Bias on estimates of causal and non-causal $h^2_{ge(t')}$ whose true values are 0.1 (purple bars) and 0 (gray bars), respectively, in the scenario of using no bias correction on tissue co-regulation scores. Error bars represent 95% confidence intervals, computed using the standard error of the mean. The value of $G_{t'}$ is set to the total number of unique *cis*-heritable genes across all tissues.

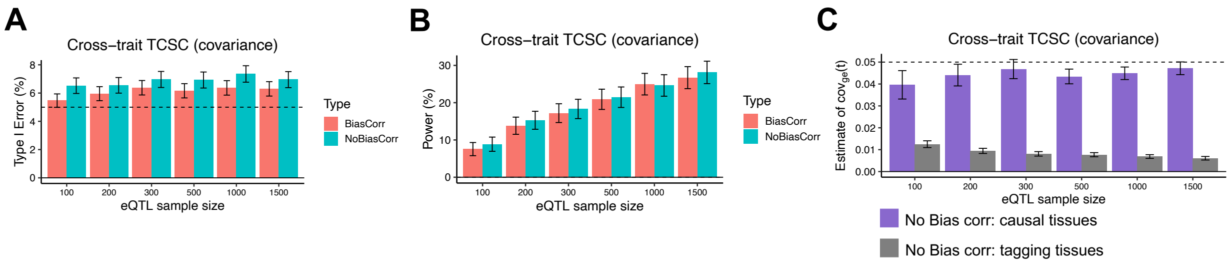


**Supplementary Figure 12. Robustness and power of cross-trait TCSC regression with or without correction for bias in tissue co-regulation scores in simulations.** (A) Type I error across 1,000 simulations involving non-causal tissues for each of two scenarios: (1) “BiasCorr”: tissue co-regulation scores estimated using bias correction as in primary simulations (pink) vs (2) “NoBiasCorr”: tissue co-regulation scores estimated without bias correction (green). False positive event defined as $\omega_{ge(t')}$ > 0 for non-causal tissues at *p < 0.05*. (B) Power to detect the causal tissue as a proportion of 1,000 simulations, in which $\omega_{ge(t')}$ > 0 for causal tissues at *p < 0.05*. (C) Bias on estimates of causal and non-causal $\omega_{ge(t')}$ whose true values are 0.05 (purple bars) and 0 (gray bars), respectively, in the scenario of using no bias correction on tissue co-regulation scores. Error bars represent 95% confidence intervals, computed using the standard error of the mean. The value of $G_{t'}$ is set to the total number of unique *cis*-heritable genes across all tissues.

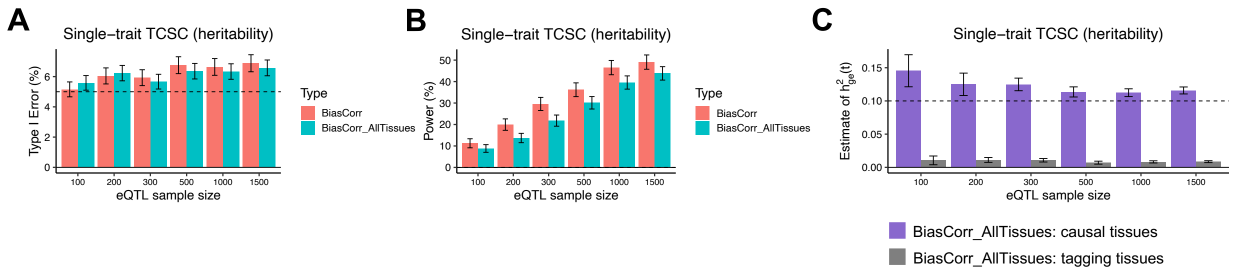


**Supplementary Figure 13. Robustness and power of single-trait TCSC regression without (default) or with bias correction applied to all pairs of tissues in simulations.** (A) Type I error across 1,000 simulations involving non-causal tissues for each of two scenarios: (1) “BiasCorr”: tissue co-regulation scores estimated using bias correction as in primary simulations, e.g. when t = t’ (pink) vs (2) “BiasCorr_AllTissues”: tissue co-regulation scores estimated using bias correction applied to all correlations of predicted gene expression (green). False positive event defined as $h^2_{ge(t')}$> 0 for non-causal tissues at *p < 0.05*. (B) Power to detect the causal tissue as a proportion of 1,000 simulations, in which $h^2_{ge(t')}$> 0 for causal tissues at *p < 0.05*. (C) Bias on estimates of causal and non-causal $h^2_{ge(t')}$whose true values are 0.1 (purple bars) and 0 (gray bars), respectively, in the scenario of using bias correction applied to all correlations of predicted gene expression in co-regulation scores. Error bars represent 95% confidence intervals, computed using the standard error of the mean. The value of $G_t$, is set to the total number of unique *cis*-heritable genes across all tissues.

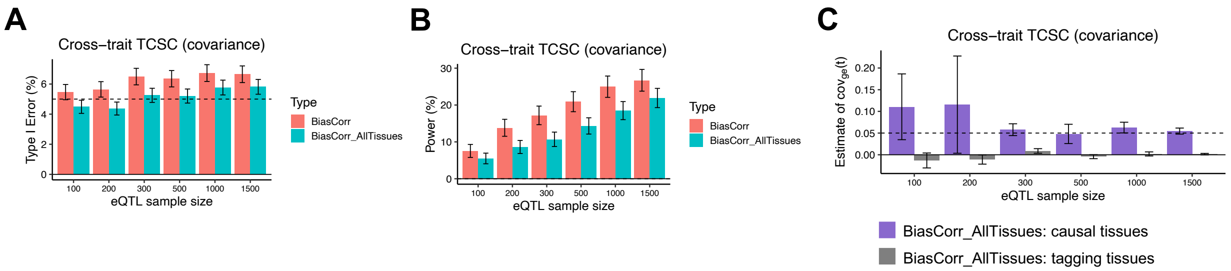


**Supplementary Figure 14. Robustness and power of cross-trait TCSC regression without (default) or with bias correction applied to all pairs of tissues in simulations.** (A) Type I error across 1,000 simulations involving non-causal tissues for each of two scenarios: (1) “BiasCorr”: tissue co-regulation scores estimated using bias correction as in primary simulations, e.g. when t = t’ (pink) vs (2) “BiasCorr_AllTissues”: tissue co-regulation scores estimated using bias correction applied to all correlations of predicted gene expression (green). False positive event defined as $\omega_{ge(t')} > 0$ for non-causal tissues at $p < 0.05$. (B) Power to detect the causal tissue as a proportion of 1,000 simulations, in which $\omega_{ge(t')} > 0$ for causal tissues at $p < 0.05$. (C) Bias on estimates of causal and non-causal $\omega_{ge(t')}$ whose true values are 0.05 (purple bars) and 0 (gray bars), respectively, in the scenario of using bias correction applied to all correlations of predicted gene expression in co-regulation scores. Error bars represent 95% confidence intervals, computed using the standard error of the mean. The value of $G_{t'}$ is set to the total number of unique *cis*-heritable genes across all tissues.

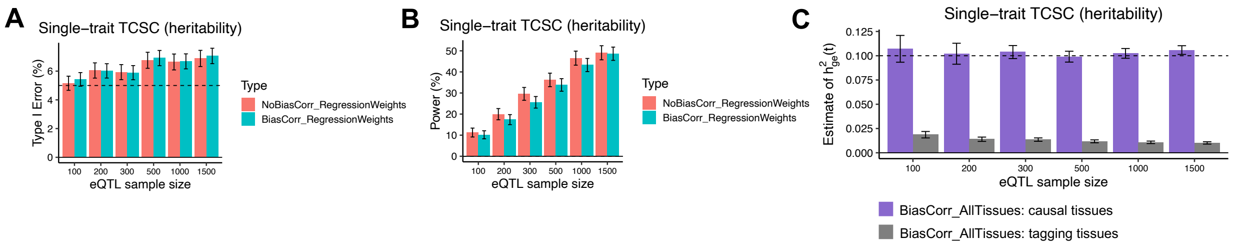


**Supplementary Figure 15. Robustness and power of TCSC regression without (default) or with bias correction for tissue co-regulation scores used to calculate regression weights in simulations.** (A) Type I error across 1,000 simulations involving non-causal tissues for each of two scenarios: (1) "NoBiasCorr_RegressionWeights": regression weights calculated using uncorrected tissue co-regulation scores as in primary simulations (pink) vs (2) "BiasCorr_RegressionWeights": regression weights calculated using bias-corrected tissue co-regulation scores (green). False positive event defined as $h^2_{ge(t')}$ > 0 for non-causal tissues at *p < 0.05*. (B) Power to detect the causal tissue as a proportion of 1,000 simulations, in which $h^2_{ge(t')}$ > 0 for causal tissues at *p < 0.05*. (C) Bias on estimates of causal and non-causal $h^2_{ge(t')}$ whose true values are 0.1 (purple bars) and 0 (gray bars), respectively, in the scenario of calculating regression weights using bias-corrected tissue co-regulation scores. Error bars represent 95% confidence intervals, computed using the standard error of the mean. The value of $G_{t'}$ is set to the total number of unique *cis*-heritable genes across all tissues. There is no methodological need to apply bias correction to the calculation of regression weights, since regression weights are used to maximize signal to noise rather than avoid bias in estimates of $h^2_{ge(t')}$ .

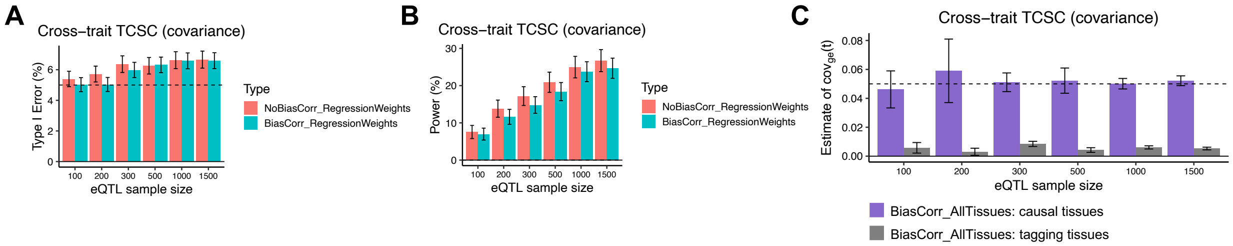


**Supplementary Figure 16. Robustness and power of cross-trait TCSC regression with or without correction for bias in tissue co-regulation scores in simulations.** (A) Type I error across 1,000 simulations involving non-causal tissues for each of two scenarios: (1) "NoBiasCorr_RegressionWeights": regression weights calculated using uncorrected tissue co-regulation scores as in primary simulations (pink) vs (2) "BiasCorr_RegressionWeights": regression weights calculated using bias-corrected tissue co-regulation scores (green). False positive event defined as $\omega_{ge(t')}$ > 0 for non-causal tissues at *p < 0.05*. (B) Power to detect the causal tissue as a proportion of 1,000 simulations, in which $\omega_{ge(t')}$ > 0 for causal tissues at *p < 0.05*. (C) Bias on estimates of causal and non-causal $\omega_{ge(t')}$ whose true values are 0.05 (purple bars) and 0 (gray bars), respectively, in the scenario of calculating regression weights using bias-corrected tissue co-regulation scores. Error bars represent 95% confidence intervals, computed using the standard error of the mean. The value of $G_{t'}$ is set to the total number of unique *cis*-heritable genes across all tissues. There is no methodological need to apply bias correction to the calculation of regression weights, since regression weights are used to maximize signal to noise rather than avoid bias in estimates of $\omega_{ge(t')}$ .

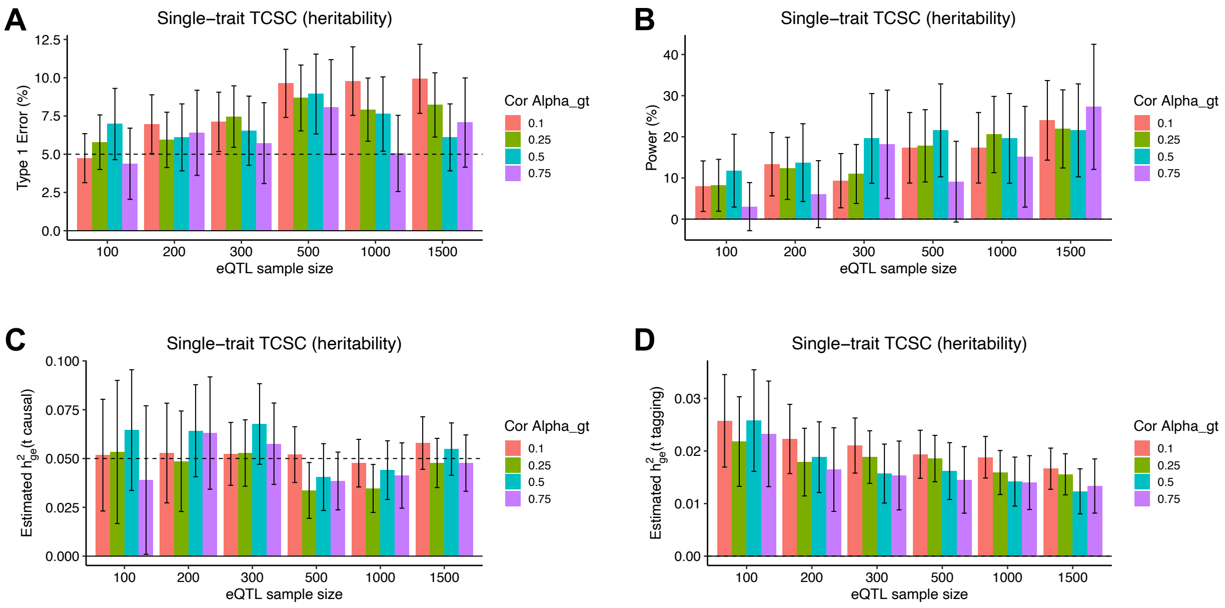


**Supplementary Figure 17. Robustness of cross-trait TCSC regression in simulations with two causal tissues with varying levels of correlated gene expression-trait effects.** To demonstrate how TCSC would behave when there are two causal tissues with correlation gene-trait effects ($\alpha_{gt}$), each tissue of which contributes 5% heritability to the trait. This is a model violation where TCSC assumes that gene expression-trait effects are i.i.d. (A) Type I error across 1,000 simulations with a varying correlation between the $\alpha_{gt}$ of each causal tissue. False positive event defined as $h^2_{ge(t\prime)}$ > 0 for non-causal tissues at *p < 0.05*. (B) Power to detect the causal tissues as a proportion of 1,000 simulations in which $h^2_{ge(t\prime)}$ > 0 for causal tissues at *p < 0.05*. (C) Bias on estimates of $h^2_{ge(t\prime)}$ for the causal tissue, while varying the correlation between the $\alpha_{gt}$ of each causal tissue. The dashed line indicates that the true value of $h^2_{ge(t\prime)}$ for either causal tissue. (D) Bias on estimates of $h^2_{ge(t\prime)}$ for non-causal tissues, while varying the correlation between the $\alpha_{gt}$ of each causal tissue. Error bars represent 95% confidence intervals, computed using the standard error of the mean. The value of $G_{t\prime}$ is set to the total number of unique *cis*-heritable genes across all tissues.

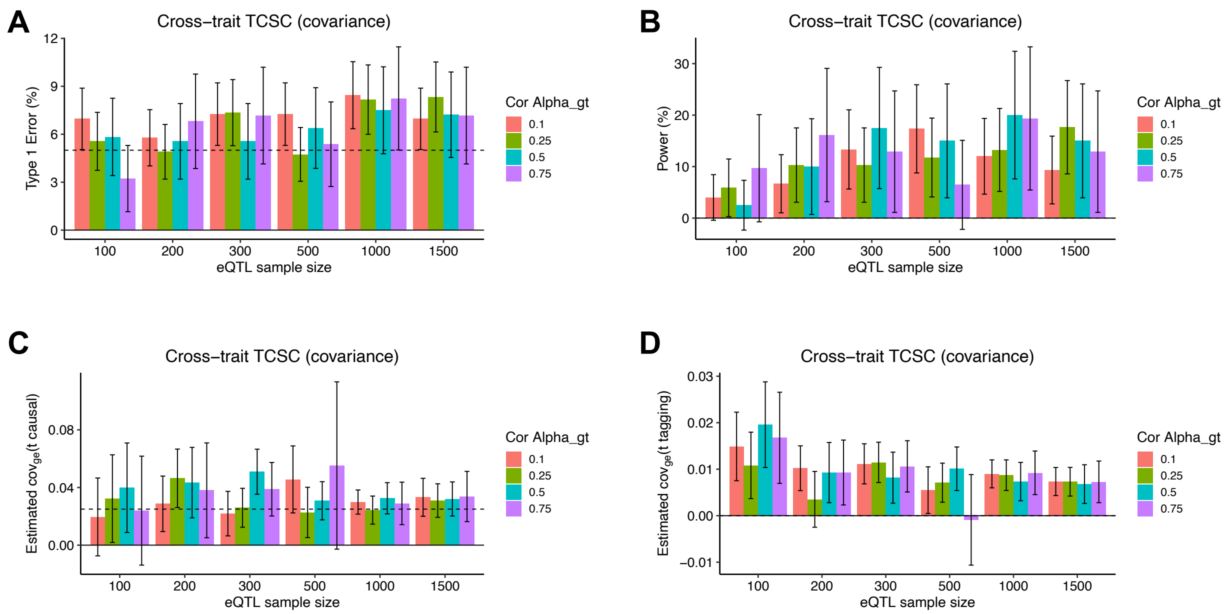


**Supplementary Figure 18. Robustness of cross-trait TCSC regression in simulations with two causal tissues with varying levels of correlated gene expression-trait effects.** To demonstrate how TCSC would behave when there are two causal tissues with correlation gene-trait effects ($\alpha_{gt}$), each tissue of which contributes 5% heritability to the trait. This is a model violation where TCSC assumes that gene expression-trait effects are i.i.d. (A) Type I error across 1,000 simulations with a varying correlation between the $\alpha_{gt}$ of each causal tissue. False positive event defined as $\omega_{ge(t')}$ > 0 for non-causal tissues at *p < 0.05*. (B) Power to detect the causal tissues as a proportion of 1,000 simulations in which $\omega_{ge(t')}$ > 0 for causal tissues at *p < 0.05*. (C) Bias on estimates of $\omega_{ge(t')}$ for the causal tissue, while varying the correlation between the $\alpha_{gt}$ of each causal tissue. The dashed line indicates that the true value of $\omega_{ge(t')}$ for either causal tissue. (D) Bias on estimates of $\omega_{ge(t')}$ for non-causal tissues, while varying the correlation between the $\alpha_{gt}$ of each causal tissue. Error bars represent 95% confidence intervals, computed using the standard error of the mean. The value of $G_{t'}$ is set to the total number of unique *cis*-heritable genes across all tissues.

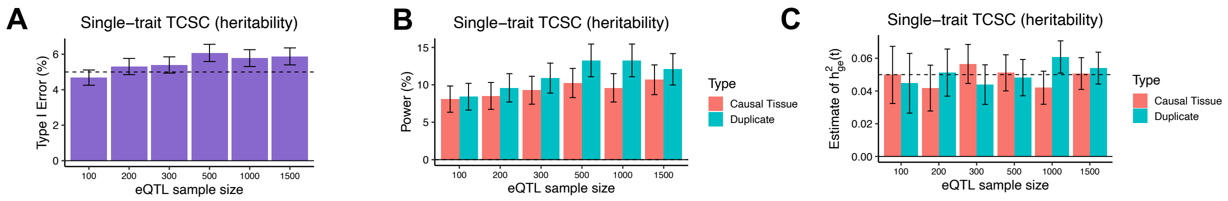


**Supplementary Figure 19. Robustness of single-trait TCSC regression in simulations with two causal tissues with identical gene expression-trait effects.** To demonstrate how TCSC would behave when there are two identical tissues contributing the same genetic component of gene expression to the trait. This is a model violation where TCSC assumes that gene expression-trait effects are i.i.d. To the original and duplicated causal tissue, we added a small amount of noise (with mean 0, variance 0.0025) to the tissue co-regulation scores of each duplicated tissue to avoid collinearity in the multiple linear regression. Across gene expression sample sizes, we find that TCSC estimates similar values of $h^2_{ge(t')}$ to each causal tissue, approximately one-half the value of the trait variance explained by the original causal tissue (0.1). Dashed line at 0.05, the expected tissue-specific contribution to heritability for both original and duplicated tissue.

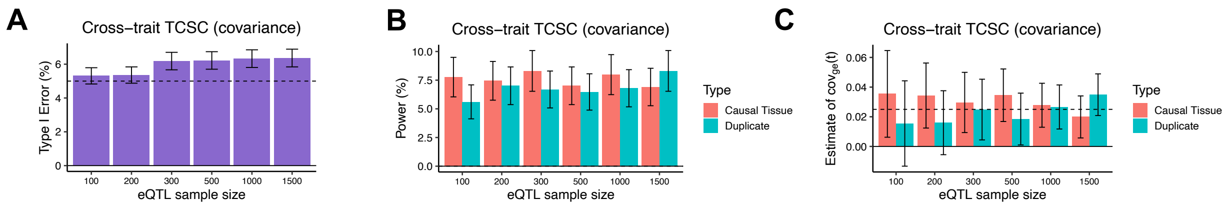


**Supplementary Figure 20. Robustness of cross-trait TCSC regression in simulations with two causal tissues with identical gene expression-trait effects.** To demonstrate how TCSC would behave when there are two identical tissues contributing the same genetic component of gene expression to the trait. This is a model violation where TCSC assumes that gene expression-trait effects are i.i.d. To the original and duplicated causal tissue, we added a small amount of noise (with mean 0, variance 0.0025) to the tissue co-regulation scores of each duplicated tissue to avoid collinearity in the multiple linear regression. Across gene expression sample sizes, we find that TCSC estimates similar values of $\omega_{ge(t')}$ to each causal tissue, approximately one-half the value of the trait variance explained by the original causal tissue (0.05). Dashed line at 0.025, the expected tissue-specific contribution to covariance for both original and duplicated tissue.

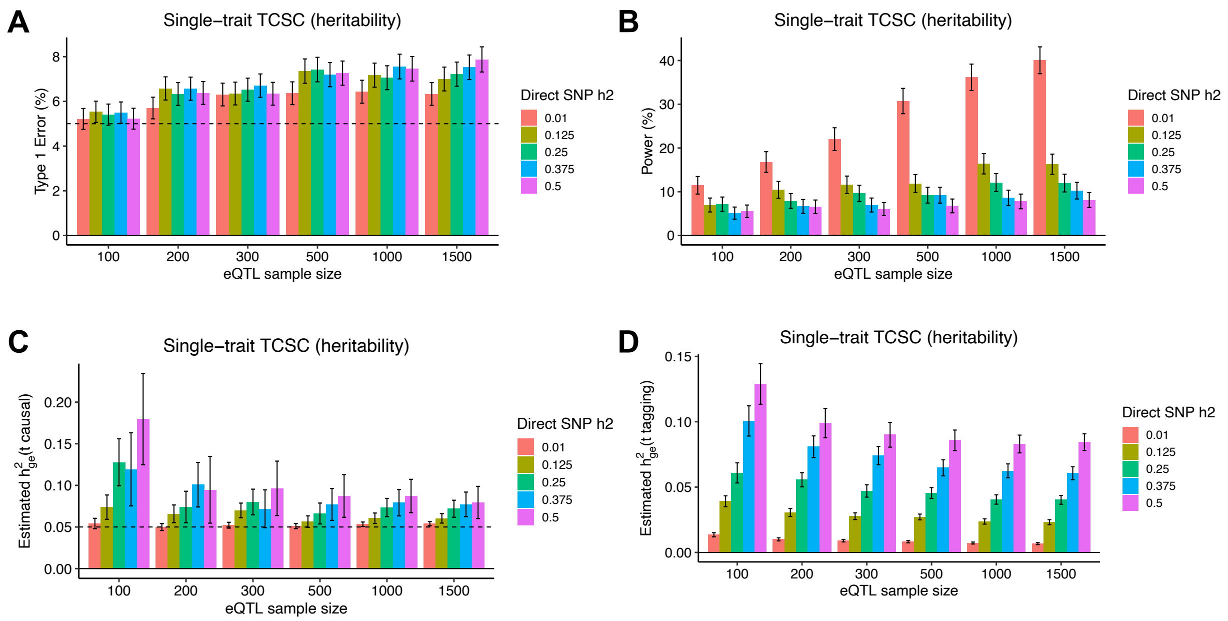


**Supplementary Figure 21**. **Robustness and power of TCSC regression in simulations with different amounts of direct SNP-trait heritability ($h^2_{SNP}$) not mediated by gene expression.** (A) Type I error across 1,000 simulations per value of $h^2_{SNP}$. False positive event is defined as $h^2_{ge(t\prime)} > 0$ for non-causal tissues at *p* < *0.05*. (B) Power to detect the causal tissue as a proportion of 1,000 simulations per value of $h^2_{SNP}$. A true positive event is defined as $h^2_{ge(t\prime)} > 0$ for causal tissues at *p* < *0.05*. (C) Bias on causal estimates of $h^2_{ge(t\prime)}$ for different values of $h^2_{SNP}$. Dashed line indicates true value of $h^2_{ge(t\prime)}$. (D) Bias on non-causal estimates of $h^2_{ge(t\prime)}$ for different values of $h^2_{SNP}$. Error bars represent 95% confidence intervals, computed using the standard error of the mean.

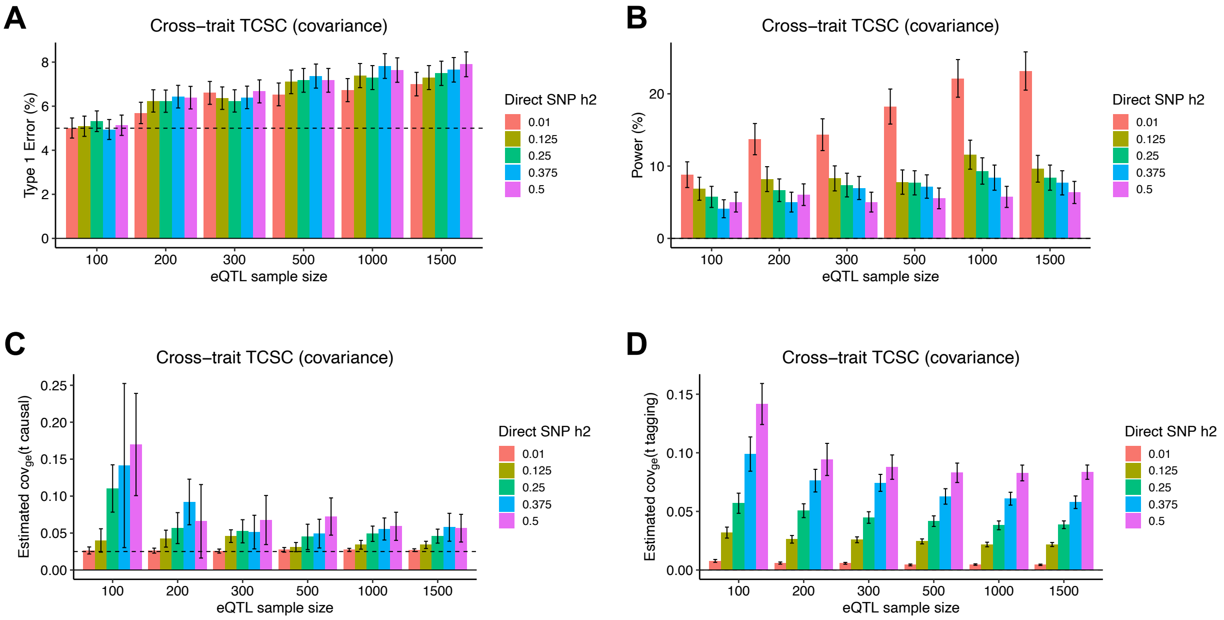


**Supplementary Figure 22. Robustness and power of TCSC regression in simulations with different amounts of direct SNP-trait heritability ($h^2_{SNP}$) not mediated by gene expression.** (A) Type I error across 1,000 simulations per value of $h^2_{SNP}$. False positive event is defined as $\omega_{ge(t\prime)} > 0$ for non-causal tissues at *p < 0.05*. (B) Power to detect the causal tissue as a proportion of 1,000 simulations per value of $h^2_{SNP}$. A true positive event is defined as $\omega_{ge(t\prime)} > 0$ for causal tissues at *p < 0.05*. (C) Bias on causal estimates of $h^2_{ge(t\prime)}$ for different values of $h^2_{SNP}$. Dashed line indicates true value of $\omega_{ge(t\prime)}$ . (D) Bias on non-causal estimates of $\omega_{ge(t\prime)}$ for different values of $h^2_{SNP}$. Error bars represent 95% confidence intervals, computed using the standard error of the mean.

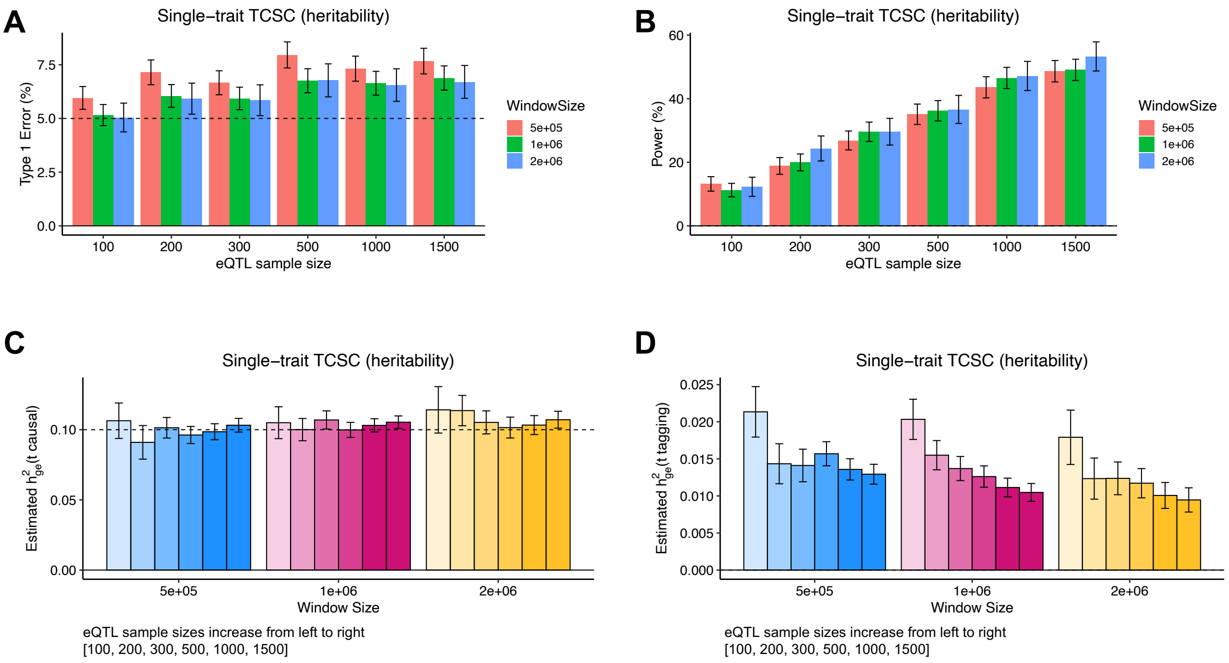


**Supplementary Figure 23. Robustness and power of TCSC regression in simulations with different values of window size used to calculate gene-gene co-regulation.** (A) Type I error across 1,000 simulations per window size. False positive event is defined as $h^2_{ge(t')}$ > 0 for non-causal tissues at *p < 0.05*. (B) Power to detect the causal tissue as a proportion of 1,000 simulations per window size. A true positive event is defined as $h^2_{ge(t')}$ > 0 for causal tissues at *p < 0.05*. (C) Bias on causal estimates of $h^2_{ge(t')}$ for different window sizes. Dashed lines indicate true values of $h^2_{ge(t')}$. (D) Bias on non-causal estimates of $h^2_{ge(t')}$ for different window sizes. Error bars represent 95% confidence intervals, computed using the standard error of the mean. The value of $G_{t'}$ is set to the total number of unique *cis*-heritable genes across all tissues.

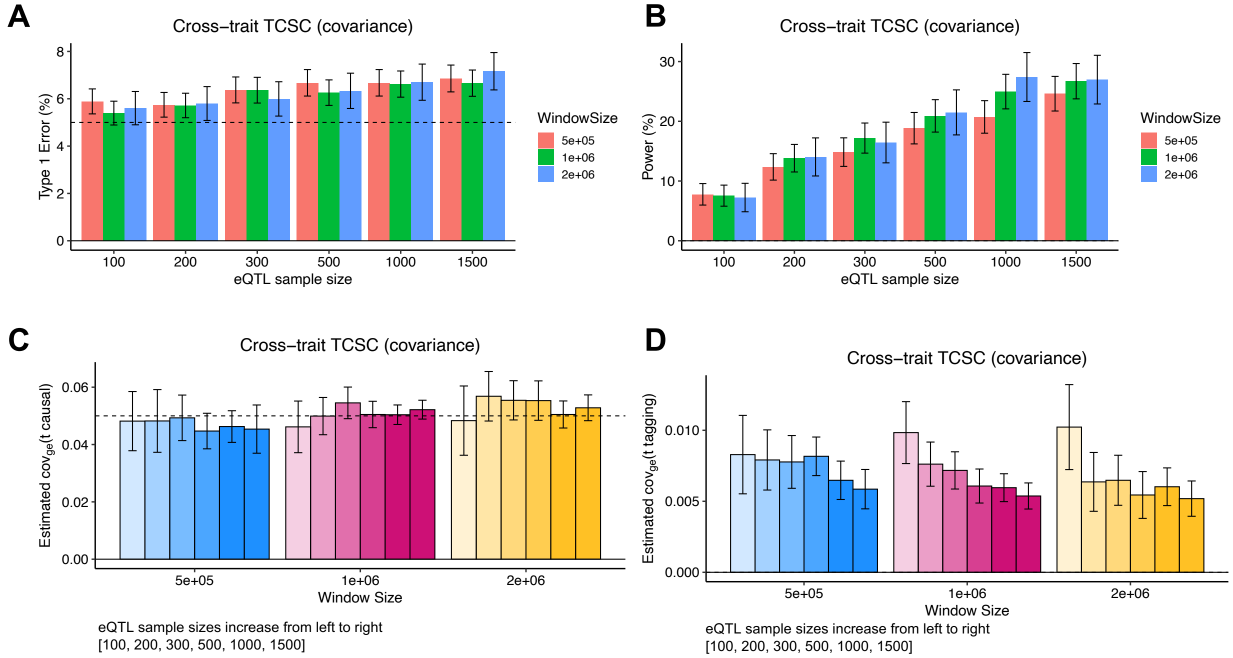


**Supplementary Figure 24. Robustness and power of cross-trait TCSC regression in simulations with different values of window size used to calculate gene-gene co-regulation.** (A) Type I error across 1,000 simulations per window size. False positive event is defined as $\omega_{ge(t')} > 0$ for non-causal tissues at *p < 0.05*. (B) Power to detect the causal tissue as a proportion of 1,000 simulations per window size. A true positive event is defined as $\omega_{ge(t')} > 0$ for causal tissues at *p < 0.05*. (C) Bias on causal estimates of $\omega_{ge(t')}$ for different window sizes. Dashed lines indicate true values of $\omega_{ge(t')}$. (D) Bias on non-causal estimates of $\omega_{ge(t')}$ for different window sizes. Error bars represent 95% confidence intervals, computed using the standard error of the mean. The value of $G_{t'}$ is set to the total number of unique *cis*-heritable genes across all tissues.

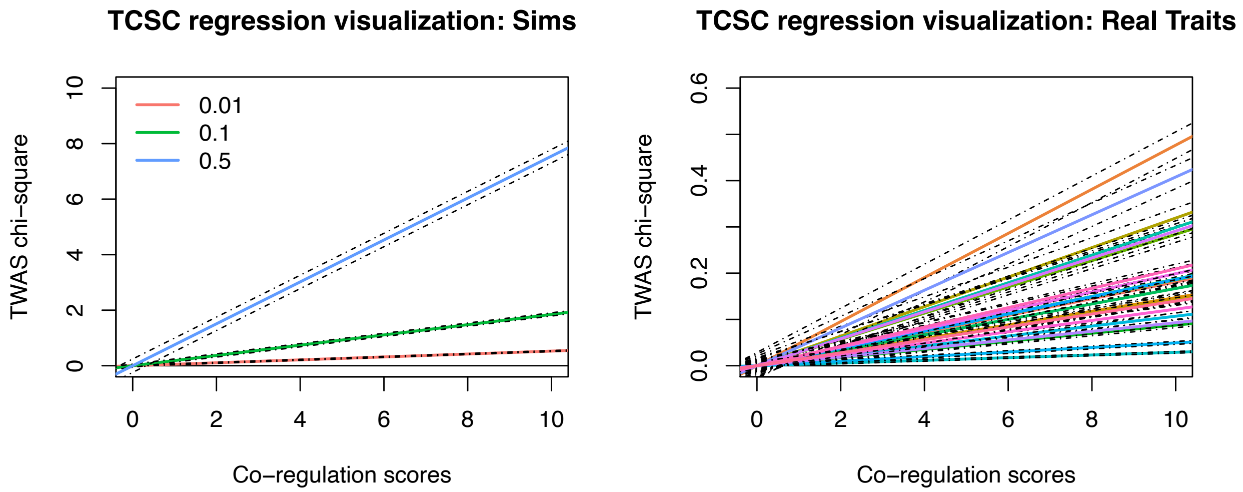


**Supplementary Figure 25. Visualization of TCSC regression.** (A) In simulations, we visualize the single-trait TCSC estimand ($h^2_{ge(t\prime)}$) as the line of best fit (slope) for one representative simulation from each of three true values of $h^2_{ge(t\prime)}$ using an intercept of 0. The dashed lines on either side of the lines of best fit represent the 95% confidence interval of the slope. All three slopes shown as significantly greater than zero. (B) In analysis of real traits, we visualize the single-trait TCSC estimand ($h^2_{ge(t\prime)}$) as the line of best fit (slope) for each of 21 significant tissue-trait pairs using an intercept of 0. The dashed lines on either side of the lines of best fit represent the 95% confidence interval of the slope. All three slopes shown are significantly greater than zero at 5% FDR.

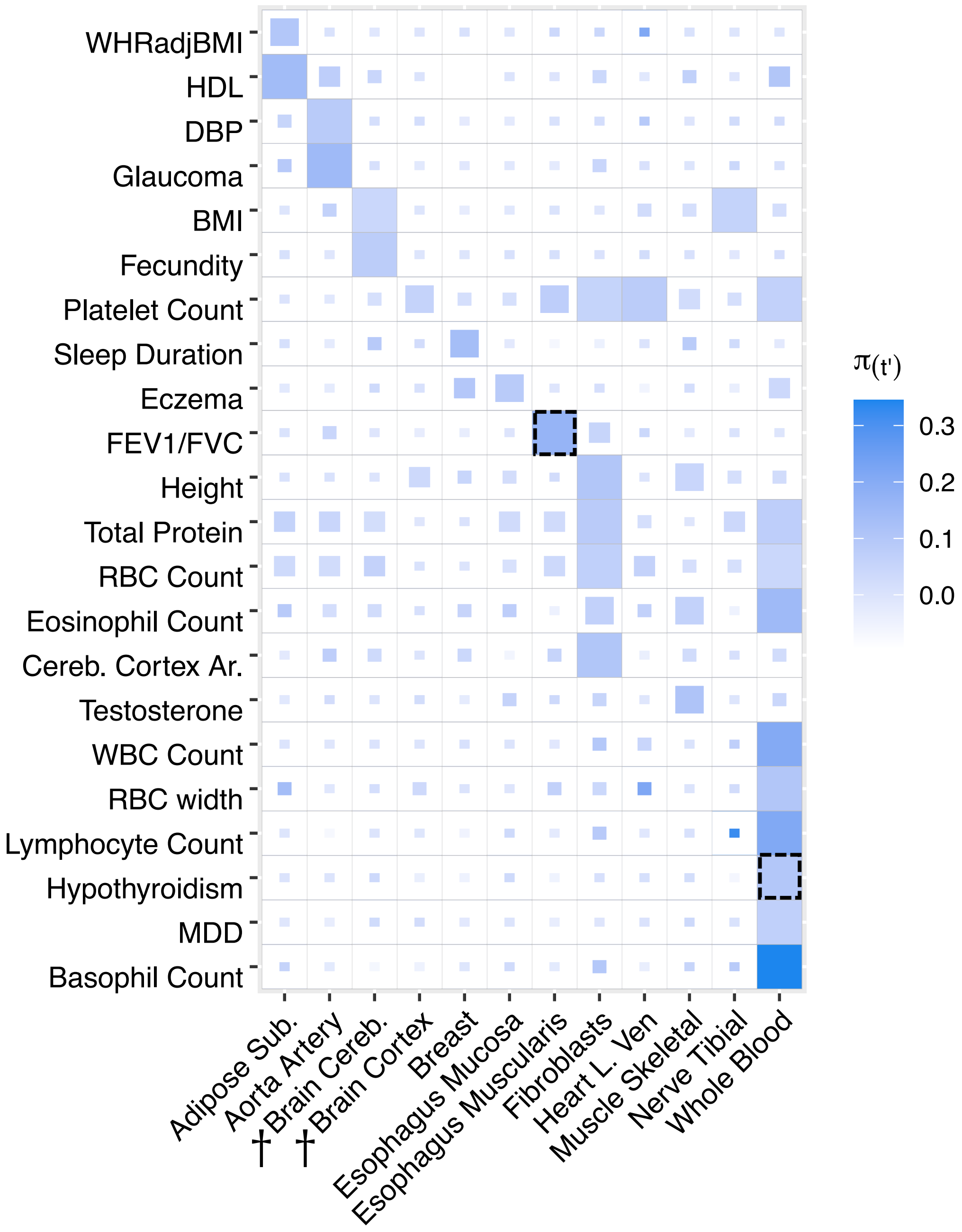


**Supplementary Figure 26. Tissue-specific contributions to disease and complex trait heritability in secondary analysis of 23 tissues, removing tissues with small eQTL sample size.** We report estimates of the proportion of disease heritability explained by the *cis*-genetic component of gene expression in tissue $t'$ ($\pi_{t'}$). Results are shown for tissue-trait pairs with FDR <= 10%, where full boxes indicate an FDR of 5%. Smaller box sizes are inversely proportional to the FDR. Tissues are ordered alphabetically. Color corresponds to $\pi_{t'}$ , the proportion of common variant heritability causally explained by predicted gene expression in tissue $t'$. These results are largely consistent with the analysis of 39 GTEx tissues (**Figure 4**). WHRadjBMI: waist-hip-ratio adjusted for body mass index. HDL: high-density lipoprotein. DBP: diastolic blood pressure. BMI: body mass index. FEV1/FVC: forced expiratory volume in one second divided by forced vital capacity. Cereb. Cortex Ar.: cerebral cortex surface area. WBC Count: white blood cell count. RBC Count: red blood cell count. MDD: major depressive disorder. Dashed lines indicate the result is discussed in detail in the main text. Daggers next to a tissue indicate a meta-tissue.

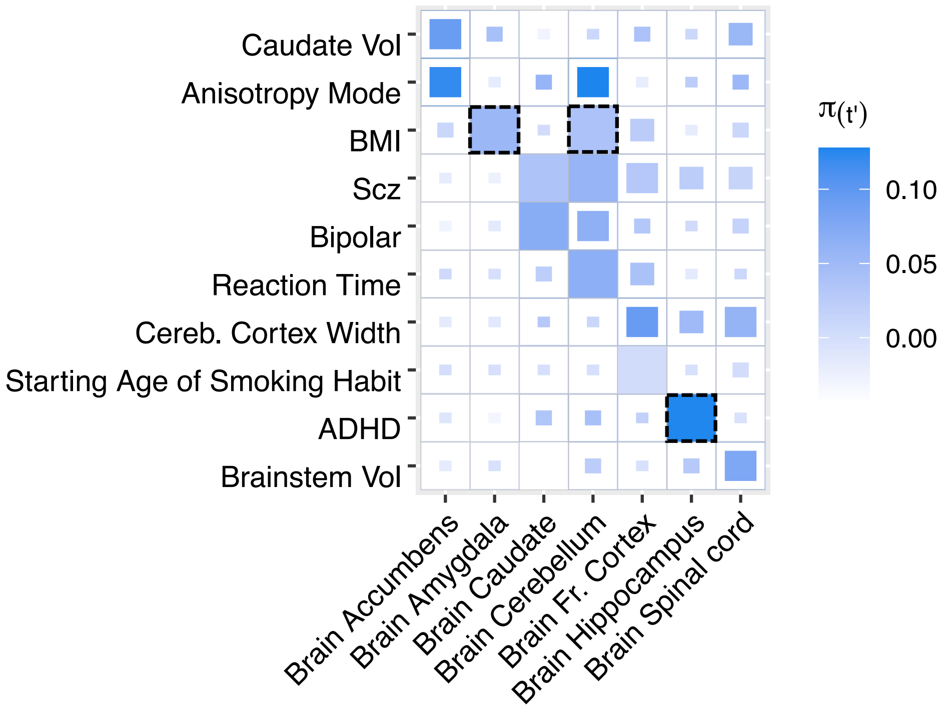


**Supplementary Figure 27. Tissue-specific contributions to disease and complex trait heritability in brain-specific analysis.** We separately analyzed results for 41 brain-related diseases and complex traits and 13 brain tissues. Results are shown for tissue-trait pairs with FDR <= 10%, where full boxes indicate an FDR of 5%. Smaller box sizes are inversely proportional to the FDR. Tissues are ordered alphabetically. Each tissue has an eQTL sample size ranging from 101 to 189 individuals. Color corresponds to $\pi_{t'}$ , the proportion of common variant heritability causally explained by predicted gene expression in tissue $t'$. Caudate Vol: caudate volume. BMI: body mass index.  Scz: schizophrenia. Bipolar: bipolar disorder. Brainstem Vol: brainstem volume. Cereb. Cortex Width: cerebral cortex width. ADHD: attention-deficit/hyperactivity disorder. Brainstem Vol: brainstem volume. Dashed lines indicate the result is discussed in detail in the main text.

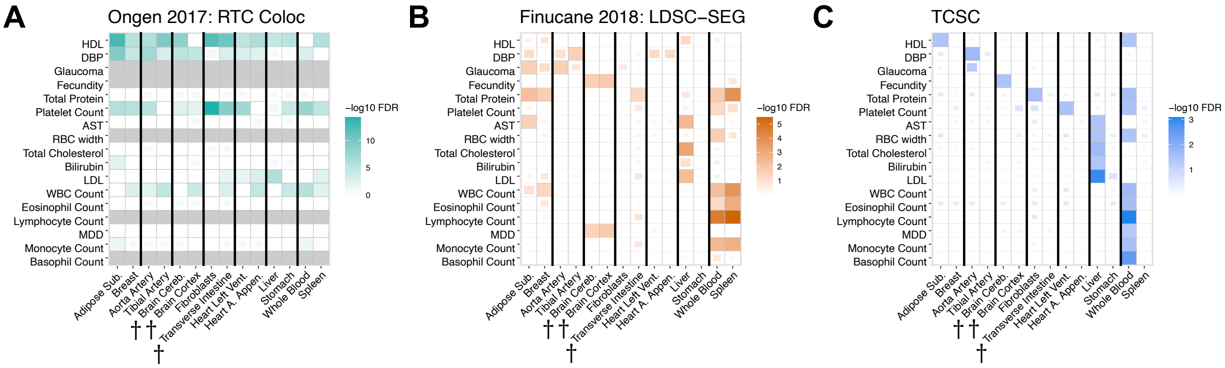


**Supplementary Figure 28. Comparison of disease-critical tissues identified by RTC Coloc, LDSC SEG and TCSC for all 17 disease/traits with causal tissue-trait associations identified by TCSC.** FDR significance of trait-tissue associations across three different methods for each of 17 traits with a significant tissue found by TCSC from **Figure 4** and 7 causal tissues, plus each tissue's most highly genetically correlated GTEx tissue. Full boxes indicate an FDR of 5%. Smaller box sizes are inversely proportional to the FDR. Thicker black lines separate the causal tissue found in the primary TCSC analysis (left) from its most highly genetically correlated GTEx tissue (right), with two exceptions. First, breast tissue was the most highly genetically correlated tissue for two causal tissues, adipose subcutaneous and thyroid; therefore, these three tissues appear as a trio.  Second, the aorta artery and tibial artery are each other's most highly genetically correlated tissue and both are a causal tissue different traits by TCSC. (A) RTC Coloc (Ongen 2017 Nat Genet), (B) LDSC SEG (Finucane 2018 Nat Genet), (C) TCSC. Per-trait FDR in panels A and C, FDR across traits and tissues in panel B. WHRadjBMI: waist-hip-ratio conditional on body mass index. HDL: high-density lipoprotein. DBP: diastolic blood pressure. BMI: body mass index. FEV1/FVC: forced expiratory volume in one second divided by forced vital capacity. Cereb. Cortex Ar.: cerebral cortex surface area. AST: aspartate aminotransferase. LDL: low-density lipoprotein. WBC: white blood cell count. MDD: major depressive disorder. Daggers next to a tissue indicate a meta-tissue. For BMI, fecundity, and cereb. cortex ar., LDSC SEG brain-specific analysis results are used.

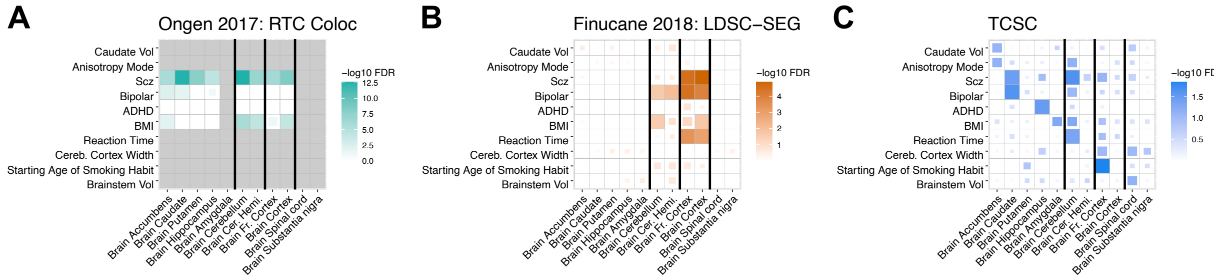


**Supplementary Figure 29. Comparison of disease-critical tissues identified by RTC Coloc, LDSC SEG and TCSC for brain-specific analysis.** Brain-specific tissue-trait pairs identified by three methods: RTC Coloc, LDSC SEG, and TCSC. Full boxes indicate an FDR of 5%. Smaller box sizes are inversely proportional to the FDR. Each tissue has an eQTL sample size ranging from 100 to 189 individuals. Additionally, we include each tissue's most highly genetically correlated GTEx tissue. Thicker black lines separate the causal tissue found in the primary TCSC analysis (left) from its most highly genetically correlated GTEx tissue (right), with one exception: brain accumbens, caudate, putamen, hippocampus, and amygdala are all highly genetically correlated, with no pairs of exclusively high genetic correlation. Caudate Vol: caudate volume. BMI: body mass index. Scz: schizophrenia. Bipolar: bipolar disorder. Brainstem Vol: brainstem volume. Cereb. Cortex Width: cerebral cortex width. ADHD: attention-deficit/hyperactivity disorder. Brainstem Vol: brainstem volume.
